# Detailed analysis of public RNAseq data and long non-coding RNA: a proposed enhancement to mesenchymal stem cell characterisation

**DOI:** 10.1101/2020.03.09.976001

**Authors:** Sebastien Riquier, Marc Mathieu, Anthony Boureux, Florence Ruffle, Jean-Marc Lemaitre, Farida Djouad, Nicolas Gilbert, Therese Commes

## Abstract

The development of RNA sequencing (RNAseq) and corresponding emergence of public datasets have created new avenues of transcriptional marker search. The long non-coding RNAs (lncRNAs) constitute an emerging class of transcripts with a potential for high tissue specificity and function. Using a dedicated bioinformatics pipeline, we propose to construct a cell-specific catalogue of unannotated lncRNAs and to identify the strongest cell markers. This pipeline uses *ab initio* transcript identification, pseudoalignment and new methodologies such as a specific k-mer approach for naive quantification of expression in numerous RNAseq data.

For an application model, we focused on Mesenchymal Stem Cells (MSCs), a type of adult multipotent stem-cells of diverse tissue origins. Frequently used in clinics, these cells lack extensive characterisation. Our pipeline was able to highlight different lncRNAs with high specificity for MSCs. *In silico* methodologies for functional prediction demonstrated that each candidate represents one specific state of MSCs biology. Together, these results suggest an approach that can be employed to harness lncRNA as cell marker, showing different candidates as potential actors in MSCs biology, while suggesting promising directions for future experimental investigations.

## 1 Introduction

The increasing popularity of RNAseq and the ensuing aggregation of this data-type into public databases enable the search for new biomarkers across large cohorts of donors or cell types for the purpose pathological conditions or cellular lineages identification. As such, RNAseq has paved the way for the discovery of novel transcriptional biomarkers such as long noncoding RNAs (lncRNAs), that have emerged as a fundamental molecular class. A growing number of lncRNAs have been identified in the last decades, with their number approaching that of coding RNAs (17910 annotated human lncRNAs in the latest v32 version of the GENCODE versus 19 965 coding genes).

An increasing body of evidence has highlighted characteristics that define lncRNAs as therapeutic targets as well as potential tissue-specific markers [1]. Indeed, despite their non-coding nature, a large spectrum of functional mechanisms have been associated to lncRNAs [2, 3]. These include: endogenous competition (miRNA sponging for example), protein complex scaffolding and guides for active proteins with RNA-DNA homology interactions. These mechanisms occur in various physiological or pathological processes such as development, cancer and immunity [4, 5, 6]. To date, there is no finite list of long non-coding isoforms, making it difficult to construct a complete lncRNA catalogue due to the high number of transcripts and their tissue-specific expression [7, 8]. The absence of a complete catalogue makes it difficult to establish a comprehensive lncRNA expression profile.Thus, currently, the best strategy for the study of lncRNAs consists in the prediction of transcripts from a selection of RNAseq data in a tissue-specific condition. This strategy was successful in novel lncRNA biomarker discovery in pathological conditions [9, 10], but was poorly explored for cell lineage characterisation. Taking into account their functional importance and specificity, these RNAs should therefore not be ignored in establishing the molecular identity of a cell type.

Cell characterisation by specific markers bring different challenges such as the importance of probing the specificity of the marker and its limits in an extended number of various cell types, rather than using a control/patient experimental model. Moreover, the cells are not in a fixed state and display a variable transcriptional activity depending on cell status, environment, culture conditions and other parameters [1]. Furthermore, the lncRNAs’ function is generally poorly assessed, except in the case of recurrent known transcripts (HOTAIR, H19). Thus, the *in silico* elaboration of a lncRNA catalogue that document the functional domains where the candidates could act, will be beneficial in the identification of lncRNAs’ role and thus, in future experiments. To this end, we have developed an integrated four-step procedure consisting of: i) an *ab initio* transcript reconstruction from RNAseq data and characterisation of novel transcripts, ii) a differential analysis using pseudoalignment coupled with a machine learning solution in order to extract the most cell-specific candidates, iii) an original step of tissue-expression validation with specific k-mers search in large and diversified transcriptomic datasets iv) an in-depth analysis to predict lncRNAs’ functional potential from *in silico* prediction approaches. The notable advantage of introducing an *in silico* verification using k-mers is to allow a precise and in-depth determination of lncRNAs expression profile and to quickly interrogate their lineage specificity. In addition to that, validation of newly identified lncRNAs has been undertaken using RT-qPCR and long read sequencing (with Oxford Nanopore technology).

Mesenchymal Stem Cells (MSCs) are defined as multipotent adult stem cells, harvested from various tissues, including Bone Marrow (BM), Umbilical Cord (UC), and Adipose tissue (Ad). MSCs are an interesting cell type to explore since these cells lack the extended transcriptional characterisation that could highlight their lineage belonging and/or the possibility of distinguishing them from other mesodermal cell types such as fibroblasts and pericytes [11, 12]. The commonly admitted surface markers for MSCs, proposed by the International Society for Cellular Therapy (ISCT), and required to identify MSCs since 2006 are THY1 (CD90), NT5E (CD73), ENG (CD105) concerning the positive markers, and CD45, CD34, CD14 or CD11b, CD79alpha or CD19 and HLA-DR concerning the negative markers [13]. These markers are not distinctive and may therefore not be sufficient for the definition of cellular or biological properties. Considering their different therapeutic properties (chondro and osteo differentiation potential, immunomodulation and production of trophic factors) [14] and given the increasing usage of these cells for academic and preclinical research [15], a detailed molecular characterisation of MSCs and predictive marker of functionality will constitute an important tool in regenerative medicine. lncRNAs have emerged as a class of transcripts with tissue-specific expression and important functions, such as the regulation of MSCs function [16, 17, 18], and remain largely unexplored in these cells.

To address this need, we performed a broad transcriptomic analysis of novel lncRNAs on human Mesenchymal Stem Cells (MSCs). We started from publicly available MSCs RNAseq, selecting ribodepleted datasets in order to enhance lncRNAs discovery and to explore the poly-(A)+ as poly(A)-lncRNAs. We restricted the differential expression analysis to a bone-marrow MSCs source compared to “nonMSC” cell counterparts. Once achieved, in depth *in silico* analysis was performed to check the lncRNA cell specific profiles with more and extensive datasets. To validate our approach, RNAseq data from eight publicly available libraries of normal MSCs containing a large diversity of noncancerous cell types were used for novel lncRNAs detection and tissue expression comparison. We initially reconstructed more than 70000 unannotated lncRNAs present in human bone-marrow MSCs. These lncRNAs were assigned depending on their position relative to annotated genes: MSC-related Long Intergenic Non Coding trancripts” named “Mlinc”, and MSC-related Long Overlapping Antisense transcripts called “Mloanc”. Among them, 35 Mlincs were specifically enriched in the cell lineage compared to the “non-MSC” counterpart group. Finally, after a further selection of the three most specific Mlincs, detailed *in vitro* and *in silico* functional exploration were performed.

## 2 Material and methods

### 2.1 Data collection and basic processing

The public RNAseq datasets (in fastq format) have been assessed using the ENCODE, EBI ArrayExpress service or SRA database at each step of the pipeline: i) lncRNAs prediction and first differential analysis (Table S1), ii) k-mer search in ENCODE data to refine lncRNAs’ specificity (Table S2), iii) k-mer search in FANTOM6 CAGE dataset and single cell analysis (scRNAseq) from Adipose MSCs by X. Liu et al raw data [19] for functional investigations (Table S3), iv) k-mer search in MSCs in different conditions (Table S4).

The reads quality were assessed with FastQC (https://www.bioinformatics.babraham.ac.uk/projects/fastqc/) to avoid the implementation of poor quality data in the analysis. Data from Peffers et al. [20], added to ENCODEs BM-MSCs RNAseq data, were selected for the MSClinc and Mloanc characterisation and the differential step analysis considering the above-mentioned features: Ribo-zero technology, stranded and paired-ends RNAseq. Peffers data had a forward-reverse library orientation instead of a reverse-forward orientation of a classic Illumina sequencing, thereby the order of paired files was manually reversed. The fastq files used for lncRNAs prediction, referred as “MSC” group, were used for the differential analysis against the other cell types as “non-MSC” group, (Table S1) were mapped using CRAC v2.5.0 software [21] on the indexed GRCh38 human genome, including mitochondria, with stranded, -k 22 and rf options.

### 2.2 *Ab initio* assembly for transcripts prediction or unannotated transcripts prediction

The aligned reads of the “MSC” group were put through *ab initio* transcript assembly. Unannotated transcripts were predicted with the following procedure: i/an *ab initio* reconstruction was performed on individual RNAseq with the StringTie [22] version 1.3.3b, with -c 5 -j 5 rf -f 0.1 (5 spliced reads are necessary to predict a junction and 5 reads at minimum are required to predict an expressed locus), ii/ the output individual gtf files obtained with the RNAseq of “MSC” group were then merged with the StringTie version 1.3.3b with -f 0.01 -m 200 and with a minimum TPM of 0.5, with the Ensembl human annotation (hg38) v90 used as guide for StringTie. The GTF was parsed with BEDTools [23] to dissociate new intergenic lncRNAs (lincRNAs) from annotated RNAs (coding or annotated lncRNAs), by applying filter criteria classically used in lncRNAs prediction [24], excluding transcript models overlapping (by 1 bp or more) any annotated coordinates. The resulting GTF of unannotated lincRNA from MSCs is referred as “Mlinc”.

In parallel, the GTF was parsed with BEDTools to dissociate overlapping-antisens lncRNAs (named Mloanc), by applying filter criteria classically used in lncRNAs prediction, keeping transcript overlapping any annotated coordinates, then excluding transcript models overlapping these annotated coordinates on the same strand. The resulting GTF of MSCs overlapping-antisens lncRNAs is referred as “Mloanc” (Figure 1).

**Figure 1:**
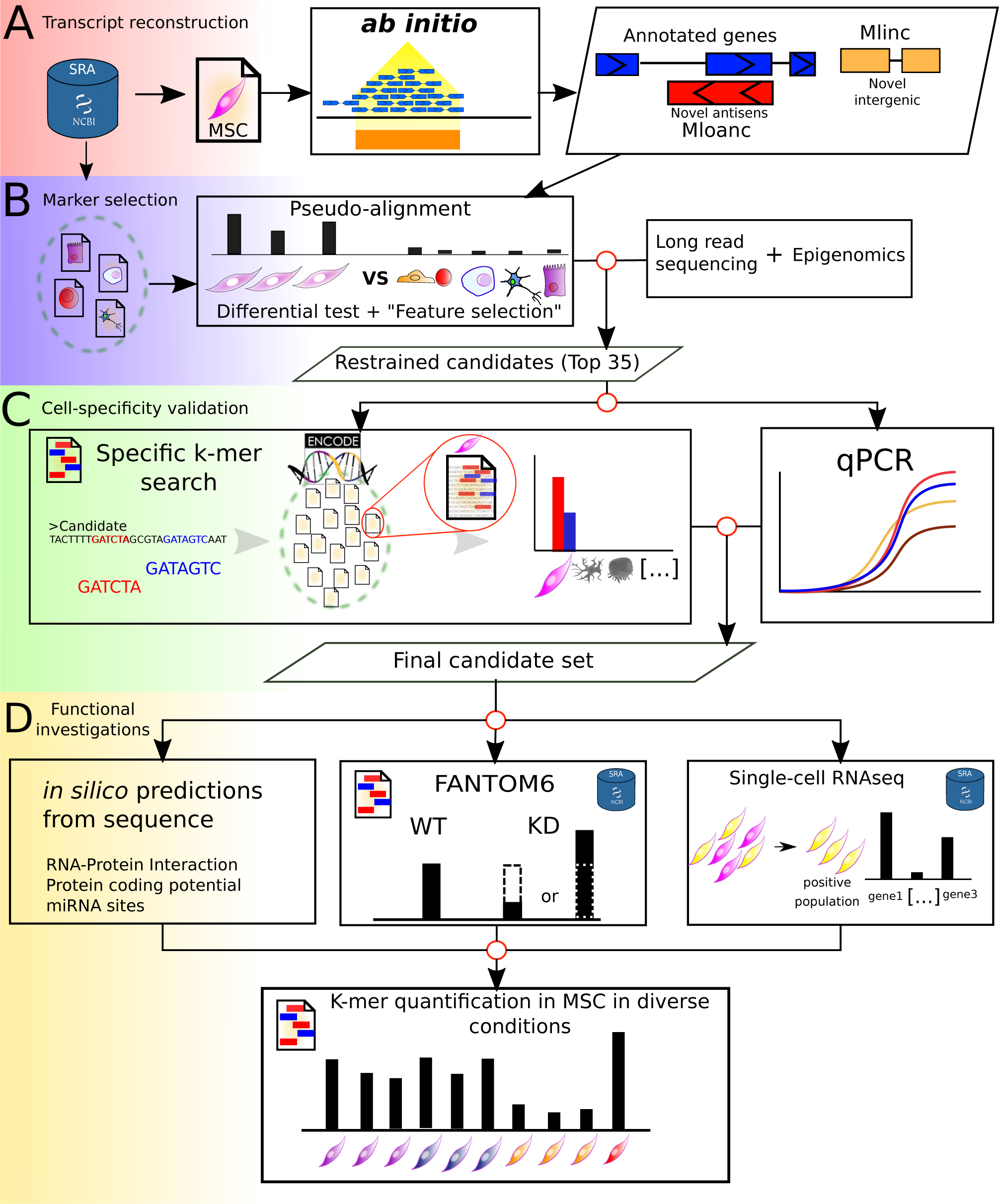
Flowchart representation of the Pipeline used for this in the study. The 4 steps of the flowchart are described A) Ab initio reconstruction of transcrit expressed in MSC from SRA dataset and creation of a reference (gtf+fasta) for quantification of Ensembl annotated genes, unannotated intergenic (Mlincs) and unannotated overlapping antisens (Mloanc). The results are shown in Fig2. B) Differential Analysis for the selection of MSC markers (restrained candidate set) with i/ kallisto pseudoalignement and Sleuth differential test followed by feature selection by random forest with Boruta package. Long read sequencing and active transcription in MSC by epigenetic marks information completed the selection step (see figures 2 and 3). C) Validation of cell expression specificity of candidates by kmer quantification in ENCODE RNAseq datasets (see Table S2 for list of data) and qPCR validation. The results are presented in figure 4. D) Functional investigations were performed with in silico prediction methods from the sequence of candidates, followed by k-mer quantification with FANTOM6 data set, single cell RNAseq and selected MSC conditions. Kmer quantification phases are shown by corresponding icons (figures 5 and 6).

### 2.3 Long-read sequencing

The library was generated with 250 ng polyA+ mRNA purified from 50 *µ*g of human BM-MSCs total RNA. The polyA+ mRNA were treated according to the cDNA-PCR sequencing kit protocol (ref SQK-PCS108) as recommended by Oxford Nanopore. 3 254 396 sequences were obtained on the Oxford Nanopore Minion sequencer. The base calling was done with albacore version 2.2.7. 2 720 928 long-reads were successfully mapped using Minimap2 [25] version 2.10-r764 on GRCh38 human genome with default options used for Oxford Nanopore technology.

### 2.4 Quantification with pseudoalignment and feature selection

Kallisto v0.43.1 [26] was used directly on RNAseq raw fastq from the “MSC” and “non-MSC” groups. This pseudoalignment was performed with a number of bootstraps (-b) of 100, using a Kallisto index containing the sequences of all transcripts: the Ensembl coding and non-coding transcripts (v90) plus the predicted lincRNA and lncoaRNAs. Sleuth version 0.29.0 [27] was used with R for differential expression statistical analysis using the Walt test method, to compare the “MSC” group against the “non-MSC” group (including Lymphocytes, Macrophages, Hepatocytes, IPS, ESCs, HUVECs, Neurons, chondrocytes). Analysis was performed at the gene level for the annotated genes and at the transcript level for the predicted lincRNA and lncoaRNAs. Genes or lncRNAs having a log2 fold-change between “MSC” and others greater than 0.5 and a p-value lower than or equal to 0.05 were selected. Finally, only transcripts/genes overexpressed in MSCs were selected. Each category (annotated transcripts, lincRNAs and lncoaRNAs) of potential candidates passing the first differentiation expression filter were separated for feature selection analysis. Boruta 6.0 [28] was used with 10000 maximum runs and a pvalue of 0.01 on each category, with multiple comparisons adjustment using the Bonferroni method (mcAdj = TRUE). Candidates passing the boruta test as “Confirmed” for each category were selected as reliable biomarkers.

### 2.5 Quantification by k-mers search

To quantify the expression of a transcript or a gene in available RNAseq with a rapid procedure, specific k-mers of 31nt length were extracted from the candidate sequence. A specific k-mer of an annotated candidate corresponds to a 31nt sequence that maps once on the genome and once on the reference transcriptome (Ensembl v90). In case of unannotated transcript (Mlinc, Mloanc) a specific k-mer maps once on the genome and is absent from reference transcriptome. The automated selection of specific k-mers is ensured by the Kmerator tool (in preparation) (https://github.com/Transipedia/kmerator). The k-mers were then quantified directly in raw fastq files using countTags (https://github.com/Transipedia/countTags). The quantification is expressed by the average count of all k-mers for one transcript, normalised by million of total k-mers in the raw file.

In FANTOM6 Dataset (https://doi.org/10.1101/700864, article not peer-reviewed) containing CAGE Analysis, to approach a “Transcript Per Millions” normalisation, the number of k-mers quantified was normalised by the total number of reads in million.

### 2.6 Genomic intervals assessment

DNase-seq intervals of enrichment were directly downloaded from ENCODE in bed format for BM-mesenchymal cells (ENCFF832FHZ) and hematopoietic progenitors (ENCFF378FCS). The H3K27ac (GSM3564514) and H3K4me3 (GSM3564510) ChIP results from undifferentiated BM-MSCs of the Agrawal Singh S. et al. study [29] were downloaded from GEO database in wig format, and remapped to the hg38 genome with CrossMap (http://crossmap.sourceforge.net/).

### 2.7 *in silico* functional prediction

We used LncADeep [30] to identify particular correlations between candidates and proteins. Beginning with our selection of 3 candidates, we filtered shared predicted proteins and selected protein uniquely predicted as interacting uniquely with the concerned candidate. The pathways concerned with these unique protein were identified with reactome.

Tarpmir was used to identify possible target site of human miRNA from miRbase (p = 0.5) [31] and FEELnc [32] to decipher the coding potential of candidates, using the coding and non-coding par of Ensembl annotation sequences as model.

### 2.8 Single-cell analysis

Single-cell data were pseudoaligned with Kallisto, with the same index used for the initial bulk RNAseq analysis. Pseudoalignment of 10X genomics data, correction, sorting and counting was made by Kallisto “bus” functions. Count matrices were processed with Seurat R package [33, 34]. Empty droplets were estimated by barcode ranking knee and inflection points, only droplet with a minimal count of 10000 were kept. In the end, 26071 droplets remain.

After normalisation, Inter-donor batch effect was corrected with ComBat method in sva R package [35] (Combat function, prior.plots=FALSE, par.prior=TRUE). Cell cycle scoring was made by CellCycleScoring Seurat function, using gene set used by the initial authors [19]. Finally, other sources of unnecessary variability as percent of mitochondrial genes, cell cycle and number of UMIs were regressed using ScaleData Seurat function.

To decipher genes enriched in cells positive for our markers, cells with a scaled expression superior or equal to 0.1 were labelled as positives, whereas cells with an expression inferior to the level were labelled as negatives. Then, markers of these cell were deciphered using FindAllMarkers Seurat function with a minimum FC threshold of 0.15. Expression of our markers in the Ad-MSCs population was made by FeaturePlot Seurat function after UMAP dimensional reduction, the gene enrichments were represented with VlnPlot function.

### 2.9 Data visualisation

Genome browser-like figures were generated with Gviz R package [36]. Bam tracks were generated from merged BAMs used for transcript prediction. Heatmaps were generated using superHeat R package (https://github.com/rlbarter/superheat).

### 2.10 Ethics approval and consents

Human primary MSCs was obtained from patients who had granted the authors written informed consent with approval of the General Direction for Research and Innovation, the department in responsible for questions of ethics within the French Ministry of Higher Education and Research (registration number: DC-2009-1052). Human primary myoblasts were collected from patients of the CHU of Montpellier, France (the Montpellier University Hospital) who had provided informed consent. All experiments were performed in accordance with the Declaration of Helsinki and approved by the ethical committee of the CHU of Montpellier (France). Samples were approved for storage by the French “Ministre de l’Enseignement et de la Recherche” (NDC-2008-594). Liver samples were obtained from the Biological Resource Center of Montpellier CHU (CRB-CHUM; http://www.chu-montpellier.fr; Biobank ID: BB-0033-00031). The procedure was approved by the French Ethics Committee and written or oral consent was obtained from the patients.

### 2.11 Cell preparation and culture conditions

MSCs were isolated from bone marrow aspirates of patients undergoing hip replacement surgery, as previously described [37]. Cell suspensions were plated in *α*-MEM supplemented with 10 % FCS, 1 ng/mL FGF2 (R&D Systems), 2 mM L-glutamine, 100 U/mL penicillin and 100 *µ*g/mL streptomycin. These were shown to be positive for CD44, CD73, CD90 and CD105 and negative for CD14, CD34 and CD45 and used at the third or fourth passage. Human skin fibroblasts were cultured in DMEM high glucose supplemented with 10 % FCS. For Ad-MSCs isolation, adipose tissue was digested with 250 U/mL collagenase type II for 1 h at 37 *^◦^*C and centrifuged (300 g for 10 min) using routine laboratory practices. The stroma vascular fraction was collected and cells filtered successively through a 100 *µ*m, 70 *µ*m and 40 *µ*m porous membrane (Cell Strainer, BD-Biosciences, Le-Pont-de-Claix, France). Single cells were seeded at the initial density of 4000 cell/cm2 in *α*MEM supplemented with 100 U/mL penicillin/streptomycin (PS), 2 mmol/mL glutamine (Glu) and 10% fetal calf serum (FCS). After 24 h, cultures were washed twice with PBS. After 1 week, cells were trypsinised and expanded at 2000 cells/cm2 till day 14 (end of passage 1), where Ad-MSCs preparations were used.

Human umbilical vein endothelial cells (HUVEC) obtained from Clonetics (Lonza, Levallois Perret, France) were cultured in complete EGM-2MV (Lonza) supplemented with 3 % FCS (HyClone; Perbio Science, Brebires, France) Primary human myoblasts were isolated and purified from skeletal muscles of donors, as described by Kitzmann et al [38]. Purified myoblasts were plated in Petri dishes and cultured in growth medium containing Dulbecco’s Modified Eagle’s Medium (Gibco) supplemented with 20 % foetal bovine serum (FBS) (GE Healthcare, PAA), 0.5 % Ultroser G serum substitute (PALL life sciences) and 50 *µ*g/ml Gentamicin (Thermo Scientific, France) at 37°C in humidified atmosphere with 5 % CO_2_. All experiments were carried out between passage 4 (P4) and P8 to avoid cell senescence.

IPSCs were maintained in mTeSR-1TM medium (STEMCELL Technologies), in Petri dishes with matrigel (Corning, France). For the passages, cells were incubated in Gentle Cell Dissociation Reagent (STEMCELL Technologies) at room temperature, dissociation medium was discarded and cells incubated in mTeSR medium. All cell cultures were performed at 37*^◦^*C with 5% of O_2_ and 10% of CO_2_.

Primary human hepatocytes (PHHs) were isolated, as described previously [39], from liver resections performed in adult patients. NSC derived from H9 or directly bought (StemPro) have been cultivated on laminine with StemPro NSC SFM medium. H9 embryonic stem cells were cultivated in ESICO medium in a coculture H9/MEF (Mouse Embryonic Fibroblasts) at 37 *^◦^*C with 5% of O_2_ and 5% CO_2_.

### 2.12 RNA preparation and reverse transcription

Total RNA was isolated using TRIzol reagent (invitrogen) or RNeasy Mini Kit (Qiagen, France) according to the manufacturer protocol. RNA was quantified using a NanoDrop ND-1000 spectrophotometer (Thermo Fisher Scientific, France). RNA quality and quantity were further assessed using the 2100-Bioanalyzer (Agilent Technologies, Waldronn, Germany). Only preparations with RNA integrity number (RIN) values above 7 were considered. Reverse-transcription was performed either with random hexamers using the GeneAmp Gold RNA PCR Core kit (Applied Biosystems) or with oligo(dT) using SuperScriptTM First-Strand Synthesis System for RT-qPCR (invitrogen, France).

### 2.13 Real-time quantitative PCR

Primer pairs were designed with primer3 online software (http://bioinfo.ut.ee/primer3-0.4.0/) from the transcripts’ sequences. Primer pairs with a perfect and unique match on the human genome were validated with ucsc blat software (https://genome.ucsc.edu). As a final verification, primers were visualised in parallel with the bam alignment using IGV (http://software.broadinstitute.org/software/igv/) to verify that the primers overlap zones with read coverage. If possible, primer-pairs were designed to span an intron when present in the genomic sequence. Primers were designed for a mean Tm of 60*^◦^*C. Quantitative PCR (qPCR) were performed using LightCycler 480 SYBR Green I Master mix and real-time PCR instrument (Roche). PCR conditions were 95 *^◦^*C for 5 min followed by 45 cycles of 15 s at 95 *^◦^*C, 10 s at 60 *^◦^*C and 20 s at 72 *^◦^*C. For each reaction, a single amplicon with the expected melting temperature was obtained.

The gene encoding ribosomal protein S9 (RPS9) was used as house-keeping gene for normalisation. The threshold cycle (Ct) of each amplification curve was calculated by Roche’s LightCycler 480 software using the second derivative maximum method. The relative amount of transcripts were calculated using the ddCt method [40].

## Results

For the purpose of generating a catalogue of all transcripts in any particular cell type, we developed a pipeline for the characterisation of all RNAs and their expression profile in a large collection of RNAseq data. The procedure includes four steps: i) an *ab initio* transcripts reconstruction from RNAseq data and identification of unannotated transcripts, ii) a differential analysis using pseudoalignment coupled with a machine learning solution in order to extract the most cell-specific candidates, iii) an original step of tissue-expression validation with a kmer approach (comparing large transcriptomic datasets), iv) an in-depth analysis to predict lncRNAs functional potential from *in silico* prediction approaches (Figure 1). To illustrate the procedure, we produced a RNA catalogue from bone-marrow MSCs (”MSC” group).

### 2.14 General features of the predicted MSC catalogue of lncRNAs

As mentioned above, we started with the *ab initio* reconstruction of any transcript from bone-marrow RNAseq with Stringtie assembler after mapping the reads with the CRAC software (see Materials and Methods for parameters). New isoforms of annotated transcripts were ignored. Of the 200 243 transcripts present in Ensembl annotation (version 90), 105 511 (52.6%) were detected in MSCs (TPM *>*0.1 in pseudoalignment quantification).

73 463 new lncRNAs were reconstructed. This fraction of unannotated transcripts represent 41% of detected transcripts, so in our case the *ab initio* reconstruction made it possible to almost double the inventory of detectable signatures in MSCs (Figure 2A). Of these, 34 712 were found to be intergenic and were thus referred to as “Mlinc” RNAs (MSC-related long intergenic non-coding RNAs), and 38 751 were found to be overlapping coding regions but in anti-sense orientation and thus referred to as “Mloanc” RNAs (MSC-related long overlapping antisense non-coding RNAs, with criteria as described in materials and methods and Figure 2A).

**Figure 2:**
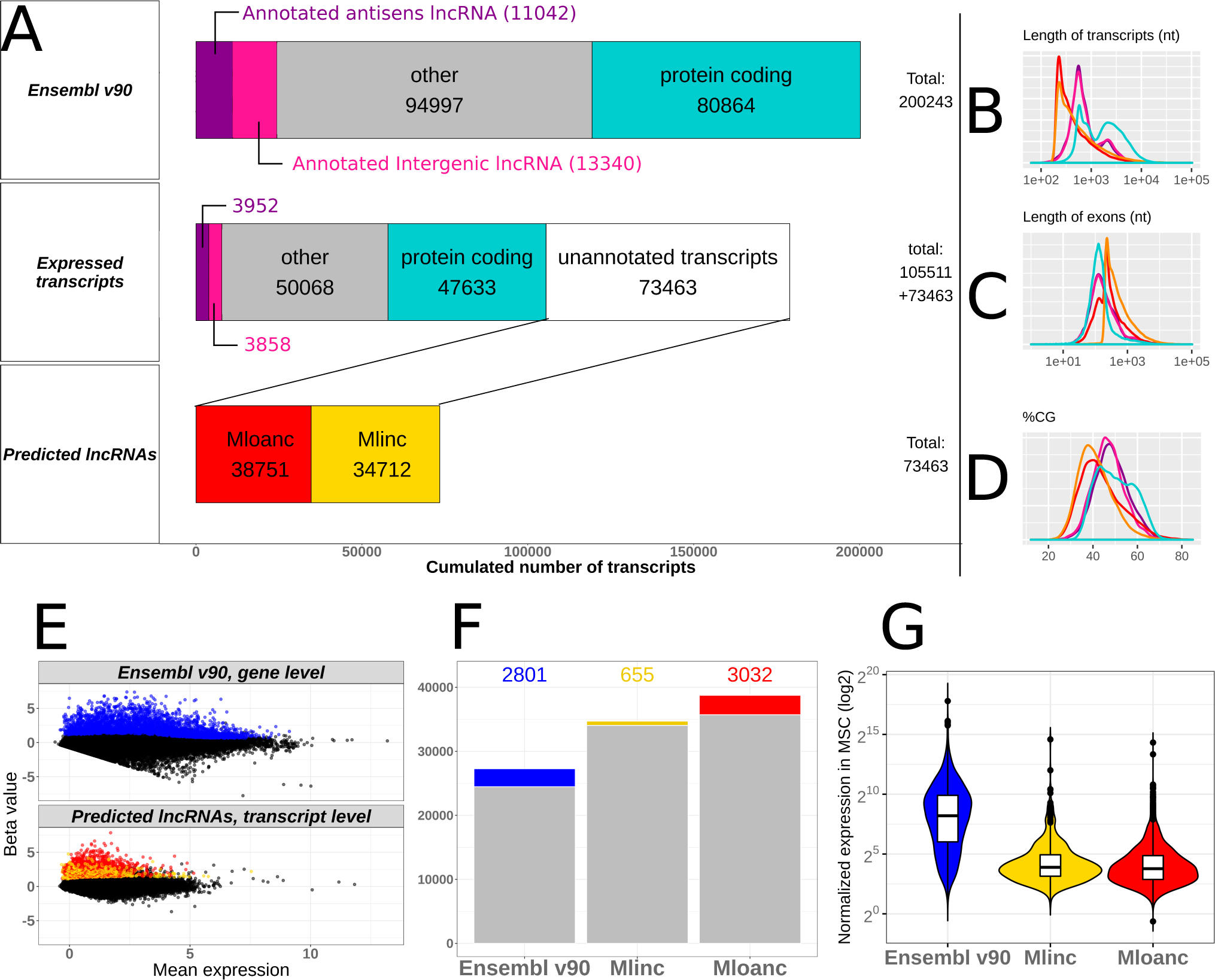
Overview of annotated genes and unannotated transcripts enriched in BM-MSC. A) Left pannels represented: i/EnsemblV90 transcript categories and distribution, ii/ transcripts distribution expressed in MSCs, showing unnatotated transcripts obtained with Ab initio reconstruction by StringTie vs annotated transcripts (expression *>* 0.1 TPM) iii/ Predicted long non-coding RNA(lncRNA) from unnanotated reconstructed transcripts include new long non coding RNA with intergenic (Mlinc) and antisens (Mlncoa) RNA categories. B-C-D) Distribution of transcript length, exon length and GC percentage across different categories respectively with the same colors as in A pannel : coding transcripts (blue), annotated lincRNA (pink), annotated overlapping antisens lncRNA (purple), novel lincRNA (Mlincs, yellow), novel overlapping antisens RNA (Mloanc, red). E) Representation of annotated genes (top pannel) and unannotated transcripts (bottom pannel) overexpresed in MSC versus non-MSC types (log2FC *>*0.5 and padj *<* 0.05), separately showed in MA plot. F) Total number of transcripts transcript by category. The colored bar indicated the number of differential expression of annotated genes (Ensemblv90) and unannotated transcripts (Mlinc and Mloanc). G) Global expression in BM-MSC (with Sleuth normalization) of the same categories as in F for annotated genes and unannotated transcripts.

The *ab initio* method by itself is not sufficient to efficiently determine the lncRNAs’ full length sequences. Moreover, this step does not preclude the possibility of false positives and at this point of the analysis, all the different rebuilt transcripts are considered to be windows of RNA expression or possible artefacts. These candidates are filtered and, for most interesting candidates, their true form is to be refined through experimental methods. We also assessed the general characteristics of predicted *de novo* lncRNAs in MSCs. Globally, Mlincs and Mloancs are shorter transcripts with longer exons compared to coding genes and annotated lncRNAs. The large majority of predicted lncRNAs are mono exonic (99% for Mlincs, 79% for Mloancs), with a length close to 200nt (Figure 2B-C). A consequence of the abundance of mono-exonic lncRNAs is an infinitesimally small number of variant forms. Only 0.15% and 0.82% of Mlincs and Mloancs are not mono isoforms, respectively.

The GC content of reconstructed lncRNAs is lower than that of coding or non-coding annotated genes (Figure 2D). This low GC proportion of around 40% is a common feature in *ab initio* transcript prediction, observed in a majority of studies of different species, from mammals, insects, plants or prokaryotes [41, 42, 43, 44].

### 2.15 Enrichment of a restricted set of Mlincs and Mloancs

In this second step, our objective was to obtain a restricted set of potential transcripts, using successive filtering approaches that would reveal their cell specificity. We quantified annotated transcripts, Mlincs and Mloancs with kallisto pseudoalignment [26] in a cohort constituted of two groups : the “MSC” group contained the BM-MSCs initially used for *ab intio* reconstruction, and the “non-MSC” group used for comparison, composed of a large panel of different cell types including hESC, hematopoietic precursors and stem cells, primary chondrocytes, IPS, hepatocytes, neurons, lymphocytes and macrophages (Table S1).

Only over-expressed transcripts in “MSC” group versus “nonMSC” group were selected. Differential statistical tests were made with Sleuth, a tool specially dedicated to Kallisto results [27] (see all selective parameters in Materials and Methods). We performed two differential expression analyses: one at the gene level for Ensembl annotation, and the other at the transcript level for unannotated transcripts, to give the most likely variant form of the predicted lncRNAs. After this differential analysis, 2801 annotated genes, 655 Mlincs and 3032 Mloancs are significantly overexpressed in BM-MSCs (Figure 2 E-F).

The lncRNAs are commonly known to be less expressed than coding genes and this was observed in our selected annotated genes and new lncRNAs (Figure 2G). As a validation of our procedure, we found the 3 positive MSCs markers of ISCT among the selected annotated genes: THY1 (CD90), ENG (CD105), and NT5E (CD73). We also retrieved some influencers of MSCs activity, for example WNT5A [45, 46], Lamin A/C [47], FAP [48] and others. The complete list of selected genes is provided in Table S5.

### 2.16 Feature selection for the most discriminating coding and non-coding markers

In an attempt to select the best candidates, we retained lncRNAs with the most discriminative profile between “MSC” and “non-MSC” groups. In our case, the limitation with a classical “top” ranking by Fold Change (FC) or p-value is the presence of subgroups of different types of cells inside the negative “non-MSC” group. The FC, estimated by the Beta value in Figure 2C, appears to be a biased indicator of differential expression as it can select strong but localised expressed lncRNAs in cells poorly represented in our negative group, leading to potential false positive results.

To avoid this problem, we used the Boruta feature selection (see material and method section), for selection of discriminating features based on “random forest” machine learning methodology. Boruta was used separately on each group of candidates (annotated genes, Mlincs and Mloancs) to extract a restricted representation of the most relevant MSCs signatures. The top 35 importance scores were selected for genes, Mlincs and Mloancs. We arbitrary chose to select the first 35 transcripts for each group based on our observation of a drop in the importance score in coding the gene series. Considering the expression profile of the top 35 coding genes and predicted Mlincs, BM-MSCs clusterised independently from other cells types (Figure 3A).

**Figure 3:**
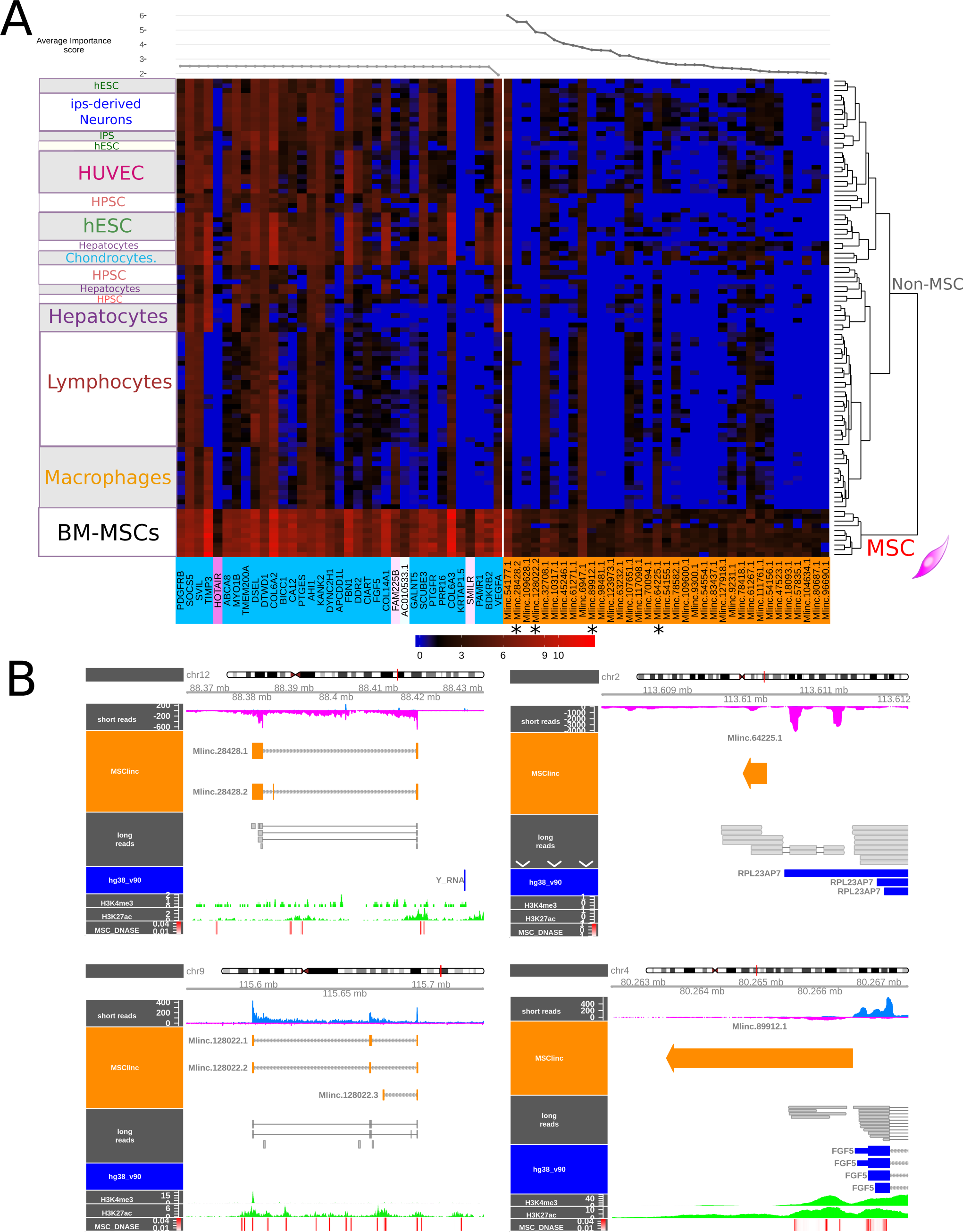
Selection of a refined set of best candidates by random forest (top35), long read sequencing and epigenetic features. A) Expression of the best MSC-specific candidates selected by boruta machine learning along MSC group and not MSC cohorts. Left pannel : top35 most relevant annotated genes (non-coding included); Right pannel: unannotated intergenic lncRNAs (Mlincs) and their average importance scores determined by Boruta method displayed in upside line plot. B) Genomic visualisation of Mlincs predictions 28428, 128022, 89912, and 64225 (MSClinc orange) from short reads alignement of all MSC group files (blue/magenta and bam visualisation), compared with long read alignements (long reads grey). Additional epigenomic features are shown to reveal active transcriptional activity from trimethylation of Histone H3 (H3K4me3), acetylation of Histone H3 H3K27 in MSCs (H3K4me3 and H3K27ac, green), and Dnase sensibitity hotspots of MSC (MSC DNAse, red).

In contrast, the selection of Mloancs did not provide a satisfying clustering as they had similar expression profiles in MSCs and other closely related cell types, in particular in primary chondrocytes (Figure S1). For this reason, Mloancs were not retained for further analysis. Selected annotated genes showed a poor specificity, with only few candidates showing a clear difference of expression between MSC and others: APCDD1L, HOTAIR, KRTAP1-5 and SMILR. The 3 positive MSCs markers from the ISCT were absent in this selection. The novel top 35 Mlincs showed less expression overall but with a more distinctive profile and a higher number of possible MSCs markers with clear contrast of expression. The characteristics, genomic intervals and sequences of the 35 candidates are presented in Table S7.

To assess the potential of genes already proposed as potential MSCs biomarkers by ISCT (Figure S2) or other potential MSCs markers proposed by different authors [14] (Figure S3), we made a separated expression heatmap, without filter. Among these previously proposed markers, THY1 (CD90) presented the most specific profile. However, each gene presented expression in distinct non-MSC types.

### 2.17 Validation of selected Mlincs with long reads sequencing

As mentioned above, classical annotation of lncRNAs with *ab initio* short read methods suffers from inaccuracies and biases. The Oxford Nanopore long read technology (ONT) can sequence entire cDNA, which constitutes a clear technological advantage, not only in confirming the existence of the transcripts but also as it makes it possible to precisely identify the genomic intervals of lncRNA candidates. We performed long-read sequencing of a poly(A) RNA library obtained from a BM-MSCs sample. Among the top 35 selected Mlincs, with around 3 million total reads, 4 transcripts are covered with the ONT sequencing.

These intergenic lncRNAs are named as Stringtie output (SetName. TranscriptNumber. VariantNumber): Mlinc.28428.2, MlincV4.128022.2, MlincV4.89912.1 and MlincV4.64225.1. To support the above transcriptional units, we compared them with our short read data and searched for epigenetic status at the locus of the Mlincs in bone-marrow stromal mesenchymal cells. We looked at DNase sensitive site, H3K27 acetylation, H3K4 trimethylation that commonly corresponds to active regulatory regions (Figure 3B). We globally observed a DNA accessibility enrichment and acetylation of Histone 3 at the promotor region of our candidates, correlating with DNAse sensitivity hotspots in BM mesenchymal cells that reinforce the prediction of the expression windows. In particular, for Mlinc.28428.2, the transcript observed with long reads sequencing corresponded to the prediction made with short reads. It was also supported by Mlinc.28428.1, a variant that differ by the absence of the second exon. Similar characteristics were observed for Mlinc.128022, which also produced two variants with a different organisation of 5 exons. The two other candidates, Mlinc.89912.1 and Mlinc.64225.1 are mono-exonic. Mlinc.89912.1 occurs at the close proximity of FGF5 3’ end, in reverse orientation at this locus. For this reason, the different epigenomic features could not be attributed with certainty to the Mlinc. For Mlinc.64225.1, the sequence is longer than the *ab initio* short read prediction. KRTAP1-5, HOTAIR and SMILR, selected for their good expression profile, were also covered by long reads (Data not shown).

### 2.18 High-throughput investigation of a marker’s specificity by specific k-mers search

A marker can only be considered specific within the limits of the diversity of samples used for its study. Considering the growing number of cell/tissues and transcriptional profiles, it is essential to probe the limits of a chosen biomarker against these various cell types. Most of published analyses highlighting new potential biomarkers of MSCs or fibroblasts have been restricted to a comparison between only few cell types and, as discussed, commonly described markers are not strictly distinctive. In order to assess the expression of Mlincs candidates in a large number of samples, we extracted specific 31nt k-mers from each of their sequences, as previously described [49]. These simplified but candidate-specific (oligonucleotide-like) probes allow a simple and fast presence/absence search to be made on large-scale cohorts and a direct quantification in raw fastq data. The k-mers were quantified in ENCODE human RNAseq database, including “primary cells” and “*in vitro* differentiated cell” categories (Table S2). Particularly, as the bibliography suggests that MSCs can also express phenotypic characteristics of endothelial, neural, smooth muscle cells, skeletal myoblasts and cardiac myocytes, RNAseq samples from this mesodermal origin were tested.

With ISCT positive markers, we observed an expected expression profile that recapitulate previous biological studies, particularly the high expression of Endoglin (ENG, CD105) in endothelial cells (Figure S4) and the overexpression of NT5E (CD73) in epithelial and endothelial cells (Figure S5). Interestingly, their expression varied among MSCs cell sources : NT5E (CD73) was strongly enriched in adipose and BM derived MSCs, and THY1 (CD90) in umbilical cord derived MSCs (UC-MSCs) (Figure S6). We next analysed the expression profile using our candidate annotated genes Mlinc specific k-mers (Figure 4). The specific k-mers search supported the stated expression profile of Mlincs previously shown : our Mlinc candidates were positive in MSCs and displayed weak or absent expression in cells of ectodermal lineage, hematopoietic or endothelial origins.

**Figure 4:**
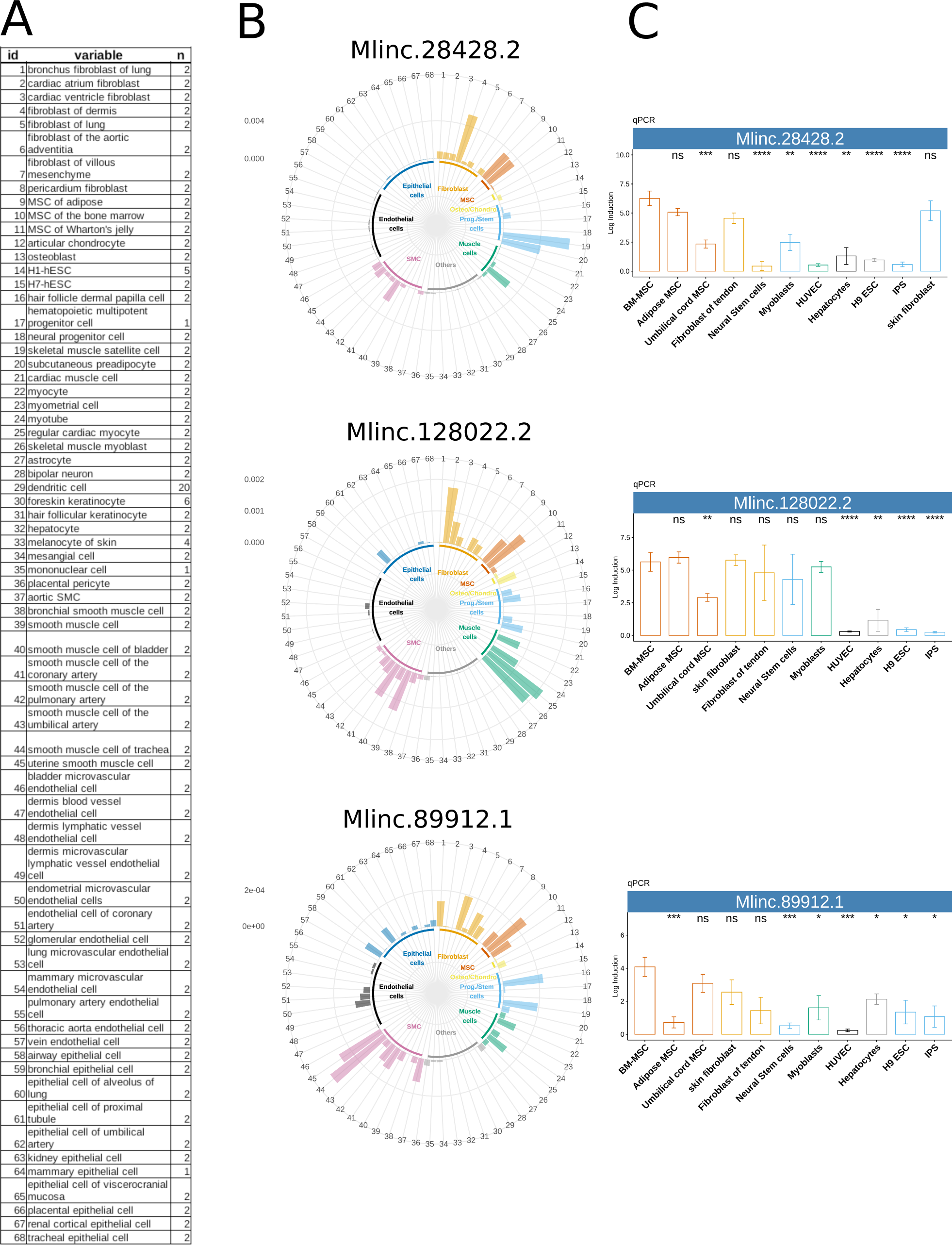
High throughput exploration of selected candidate across strong variety of samples by k-mer quantification in RNAseq and biological validation by RT-qPCR. A : List of tissues for the cell specific expression exploration (samples with ID numbers are listed in Table S1) B: Relative expression of Mlinc.28428.2, Mlinc.128022.2, and Mlinc.89912.1 across ENCODEs ribodepleted RNAseq datas, made by k-mer quantification, normalized by k-mer by million. C: qPCR relative quantification was performed on the selected 3 Mlincs in MSC of different origins (BM-MSC, Ad-MSC, Umbilical cord msc) and other indicated cell types. Relative quantification (Log induction) was quantified by ddCt method using non MSC types types as calibrator (mean of triplicates). Student tests have been made between triplicates, each test use BM-MSCs as reference group (ns : P *>* 0.05, *: P *≤* 0.05, **: P *≤* 0.01, ***: P *≤* 0.001, ****: P *≤* 0.0001).

However the high throughput and naive quantification in the ENCODE cohort made it possible to extend the observation of this absence of expression into cell types not previously studied. Moreover, this diversity showed that the expression of most of the candidates, contrarily to positive markers of the ISCT, were exclusive of cells with mesodermal origin. All candidates were expressed in at least one type of fibroblasts and differentially present in other mesodermal cell types. For the 4 selected Mlincs, they shared: (i) a systematic and strong expression in cell types like skin fibroblasts, and cells derived from reservoir of mesenchymal progenitors (muscle satellite cells or dermis papilla cells), (ii) a regular over-expression in regular cardiac myocytes, (iii) a significant and irregular expression in smooth muscle cells. The ENCODE cohort containing MSCs of different origins, we can therefore observe that the Mlincs show differences of expression depending of the tissular origin, these candidates being mainly expressed in two MSCs types. The results permitted the classification of our Mlincs according to observed specificity, from the most promising to the least restricted profile : Mlinc.28428.2 is expressed in Ad and BM derived MSCs. It is the candidate with the clearest absence of expression in non-mesodermal cells, and with the poorest relative expression in Smooth Muscle Cells (SMC). Mlinc.128022.2 is expressed in Adipose and Bone Marrow MSCs, and particularly in preadipocytes and muscle cells : (myoblasts, myocytes and myotubes). Mlinc.89912.1 is principally expressed BM-MSCs and less in UC-MSCs and AD-MSCs, but shows expression in epithelial and endothelial cell. Finally, Mlinc.64225.1 differs from other Mlincs as it is also strongly expressed in keratinocytes, HSC, and epithelial cells (Figure S7). Its expression in critical non-MSC types has led us to retain the three other Mlincs for further investigations.

### 2.19 RT-qPCR mimics the *in silico* prediction and deciphers multiple transcript variants

To confirm the specificity of selected Mlincs’ expression experimentally, we performed RT-qPCR on a set of 80 RNA preparations from different primary cell (Figure 4C). These includes MSCs from Bone Marrow (BM), Adipose (Ad) and Umbillical Cord (UC), fibroblasts of different tissue origins, IPS cells, neural stem cells, myoblasts, HUVECs and hepatocytes (Table S6). RT-qPCR and amplicon sequencing using sets of specific primers (Table S7) confirmed different predicted forms of the Mlincs candidates in BM-MSCs (Figure S8). We designed two primer pairs for both Mlinc.128022 variants to validate the existence of first splice, and two pairs for Mlinc.28428 variants, one overlapping the second exon and another corresponding to a splice between first and third exon. All variations captured by the primer designs were quantified, suggesting that all these different variations predicted *in silico* exist biologically in MSCs. We confirmed most of the expression profiles obtained by k-mers quantification using RT-qPCR, notably the specificity of expression dependency on the MSCs tissular origin: over expression of Mlinc.28428 and 128022 in BM- and Ad-MSCs. Nevertheless, few exceptions, such as Mlinc.89912.1, present an enrichment in UC-MSCs not found in k-mers quantification. Moreover, the restricted expression to cells of mesodermal origin is replicated in our RT-qPCR results. We obtained similar observations with annotated candidates : overexpression of KRTAP1-5 and SMILR in BM-MSCs specifically, and of HOTAIR in UC- and BM-MSCs.

### 2.20 *In silico* prediction of lncRNAs interactions and functions

The relative specificity of selected Mlincs for mesenchymal cells could be an indication of their roles in MSCs function. The prediction of their possible function could therefore suggest their suitability as markers of MSCs’ function potential. To this end, we explored assumptions on the function of Mlinc.28428.2, Mlinc.128022.2 and Mlinc.89912.1 candidates using different published methods. We first used bioinformatic tools based on machine learning and deep learning to decipher general characteristics of our candidates : FEELnc to assess coding potential, tarpMir to decipher “miRNA sponge” function, and LncADeep to analyse potential interactions with proteins.Only two of the 35 selected Mlincs and none of the 3 selected Mlincs with validated specificity were revealed as potentially coding RNAs with the majority being predicted as non-coding by FEELnc (33/35). None candidate had more than five target sites for a given miRNA, indicating a low probability of a “miRNA sponge” activity (Table S7). For the 3 retained Mlincs, predicted interacting proteins by LncADeep were submitted to Reactome (Table S8).

We noted a predicted interaction between Mlinc.28428.2 and Betacatenin (CTNNB1) as part of apoptosis-linked modules, 5’-3’ Exoribonuclease 1, component of the CCR4-NOT complex, mRNA Decapping Enzyme 1B as part of the mRNA decapping and decay pathways. The interaction was also predicted with different mediators of RNA polymerase II transcription subunits (MED), ATP Binding Cassette Subfamily B Members as part of the PPARA activity linked to ER-stress [50], and Proteasome subunits for intracellular transport, response to hypoxia and cell cycle modules. Mlinc.128022 could interact with important genes like THY1 (CD90), NRF1 (mitochondria metabolism) with no module clearly highlighted. Mlinc.89912 could interact with tubulins, UBB (ubiquitin B), SMG6 nonsense mediated mRNA decay factor and ribosomes subunits (RPSX) proteins, (RPL24) for nonsense Mediated Decay (NMD), PINK1 (mitophagy) and finally MGMT, as part of the MGMT mediated DNA damage reversal module.

We further quantified expresssion of candidates by counting their specific k-mers in the entire FANTOM6 set of 154 Known-Downed (KD) annotated lncRNAs in human dermal fibroblasts (https://doi.org/10.1101/700864, not peer-reviewed). We selected the KD experiments where expression of the Mlincs was statistically differential when compared with controls. Particular attention was paid to KD lncRNAs with reported function(s) in bibliography, and to KD lncRNAs overlapping a gene with reported functions. Mlinc.28428.2 is down-regulated when JPX, SERTAD4-AS1, BOLA3-AS1, and SNRPD3 are KD, and over-expressed with the KD of PTCHD3P1, ERVK3.1, MEG3, among other lncRNAs without reported function (Figure 5A). Interestingly, interactions between p53 pathway and JPX [51], SNRPD3 [52] and MEG3 [53, 54]) respectively, have been previously reported. All these features converge on the hypothesis of a link between the function of Mlinc.28428, stress response, senescence and cellular maintenance. The implications of BOLA3 [55, 56]) and PTCHD3P1 [57] in mitochondria homeostasis and glycolysis, the role of BOLA3 in stress response [58], the status of SERTAD4 as target of the YAP/TAZ pathway [59], vital pathway of stress response [60], and the role of MEG3 in aging [61], all reinforce this hypothesis.

**Figure 5:**
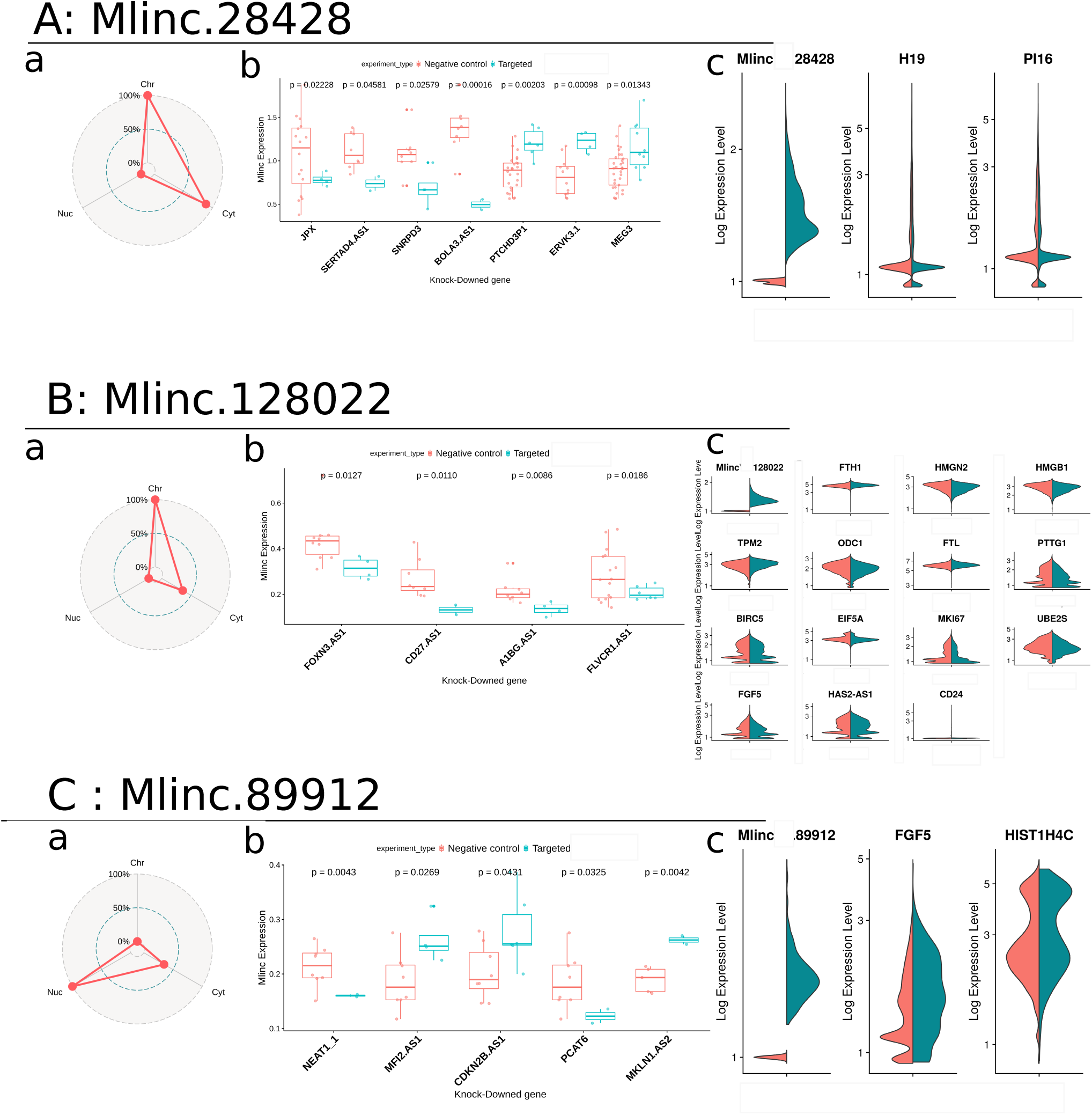
Prediction of potential functions of candidates with k-mer quantification and single-cell. For each Mlinc (Mlinc.28428 (A), Mlinc.128022 (B) and Mlinc.89912 (C) respectively) 3 steps of prediction were performed. a. Enrichment in the different subcompartments of fibroblasts from FANTOM6 dataset: Free nuclear fraction (Nuc), chromatin (Chr) and cytoplasm (Cyt); b. Expression of marker in FANTOM6 data depending of the KnockDown (KD) of an annotated lncRNA. Normalised count of all specific k-mers is averaged by sample (zeros vlues deleted) and t-tests are made between control and KD fibroblasts. c. General expression of Mlincs inside Ad-MSC population, dimensionnal reduction made with UMAP method, made from batch corrected counts. Expression of differentially expressed annotated genes between positive and negative cells for Mlinc.28428, Mlinc.128022 and Mlinc.89912 respectively.

Mlinc.128022.2 is down-regulated with the KD of FOXN3-AS1, A1BG-AS1, CD27-AS1, and FLVCR1-AS1 (Figure 5B). FOXN3 seems to be more than a regulator of cell cycle, it is also described as a regulator of osteogenesis in different cases of defective craniofacial development [62, 63]. Moreover, the reported over-expression of FOXN3 during the early stages of MSCs osteodifferentiation [64], and downregulation of CD27-AS1 in MSCs of donors with bone fracture [65], allow us to hypothesise a possible function of Mlinc.128022 in bone remodelling and osteogenesis. In addition, both A1BG-AS1 and FLVCR1-AS have an influence in osteogenesis and cell differentiation. A recent study showed that A1BG-AS1 interacts with miR-216a and SMAD7 in suppressing hepatocellular carcinoma proliferation [66], both partners having an important role in the positive regulation of osteoblastic differentiation in mice [67] [68]. FLVR1 participates to resistance of oxydative stress by heme exportation in mouse MSCs [69], iron metabolism being closely linked with bone homeostasis, formation [70] and cell differentiation [71].

Finally, Mlinc.89912.1 is down-regulated after the KD of NEAT1-1 and PCAT6, and over-expressed when MFI2.AS1, CDKN2B.AS1 (or ANRIL) and MKLN1.AS2 are KD (Figure 5C). The manifest relations between cell proliferation and CDKN2B-AS1 [72, 73], MFI2 [74], MFI.AS1 [75], PCAT6 [76] and NEAT1 [77, 78], an combination with the between ones and DNA damage repair response, [79, 80] reinforces the prediction of a role of Mlinc.89912 in these mechanisms. Moreover, we explored RNAseq from chromatin, nucleus and cytoplasm subcellular compartments of fibroblastic cells in the FANTOM6 Dataset. Mlinc.28428 and Mlinc.128022 are enriched in at least cytoplasm, whereas Mlinc.89912 is enriched in free nucleus fraction suggesting interaction with nuclear component (Figure 5C).

### 2.21 The single cell RNAseq: an emergent level of completion in marker search

We analysed the single cell RNAseq data from 26 071 adipose MSCs (Ad-MSCs) to assess the heterogeneity of the 3 Mlincs and to explore their expression at the single-cell level. No clear correlation between cell cycle and expression of our Mlincs was identified (Figure S9). We observed a high variability of the number of cells expressing the markers (Threshold *≥* 0.1). 11 927/26 071 were Mlinc.28428-positives, 4944 were Mlinc.128022-positives, and 404 were Mlinc.89912-positives.

For each Mlinc, we performed a differential test to decipher genes differentially expressed in Ad-MSCs Mlinc-positive and Mlinc-negative cells.

We found that Mlinc.28428-positive cells under-expressed H19 and PI16 (Figure 5A). These genes, that present a diversity of functions, are involved in stress mechanisms (oxydative response and shear stress), inflammation in fibroblasts and MSCs, and senescence pathways [81, 82, 83, 84]. Despite the low number of differentially expressed genes in Mlinc.28428-positive cells, their functional behaviour and their known targets suggest a pathway linked to stress response and senescence establishment that reinforce our previous assumptions on Mlinc.28428 function.

Mlinc.128022-positive cells are enriched in FTH1, TPM2, FTL and CD24 and present a lower expression in HMGN2, HMGB1, ODC1, PTTG1, BIRC5, EIF5A, MKI67, UBE2S, FGF5, HAS2-AS1 (Figure 5B). A significant portion of these genes are linked to osteogenic properties of MSCs as previously observed with FANTOM analysis. The Mlinc.128022-positive cells have an increased expression of ferritin (light and heavy chains), major actor in iron metabolism in osteoblastic cell line [85], that is also involved in osteogenic differentiation [86] and osteogenic calcification [87]. Two genes, enriched in Mlinc.128022-positive cells are positively linked to the osteogenic differentiation potential of MSCs : the tropomyosin 2 (TPM2), downregulated when hMSCs were cultured in OS medium for the induction of osteoblasts at the calcification phase [88], and CD24 a membrane antigen recently proposed as a new marker for the sub-fraction of notochordal cells with increased differentiation capabilities [89]. In addition ODC1, under-represented in Mlinc.128022-positive cells, inhibited the MSCs osteogenic differentiation [90, 91]. Finally, the decrease of FGF5, MKI67, BIRC5 (Survivin) and PTTG1 (securin) expressions, all linked to proliferation active phases of cell cycle, tend to show cell with arrested cell cycle. These data suggest that the expression profile of Mlinc.128022 positive cells indicate a subpopulation of undifferentiated osteogenic progenitors, probably in senescence or quiescence.

Mlinc.89912-positive cells are enriched in FGF5 and HIST1H4C (Figure 5C). FGF5 is a protein with mitogenic properties, identified as an oncogene, that facilitates cell proliferation in both autocrine [92] and paracrine manner [93]. HIST1H4C, the Histone Core number 4, is a cell cycle-related gene. Modification of histone H4 (post-transcriptional or mutation) has been highlighted as important for Non-Homologous End-Joining (NHEJ) in yeast [94]. Its mutation cause genomic instability, resulting in increased apoptosis and cell cycle progression anomalies in zebrafish development. It reinforces our assumptions that Mlinc.89912 has a role in cell proliferation and DNA damage repair. In conclusion, the single cell RNAseq analysis enabled the observation of different features that characterise the phenotype of Mlincs positive cells and reinforced hypotheses on their functions previously observed through k-mers quantification.

### 2.22 K-mers analysis of markers in functional cell situation

Previously, we have presented a number of strategies to formulate hypotheses on the functions of an unannotated lncRNAs, suggesting directions of future experimental investigations. To evaluate the relevance of these strategies, we sought to quantifiy with specific k-mers search the expression of our Mlincs in MSCs in different conditions, linked to above mentioned findings: stress and senescence for Mlinc.28428.2, osteodifferentiation for Mlinc.128022.2 and cell cycle/proliferation for Mlinc.89912. We downloaded RNAseq data corresponding to the above-mentioned focus, described in Table S4.

As shown in Figure 6, we observed a statistically relevant increase of Mlinc.28428 expression in MSCs under replicative stress and in MSCs with CrisprCas9 depletion of genes with important role against senescence. In the Wang et al. study [95], MSCs senescence was observed with the KO of ATF6 and the stress induced with tunicamycin (endoplasmic reticulum stress) and late passage (replicative stress). Mlinc.28428 expression increased with tunicamycin treatment, late passage and ATF6 KO. The highest increase is observed in ATF6 KO MSCs associated with late passage condition.

In Fu et al. study [96], YAP, but not TAZ, was found to safeguard MSCs from cellular senescence as shown by KO experiments. Interestingly, YAP KO, but not TAZ, significantly increases the expression of Mlinc.28428.2. This would lead us to conclude that Mlinc.28428 is overexpressed in senescence and stress conditions, suggesting a role in one or both of these phenomena.

The change in Mlinc.128022 expression is strictly linked to osteodifferentiation conditions. Mlinc.128022 expression shows a relevant increase in MSCs exposed to fungal metabolite Cytochalasin D (CytoD). The cytoD is reported as a osteogenic stimulant in the concerned study [97]. Moreover, no expression variation was observed between MSCs and MSC-derived adipocytes from Wang et al. study, implying a role in adipodifferentiation. Agrawal Singh et al. have studied osteogenic MSCs differentiation [29], with a similar increase of Mlinc.128022 being observed after ten days.

We then quantified the expression of Mlinc.89912 in a study that compare proliferating MSCs versus confluent MSCs [98, 99]. Our candidate was clearly over-expressed in proliferating cells, validating its capacity to mark the MSCs in proliferation. Moreover, its expression was not statistically modified when MSCs were exposed to EGF with pro-mitotic capabilities [100]. However Mlinc.89912 expression was reduced when IWR-1, an inhibitor of Beta-catenin nuclear translocation, that reduced the proliferation of MSCs, was added to the medium. The functional domains of these genes is summarised in table 1 and confirm the potential functional role suggested from FANTOM data: stress-related pathways for Mlinc.28428, MSCs differentiation with a presumed orientation in osteo-progenitors for Mlinc.128022 and a more restricted role in proliferation and DNA repair for Mlinc.89912.

**Figure 6:**
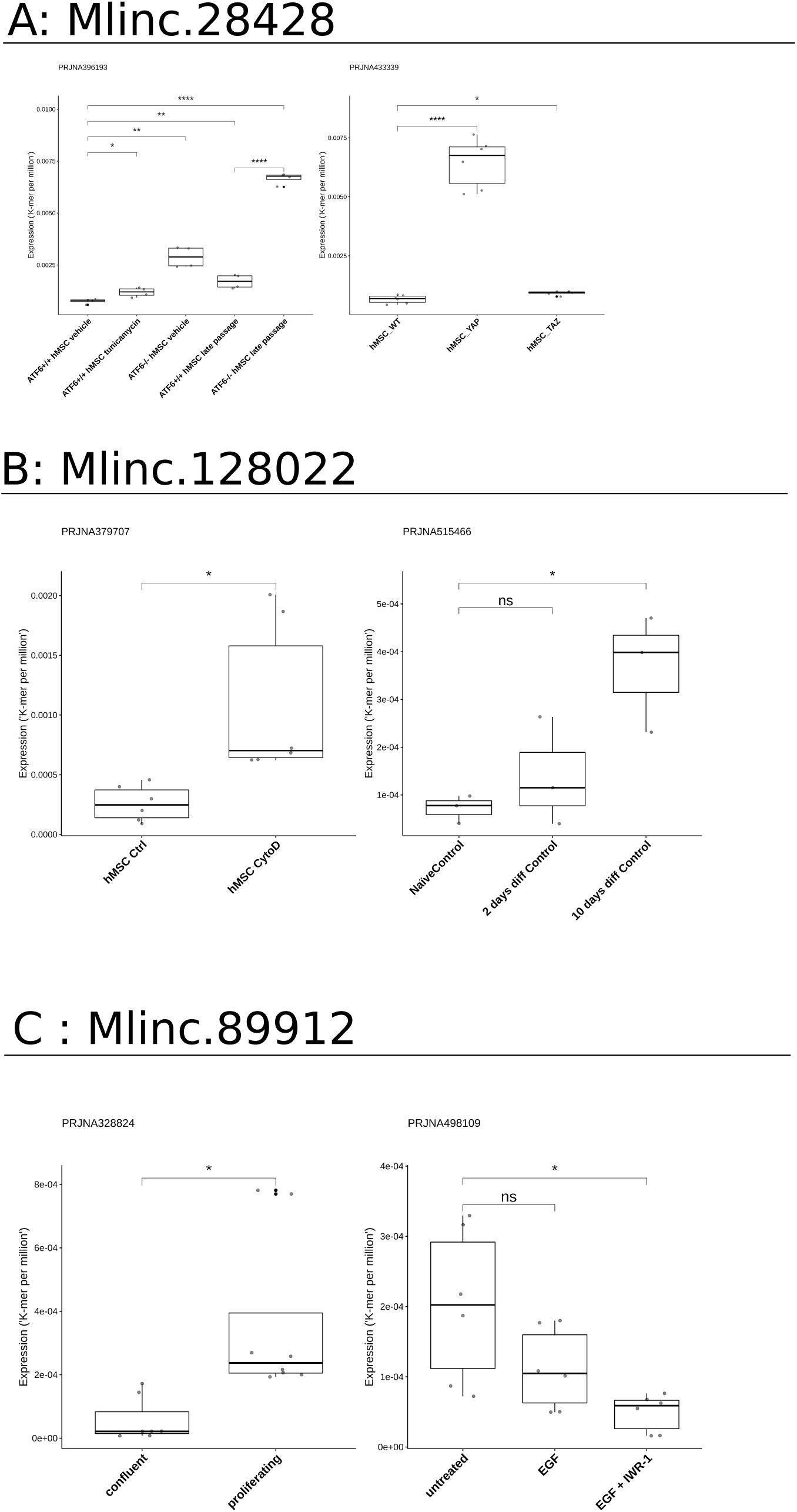
Expression of markers in different datasets from SRA in cell conditions related to previous findings. A) Expression of Mlinc.28428.1 in the context of oxydative, replicative, or KO-driven, stress and senescence (PRJNA396193, PRJNA433339). Relevant changes of expressions are showed with t-test results (ns : P *>* 0.05, *: P *≤* 0.05, **: P *≤* 0.01, ***: P *≤* 0.001, ****: P *≤* 0.0001). B) Expression of Mlinc.128022 in osteodifferentiation conditions (PRJNA515466) or osteodifferen-tiation potential (PRJNA379707). Relevant changes of expressions are showed with t-test results (ns : P *>* 0.05, *: P *≤* 0.05, **: P *≤* 0.01, ***: P *≤* 0.001, ****: P *≤* 0.0001). C)Expression of Mlinc.89912 in the context of proliferation (PRJNA328824 and PRJNA498109). Relevant changes of expressions are showed with t-test results (ns : P *>* 0.05, *: P *≤* 0.05, **: P *≤* 0.01, ***: P *≤* 0.001, ****: P *≤* 0.0001). The detailed list of datasets is provided in Table S4.

**Table 1.**
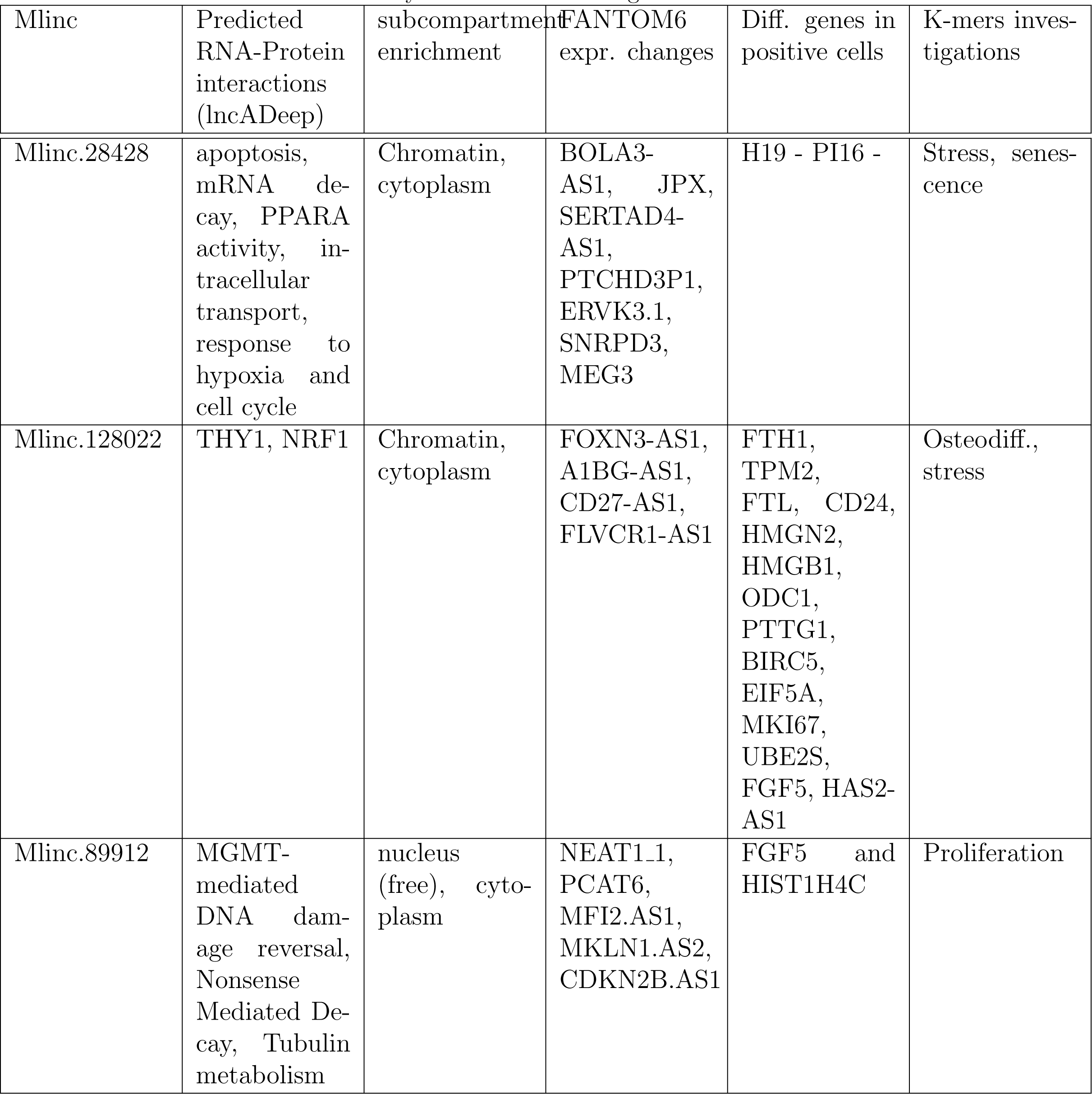
Summary of functional investigation results

| Mlinc | Predicted RNA-Protein interactions (lncADeep) | subcompartment enrichment | FANTOM6 expr. changes | Diff. genes in positive cells | K-mers investigations |
| --- | --- | --- | --- | --- | --- |
| Mlinc.28428 | apoptosis, mRNA decay, PPARA activity, intracellular transport, response to hypoxia and cell cycle | Chromatin, cytoplasm | BOLA3-AS1, JPX, SERTAD4-AS1, PTCHD3P1, ERVK3.1, SNRPD3, MEG3 | H19 - PI16 - | Stress, senescence |
| Mlinc.128022 | THY1, NRF1 | Chromatin, cytoplasm | FOXN3-AS1, A1BG-AS1, CD27-AS1, FLVCR1-AS1 | FTH1, TPM2, FTL, CD24, HMGN2, HMGB1, ODC1, PTTG1, BIRC5, EIF5A, MKI67, UBE2S, FGF5, HAS2-AS1 | Osteodiff., stress |
| Mlinc.89912 | MGMT-mediated DNA damage reversal, Nonsense Mediated Decay, Tubulin metabolism | nucleus (free), cytoplasm | NEAT1.1, PCAT6, MFI2.AS1, MKLN1.AS2, CDKN2B.AS1 | FGF5 and HIST1H4C | Proliferation |

## 3 Discussion

With recent evolution of omics analysis, the landscape of biomarkers has been extended beyond known genes to the exploration of the unexplored transcriptome. This potential has been assessed in pathological conditions but to a lesser extent in cell-specific conditions, where this new pool of potential markers could be used to identify less well-characterised cells and hence predict their function. In this article, we propose an integrated procedure and strategies to identify the best markers (annotated or not) in a cell-specific condition, and predict their potential functions, primarily from RNA sequencing data (Figure 1). RNAseq facilitates the creation of large lncRNA catalogues [8, 101] through the total catalogue of lncRNAs remains incomplete given the diversity of biological entities and lncRNAs specific expression in non-pathological, cell-specific conditions. The creation of a “home-made” catalogue associated with a specific condition remains the best way to assess the full diversity of potential biomarkers in a cell, rather than resorting to a global catalogue made from diverse samples. To give an idea of the completeness of such a focused lncRNA catalogue when compared to a global one, Jiang et al. recently published “an expanded landscape of human long non-coding RNA” with 25 000 new lncRNAs from normal and tumor tissues, whereas in our focused analysis only 50% of our 35 selected MSClinc can be found in this collection [101].

Futhermore, providing new candidates of good quality to improve lncRNA collection remains a complex task. As could be expected, the raw catalogue in our study contains predictions of disparate quality observed with a large number of mono-exonic transcripts. Without any filter, *ab initio* methods are insufficient to adequately reconstruct full length transcripts. The usage of long-read sequencing has been particularly effective in helping to validate our predictions. Given the benefits of full-length RNA sequence, long-read RNAseq should become the standard for lncRNA validation. A specific lncRNA can be the one presenting the most relevant properties after *in silico* analysis. The first task remain the identification of the more specific markers for a given cell type, task that present differences from classic comparative analysis. The MSCs markers proposed in the past were determined through a simple comparison between MSCs of a certain origin with negative cell whose types are either unique or few in number.

Historically, MSCs have been compared to bone marrow haematopoietic stem cells. However, our initial RNAseq analysis revealed that all potential MSCs markers proposed in the past are expressed in at least one other non-mesenchymatous cells type, and so do not constitute exclusive MSCs markers at the transcriptome level. Even if all cell types cannot be investigated, the diversity of the negative cell set is a critical criterion in selecting the most specific transcripts. In keeping with this idea, we then restricted the list of potential biomarkers with an enrichment step based on a differential expression comparing BM-MSCs to other cells including mainly stem cells, as well as differentiated cells of various lineages (lymphocytes, macrophages, primary chondrocytes, hepatocytes and neurons). In the enriched list, the overexpressed annotated genes contained members of MSC-related pathways as well as the ISCT markers. This result supported the MSCs characterisation made by the original authors [13], thus validating the identity of MSCs used for this RNAseq analysis with the currently defined criteria. The problem with classical differential analysis used on diverse “non-MSC” group is that all the “non-MSC” group is considered to be homogeneous. As a result, candidates with positive expression in small cell groups could pass statistical test, creating false positives. For this kind of differential analysis, we propose to select the most discriminating transcripts by feature selection, a machine learning methodology, that reduces the number of non-discriminating candidates after selection. We used feature selection through Boruta, a method based on random forest, to retain the top 35 of the most relevant MSCs signature for annotated genes, Mlincs and Mloancs separately. Putting aside our initial focus on unannotated lncRNAs, different annotated lncRNAs or coding genes with interesting profiles were also selected by feature selection : among them, KRTAP1-5 have been exclusively studied in BM-MSCs [102], where its preferential expression was validated by our results. These discoveries can bring new features concerning these genes and suggest directions for future investigations concerning their impact on the MSCs.

However, a marker is classically considered as specific on condition that its positive expression cannot be observed in any other cell type. Therefore, the expression of these potential markers should be explored in an entire RNAseq database to further validate its specificity. The exploration of a wide set of RNAseq data as proposed by ENCODE including a diversified set of primary and stem cells could support or invalidate the specificity of potential markers. In order to assess the expression of Mlincs candidates in a large number of samples, we used a signature for each candidate, extracting specific 31nt k-mers from their sequences. The specific k-mers extraction was made using Kmerator software. These k-mers were then quantified in the ENCODE human RNAseq databaseThe new and simplified procedure based on k-mers counting and large scale RNAseq exploration has the following advantages: i) a direct textual search that requires less time and CPU resources than classical methods, ii) a restricted set of lncRNAs supported by different results in the biological (wet) and *in silico* level (RNAseq data). The counterpart of the extensive vision of marker expression is that we observe a limit of specificity among our best candidates. We observed expression in fibroblasts, in close primary cells of common embryonic origin like smooth muscle cells (SMC) and other tissue-specific fibroblastic cells. Other tissue resident fibroblastic cells like skeletal muscle satellite cells, pre-adipocytes, and fibroblasts from different sources, especially dermis, express our selected MSClincs markers. The question of the differences between MSCs and related cell types is crucial to the issue. Specifically, the differences between MSCs and fibroblasts remain a subject of debate [103, 12]. According to the ISCT statement, no phenotypical differences have been reported between fibroblasts of different sources and adult MSCs [104], suggesting a hypothesis of a uniform cell type that show functional variation depending on the tissue source. Our results support this idea: distinguishing MSCs from fibroblast with only few positive markers remains a complicated task.

Moreover we observe low to medium expression of our candidates in close cell type from the same embryonic origin such as muscular cells and smooth muscle cells (SMC). This could be due to a shared phenotype between cells with close embryonic origin. Common markers between MSCs and SMCs have already been described. Notably, MSCs can express similar levels of SMC markers such as alpha-actin [105, 106]. Moreover Kumar et al. [107] determined that MSCs, pericytes, and SMCs could have the same mesenchymo-angioblast progenitor and that SMCs share a certain plasticity with MSCs as they can be differentiated in chondrocyte-like and beige adipocytes or myofibroblasts. However, a lot of cell types in ENCODE have not been actively sorted by expression of their respective surface markers, and fibroblast contamination is a classical feature in primary cell culture. We should not therefore exclude the possibility of fibroblast contamination when investigating marker for MSCs by bulk omics technology. Given this, single-cell RNAseq could be the best solution to identify the source of marker expression in counterpart cells.

To conclude, our extensive cell type comparison shows that the discovery of a marker of MSCs as distinct cell type is not plausible. After deepening our own research on MSCs biomarkers at the annotated and unannotated level, we were unable to find a marker that could simultaneously i) distinguish MSCs to close or homologuous cell types (fibroblasts, satellite cells, SMCs) ii) be present in all MSCs types iii) distinguish MSCs from more characterised cell types (Hematopoietic lineage, neurones etc). Our results suggest, like other studies, a strong proximity between MSCs, fibroblast and mesodermal cell types.

More than a marker of MSCs, candidates extracted by our method could be used to explore important features in MSCs biology and therefore warrant investigation into their function, assuming that the specificity of RNA for a cell type can highlight its importance in cell activity. Even if the functional invalidation stands as the principal method to efficiently determine the function of a lncRNA, its expression and co-expression with known genes can potentially characterise a function or an intrinsic state of a cell type, particularly for MSCs with reported diversity of states and function (ex : differentiation, immunomodulation, senescence, proliferation…). In our opinion, it is vital that during the creation of a catalogue of lncRNAs, a restricted set of selected biomarkers should be studied more intensively, both in term of specificity and functions. Assumptions on functional domains, where lncRNAs could act, could increase the relevance and visibility of discovered lncRNAs, and far from the bioinformatics implications, encourage future biological investigations. We decided to investigate the three selected MSClincs, validated by k-mers search, RT-qPCR and long-read sequencing, in term of biological impact with complementary *in silico* experimental approaches. We propose to use different *in silico* strategies, depending on the amount and diversity of the available data. The analysis confirms the non-coding potential of candidates and indicates a low probability of “miRNA sponge” activity. However, protein potential interaction results give interesting paths that were then investigated by complementary exploration. The k-mers quantification permits a naive high throughput exploration of numerous RNAseq data, simultaneously exploring potential functions and specificity to assess their potential. Instead of different cells, each candidate’s expression was quantified in MSCs in different experimental conditions. FANTOM6 data recently offered a pilot about lncRNAs functional investigation, with a high-throughput invalidation of 154 lncRNAs and coding genes in fibroblasts and their RNAseq counterpart added to phenotypical observations. The utilisation of co-expressions between knock-out genes and candidates lncRNAs remains an efficient way to decipher lncRNAs function, provided number of KD genes is high. Moreover, the availability of recent single cell data of MSCs have been a good complement to lncRNAs functional investigation.

Using scRNAseq from Ad-MSCs [19], we observed that our markers are not expressed in all cells but constitute different subpopulations with different levels of rarity in Ad-MSCs. FANTOM6 and single-cell analysis could permit tracing three components of these states : stress inducible cells, lineage commited osteogenic progenitors and proliferating cells. Globally, we observed a global concordance of the results between the different strategies used for functional prediction. Mlinc.28428 has concomitant expression with genes related to the stress response pathway. Mlinc.28428 could be a good target for treatment to study the senescence process, age pathologies or stress response. Mlinc.128022 potentially interacts with THY1 (CD90) and has co-occurences with genes linked to osteoprogenitors and cell differentiation. The k-mers search highlights its participation in MSCs’ osteo-differentiation. Finally, Mlinc.89912 potentially interacts with damage repair and RNA decay, and tubulin metabolism, all linked to cell proliferation and cell cycle. Moreover, the subcompartment enrichment corresponds to this prediction: Mlinc.89912.1 is the only candidate to have possible interactions with DNA-repair system, a hypothesis corresponding to his observed enrichment in the nucleus. A final selection of bulk RNAseq of MSCs in specific biological conditions allowed confirmation of our initial assumptions, showing that the different strategies we propose could be used to give relevant indication of the lncRNAs’ functions. These results show that a lncRNA selected by its expression specificity has a high probability of being part of a functional mechanism.

In conclusion, we have predicted genes and lncRNAs enriched in MSCs and proposed several selection steps including feature selection (machine learning), large scale signature search, RT-qPCR validation, *in silico* tools and single cell analysis. We present the application of a new way of quantification in RNAseq : The specific k-mers search could be used as a naive information in lncRNA catalogue creation. The strategies presented here are transferable to other cell types and different studies while the specificity and functional assumption present a significant potential in long non-coding transcriptome exploration. We present three lncRNAs markers of bone marrow and adipose MSCs that passed all selection steps and present interesting features: Mlinc.28428.2, Mlinc.128022.2 and Mlinc.89912.1. These markers could be used by the scientific community as potential targets for functional biological experiments on MSCs, with pre-indications of potential functions to orientate the experiments, and finally initiate the objective of transition between informatical problematics and cell biology.

## Acknowledgements

We thank for their generous gifts, G.Carnac for myoblasts, M.Le Quintrec-Donnette for HUVECs, E. Sanchez for dermal fibroblasts, D. Noel and ML. Vignais for mesenchymal stromal cells, C. Crozet for IPS, S. Gerbal and M. Daujat for hepatocytes. We thank Philippe Clair for his advice on qPCR, the qPHD plateform, Montpellier GenomiX and Jean-Marc Holder (SeqOne) for text corrections.

---

**Supplementary Figure 1:**
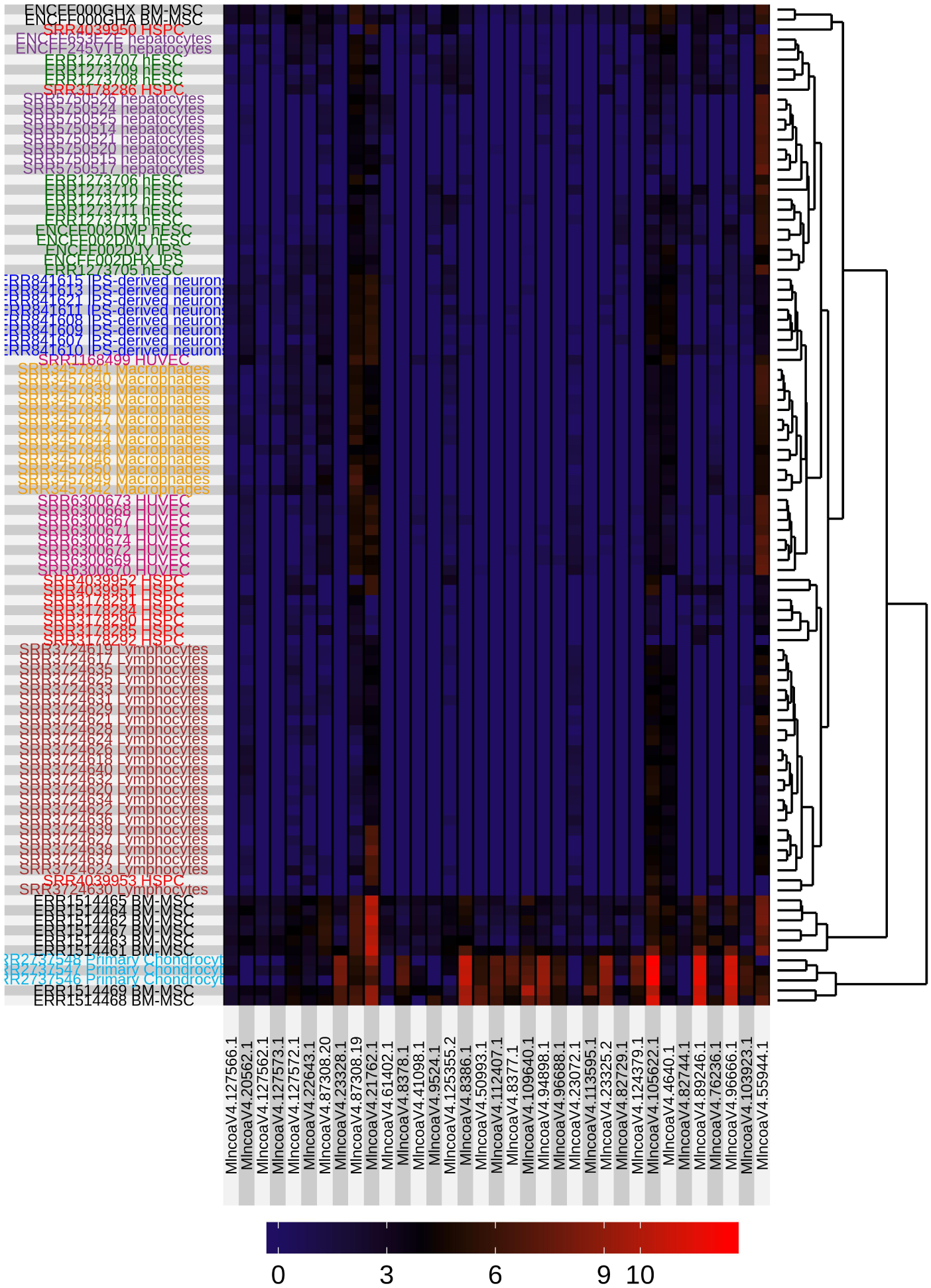
Expression of Mloanc (Antisens unannotated RNA) selected after feature selection in the differential analysis cohort

**Supplementary Figure 2:**
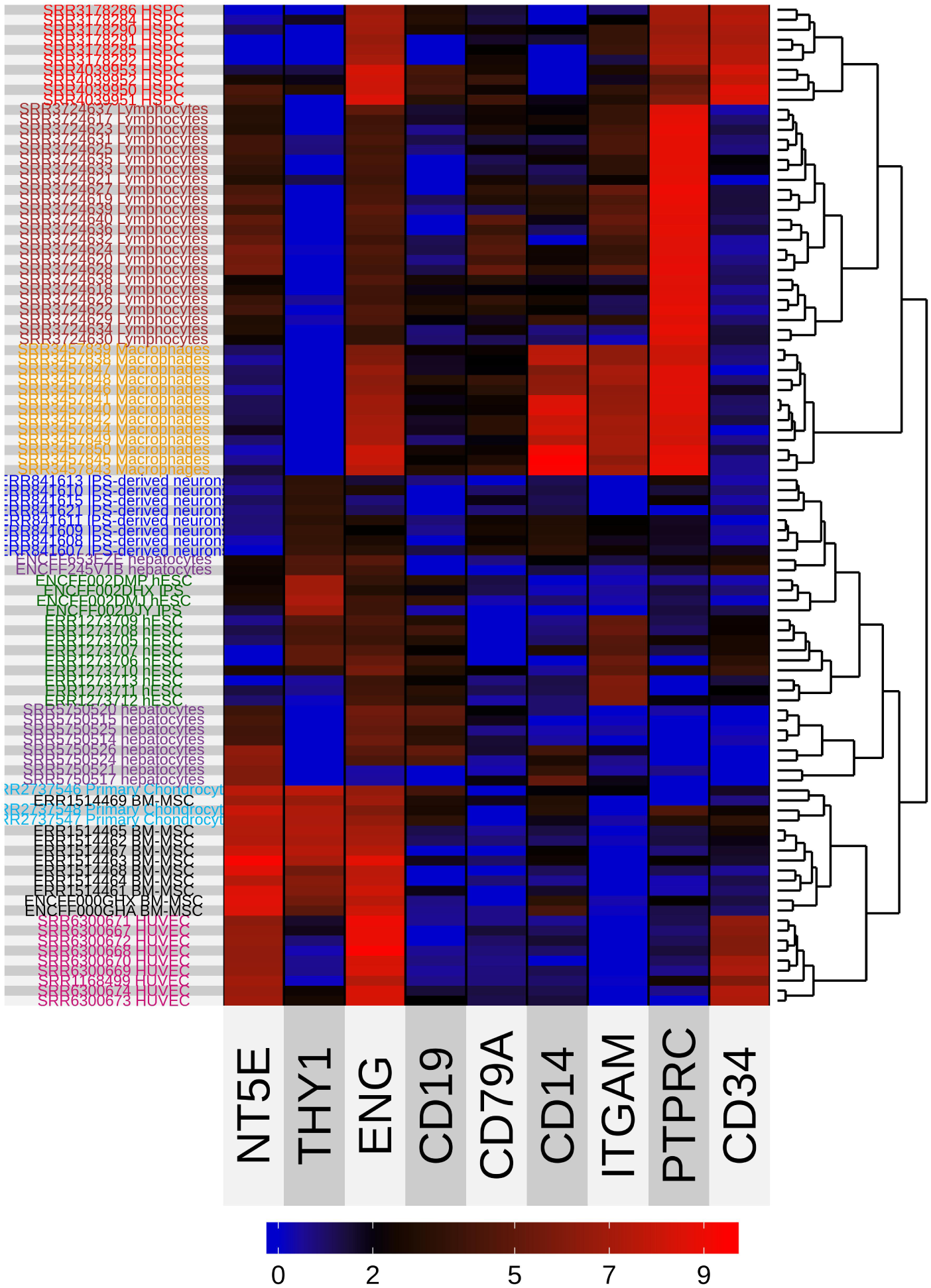
Expression of ISCT’s MSC markers in the differential analysis cohort THY1 = CD90, NT5E = CD73, ENG = CD105, ITGAM = CD11B, PTPRC = CD45

**Supplementary Figure 3:**
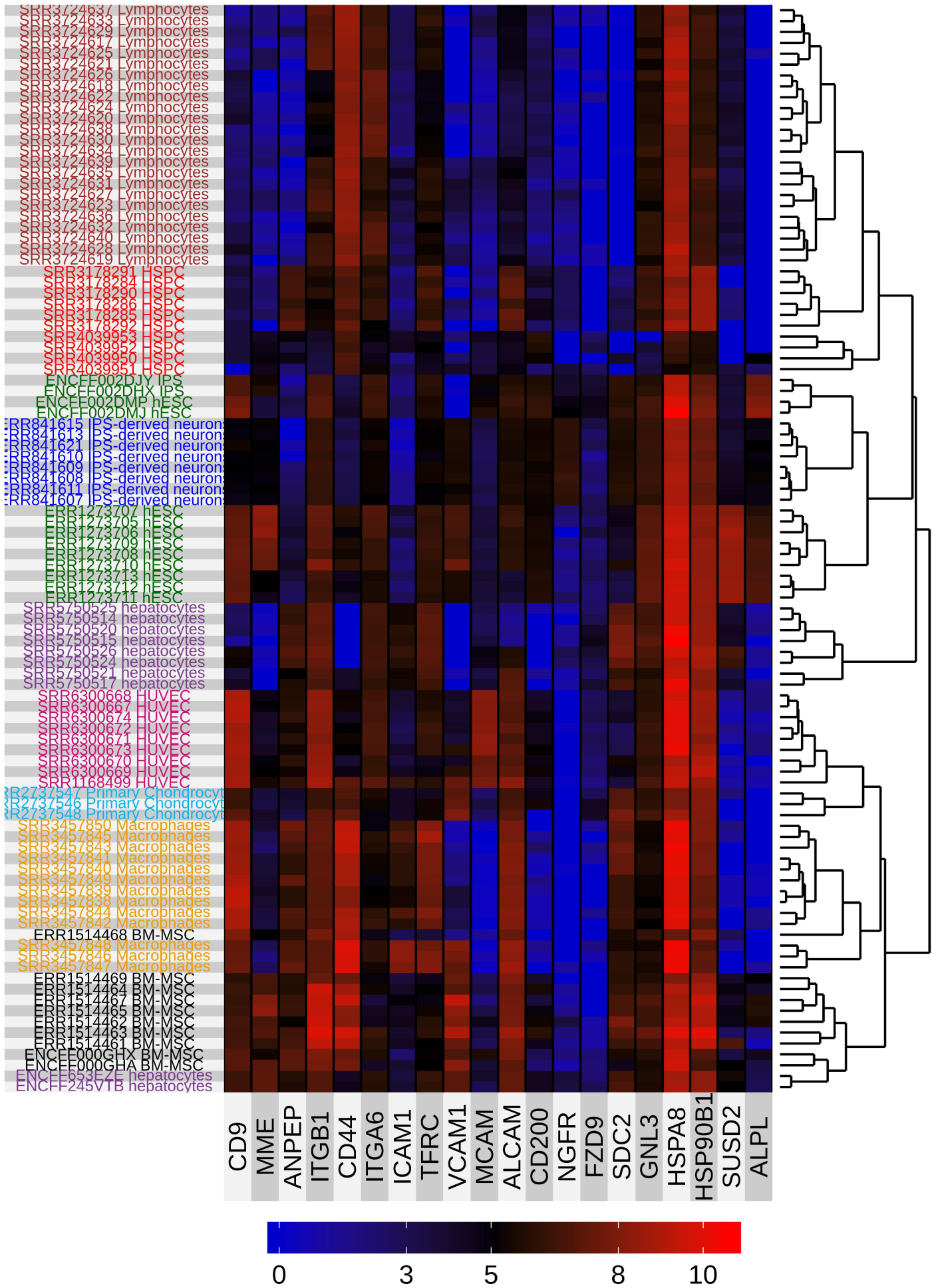
Heatmap present positive markers for MSC proposed in the bibliography

**Supplementary Figure 4:**
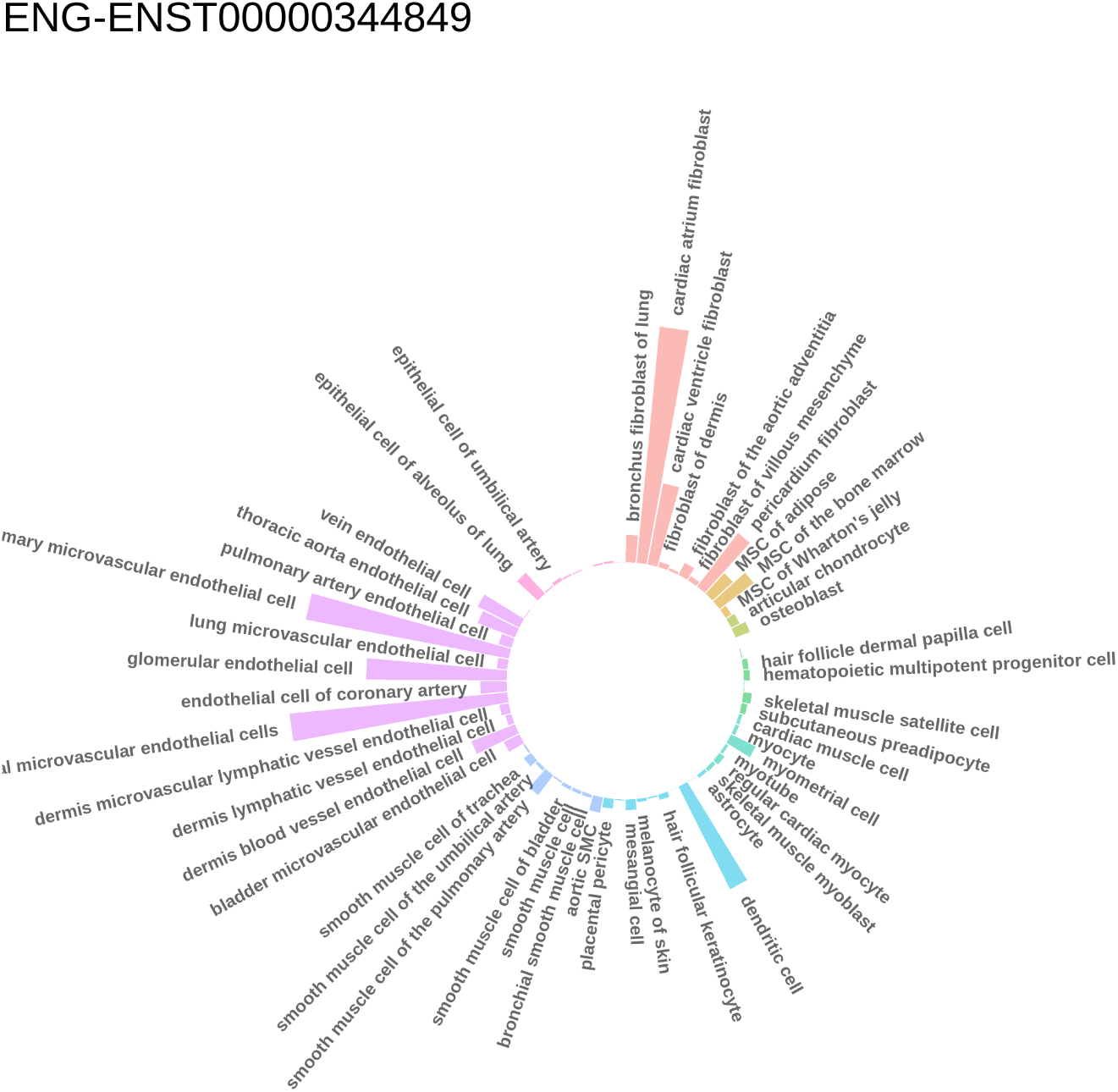
Relative expression of 3 positive markers of ENG (CD105) across ENCODE’s ribodepleted RNAseq datas, made by K-mer quantification, normalized in kmer by million

**Supplementary Figure 5:**
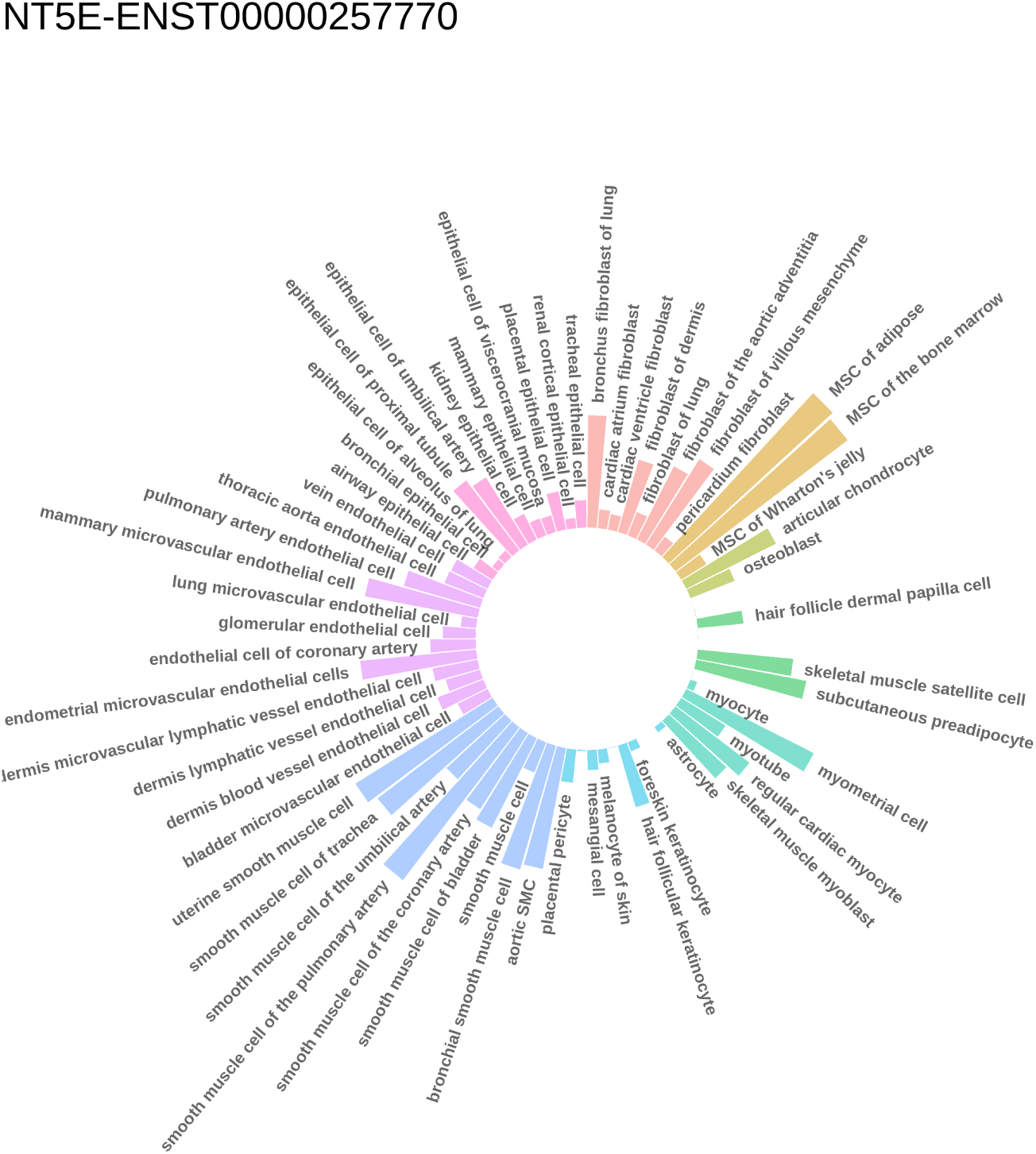
Relative expression of 3 positive markers of NT5E (CD73) across ENCODE’s ribodepleted RNAseq datas, made by K-mer quantification, normalized in kmer by million

**Supplementary Figure 6:**
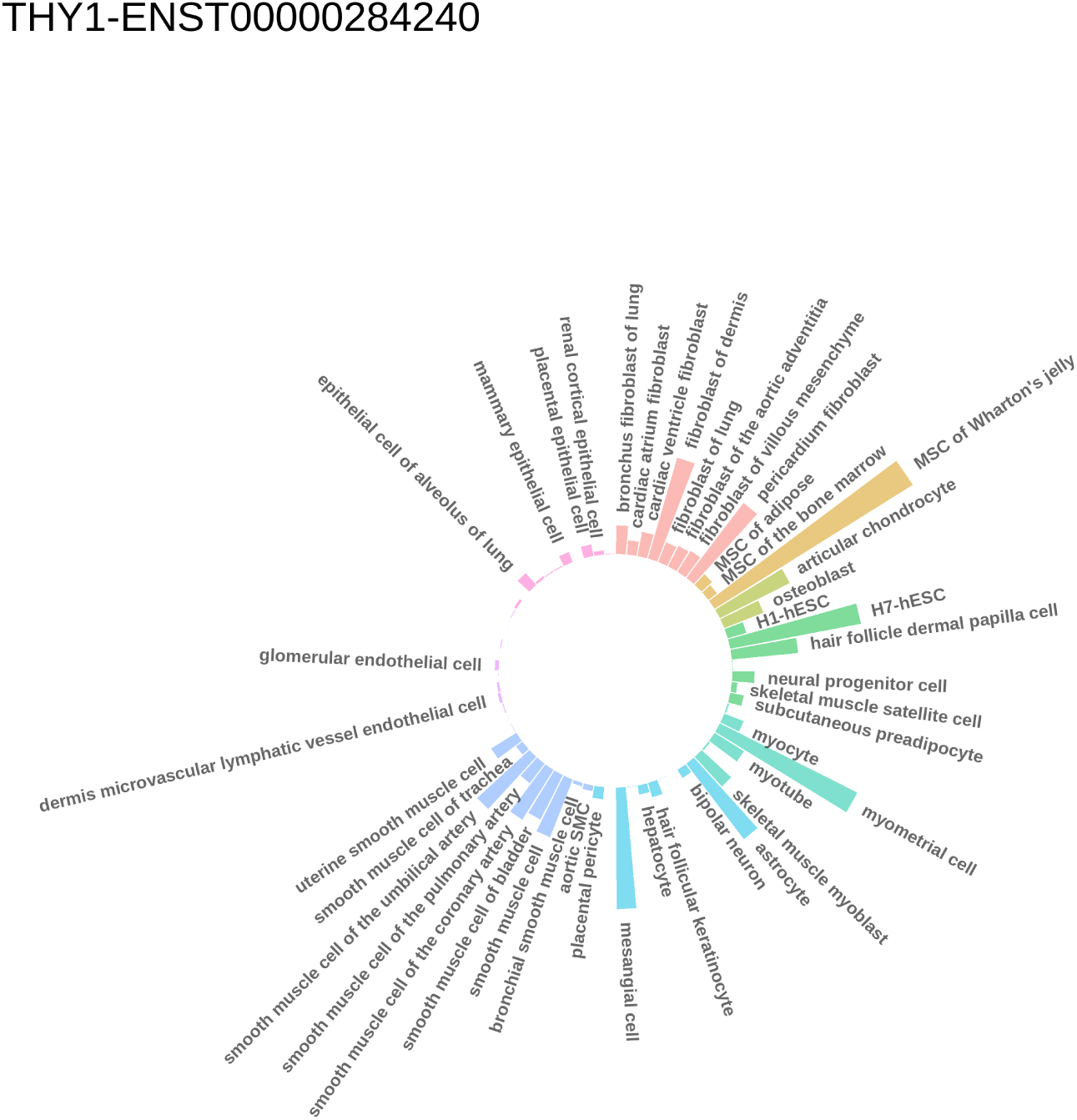
Relative expression of 3 positive markers of THY1 (CD90) across ENCODE ribodepleted RNAseq datas, made by K-mer quantification, normalized in kmer by million

**Supplementary Figure 7:**
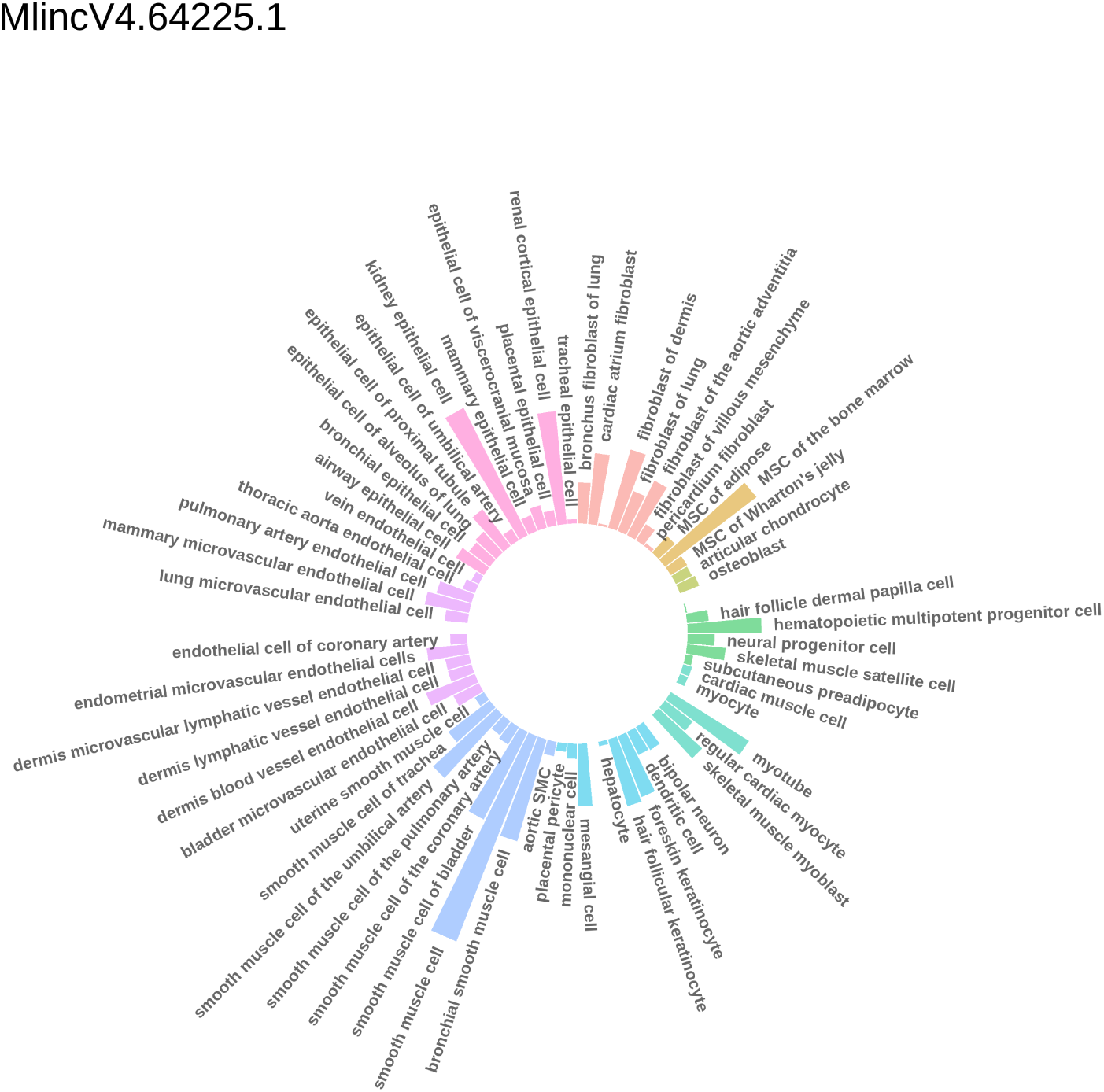
Relative expression of 3 positive markers of Mlinc.64225.1 across ENCODE’s ribodepleted RNAseq datas, made by K-mer quantification, normalized in kmer by million

**Supplementary Figure 8:**
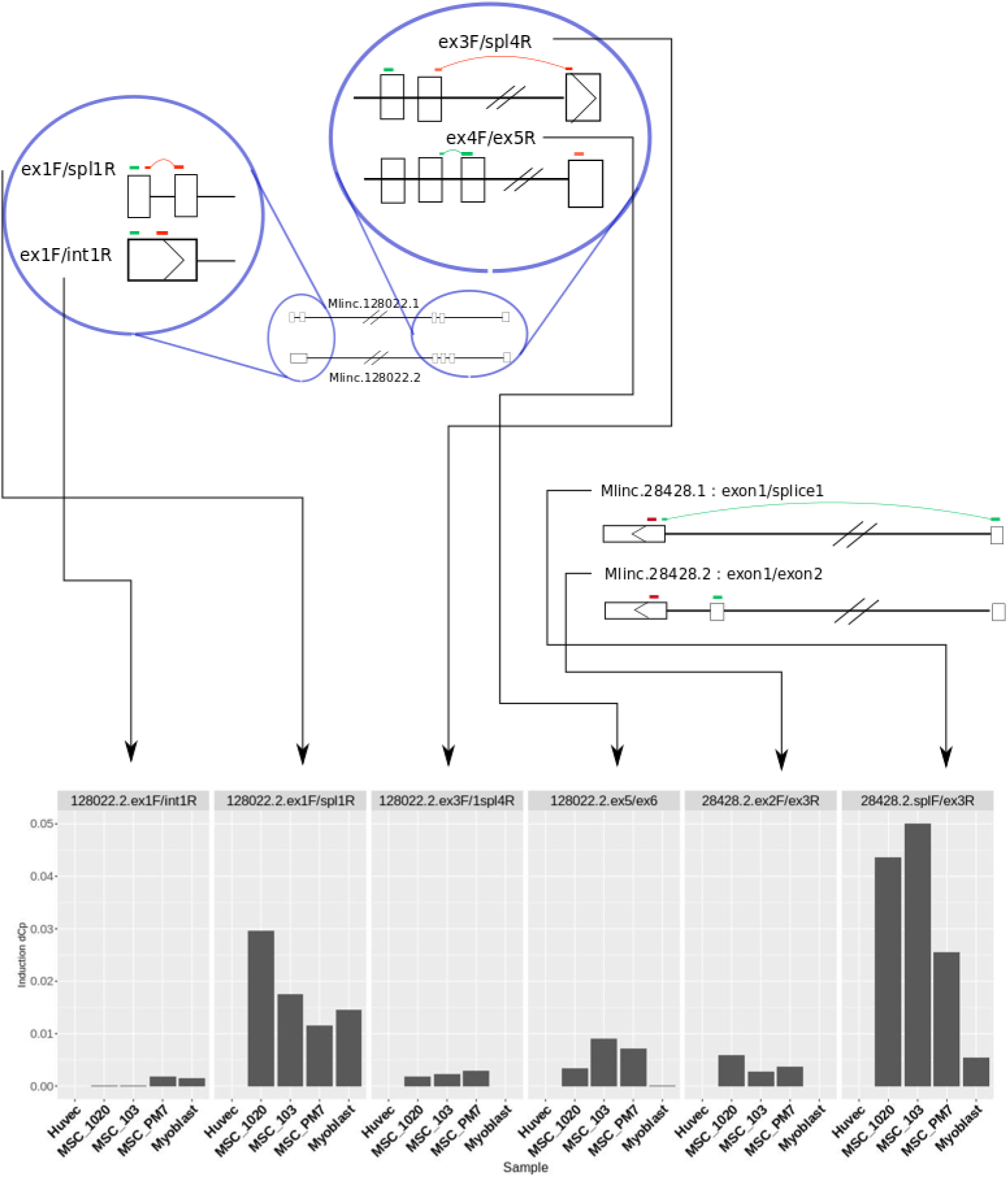
Primer position on selected Mlinc candidates and corresponding expression in MSCs, HUVECs and Myoblasts

**Supplementary Figure 9:**
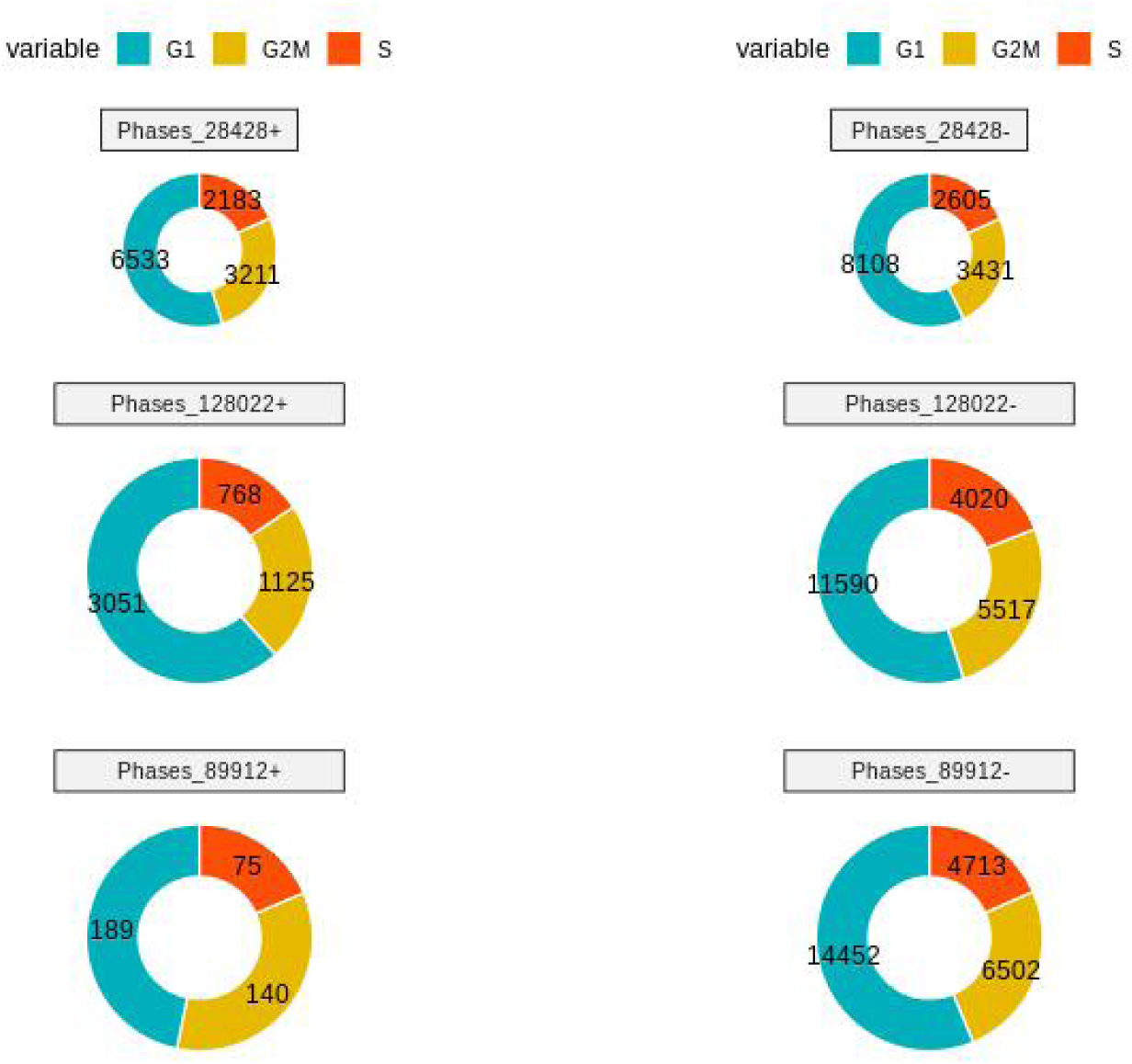
cell cycle and single cell.

**Table_S1.**
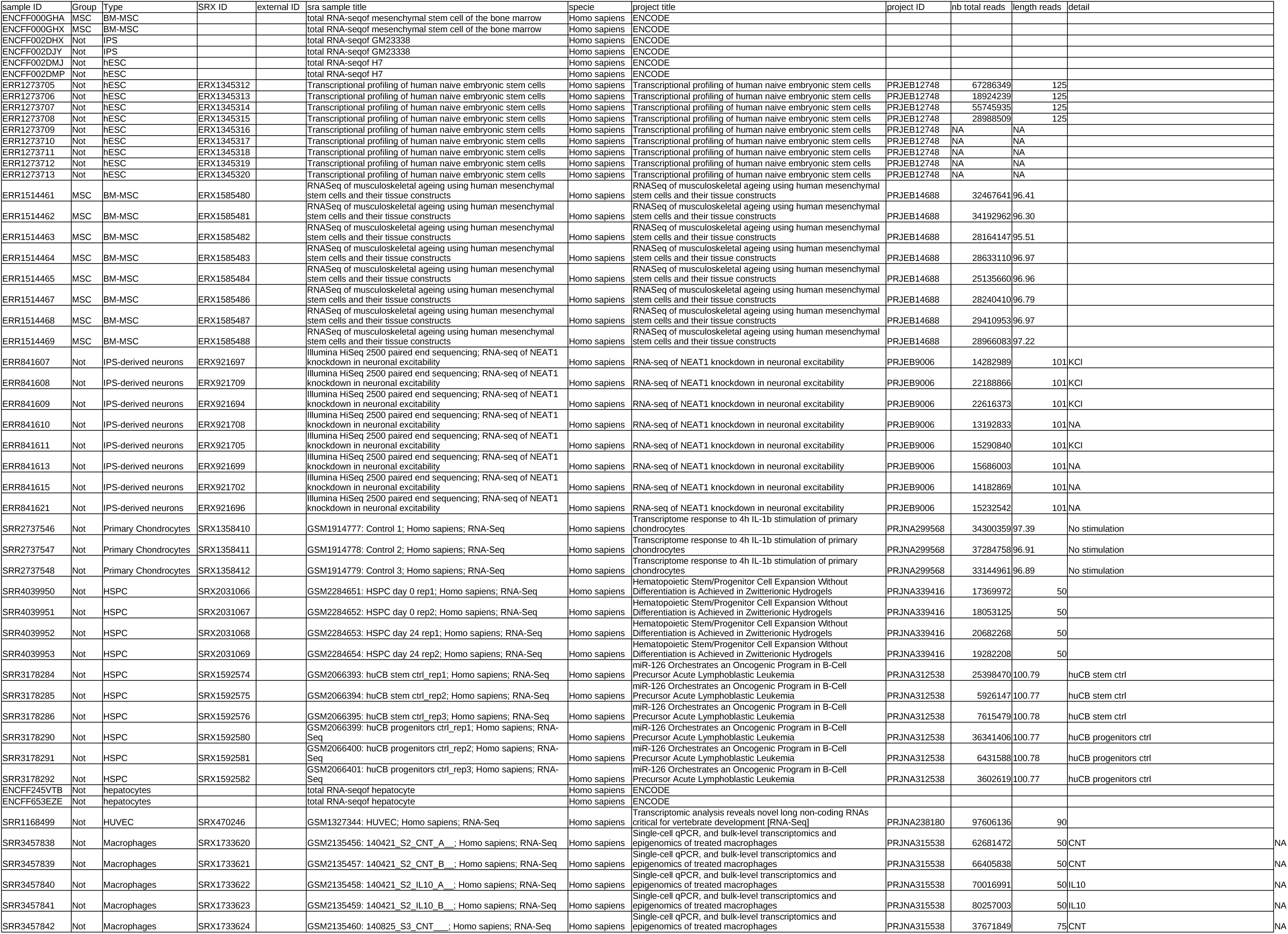
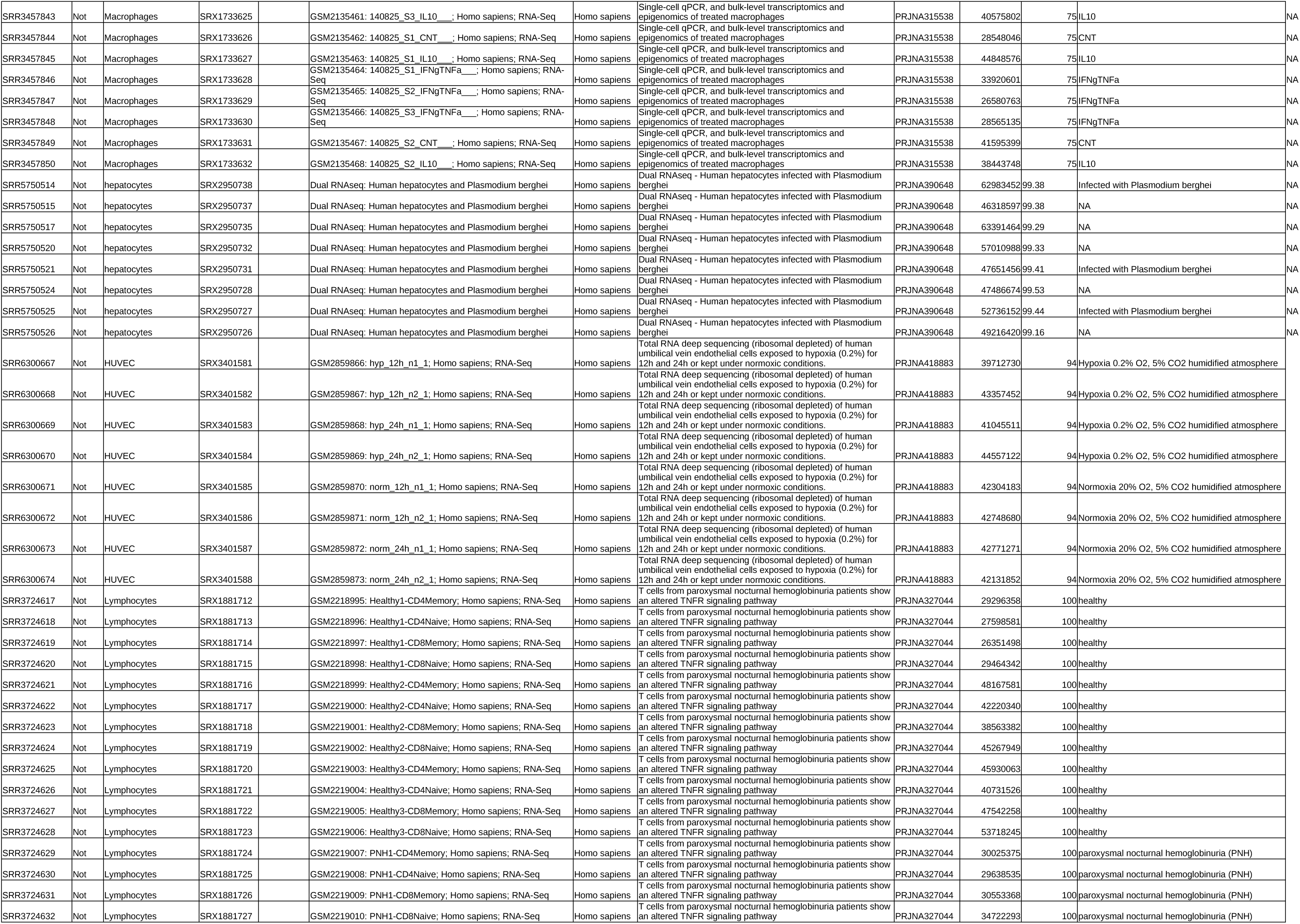
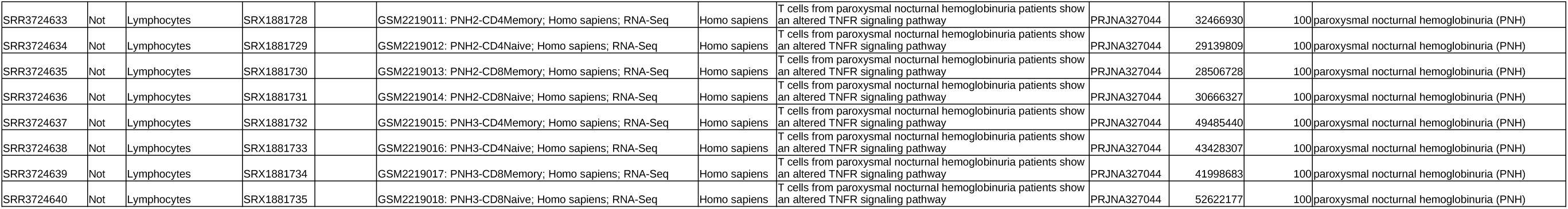
Metadata for differential analysis

**Table_S2.**
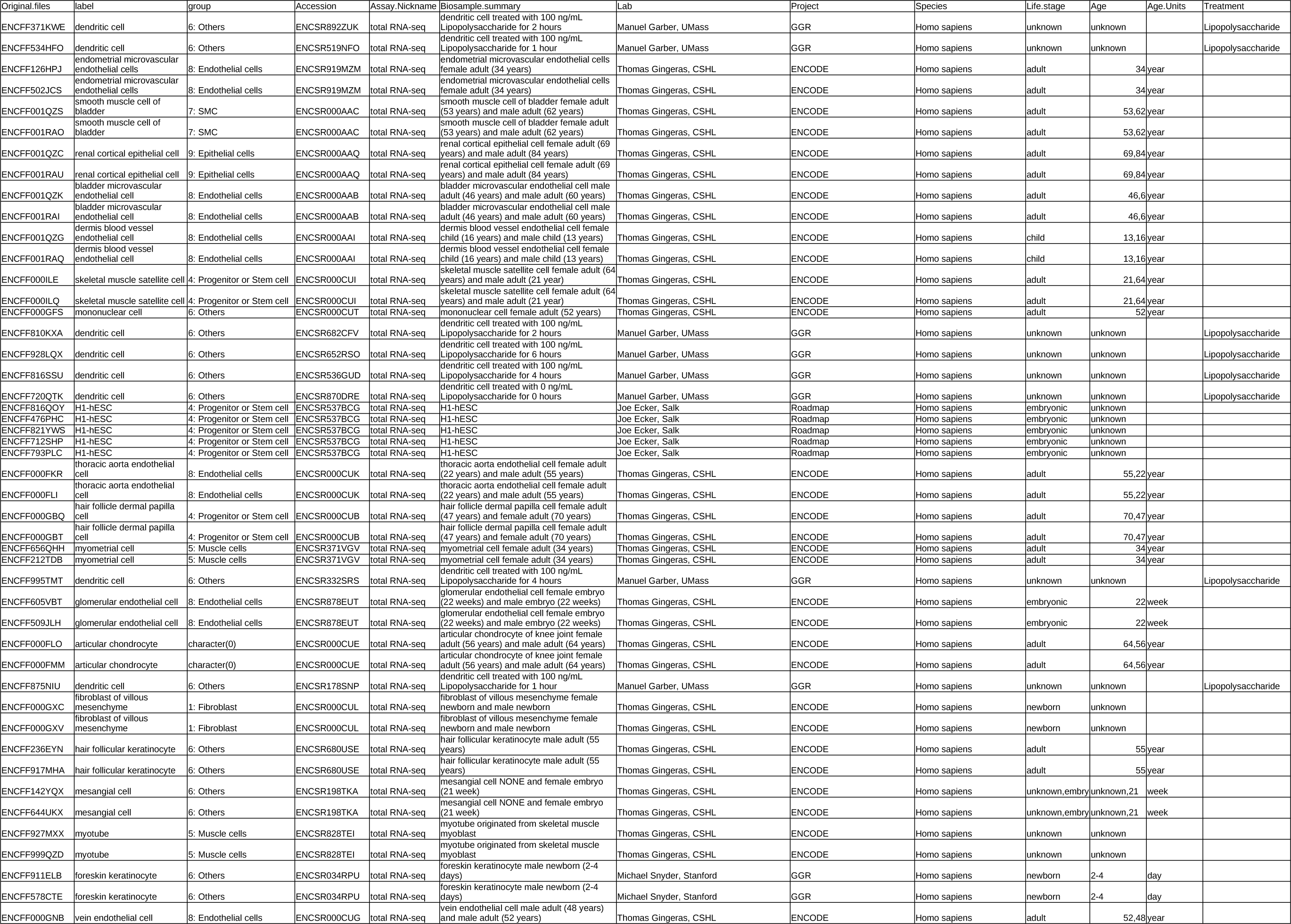
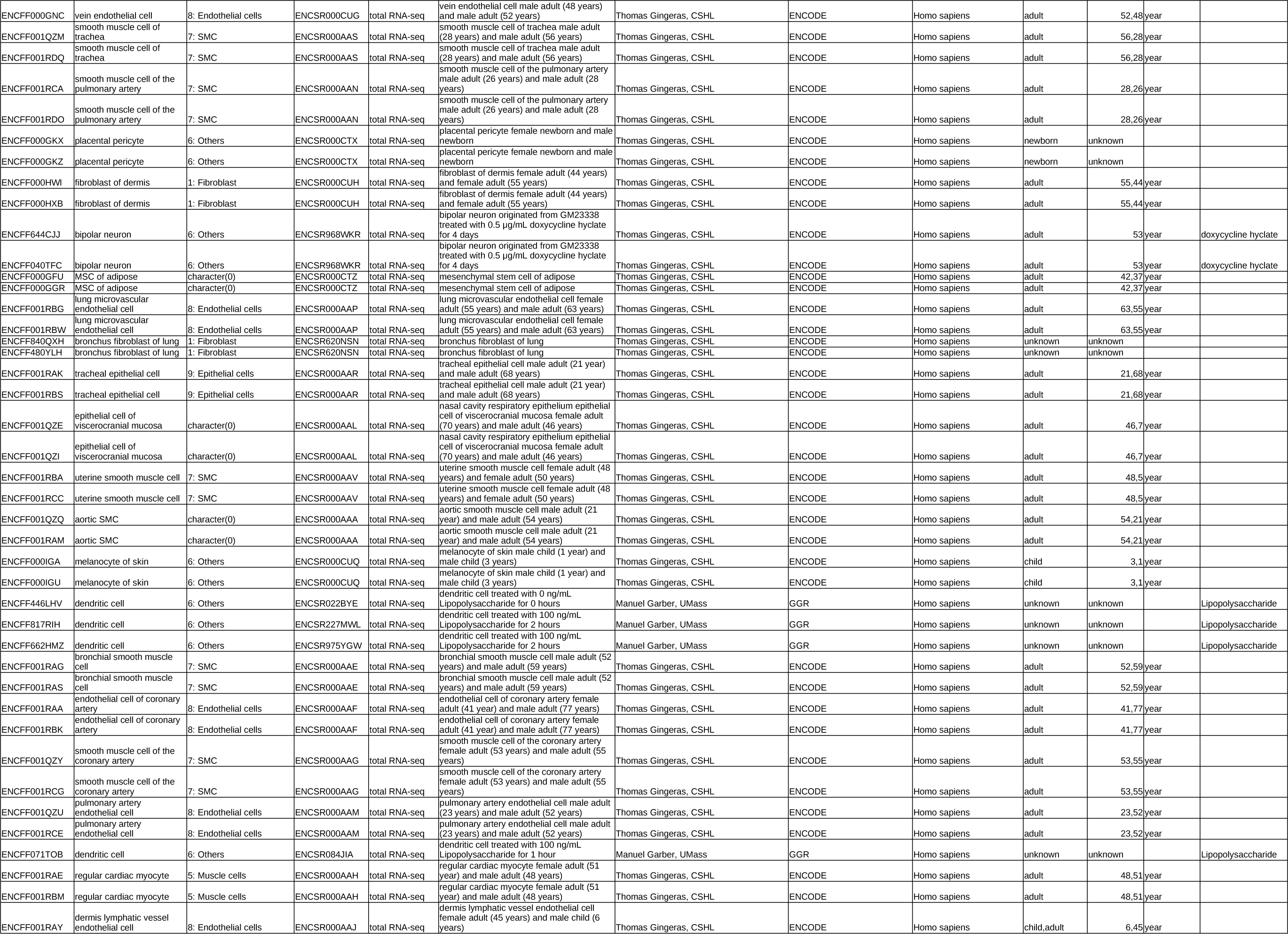
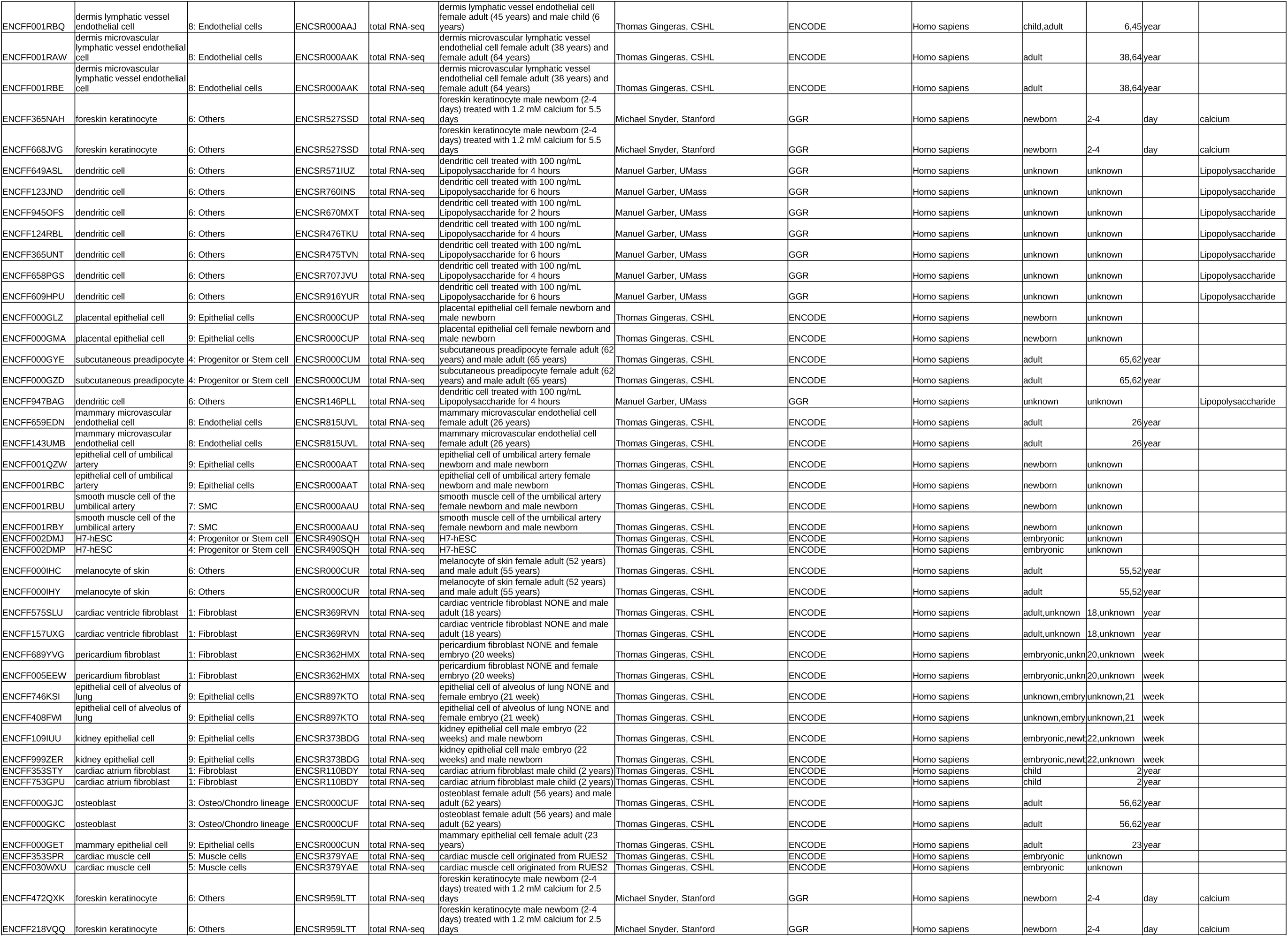
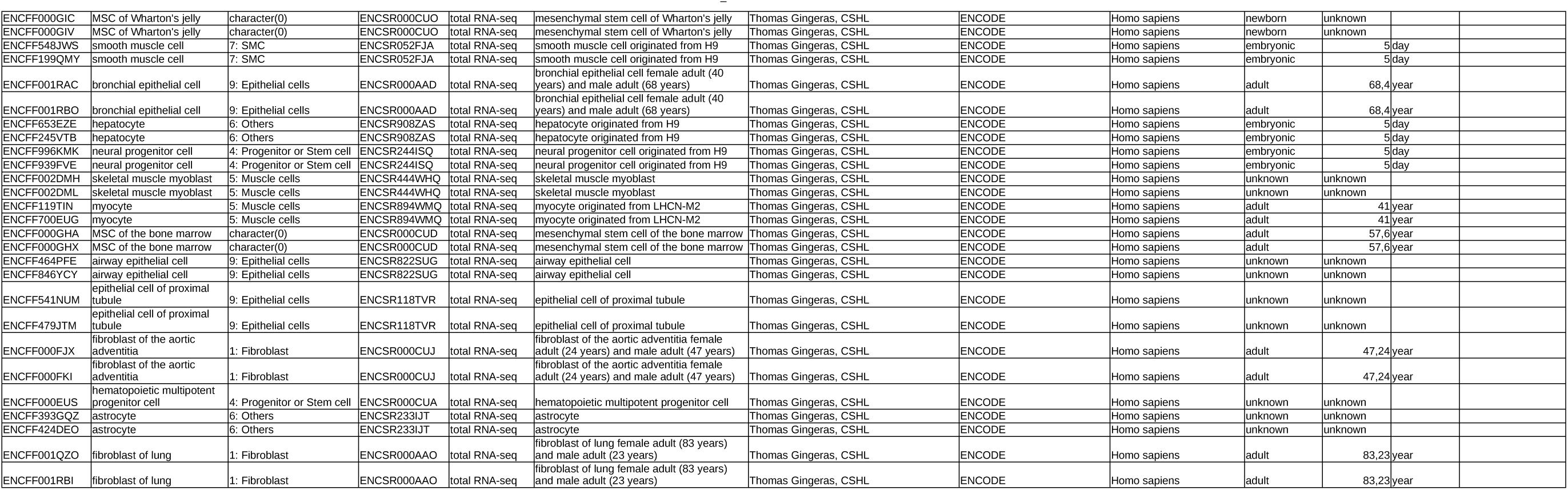
Metadata for ENCODE kmer research

**Table_S3.**
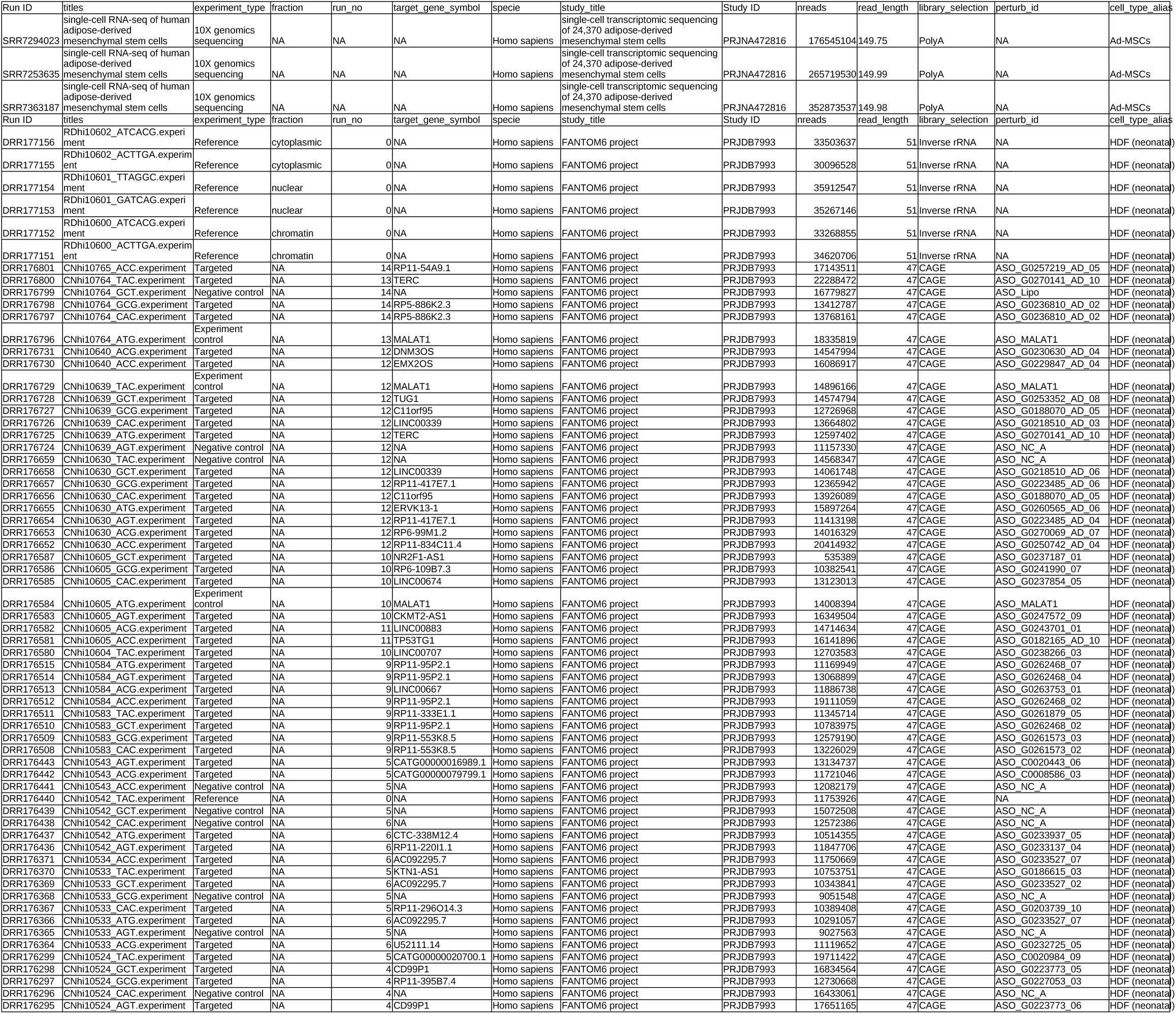
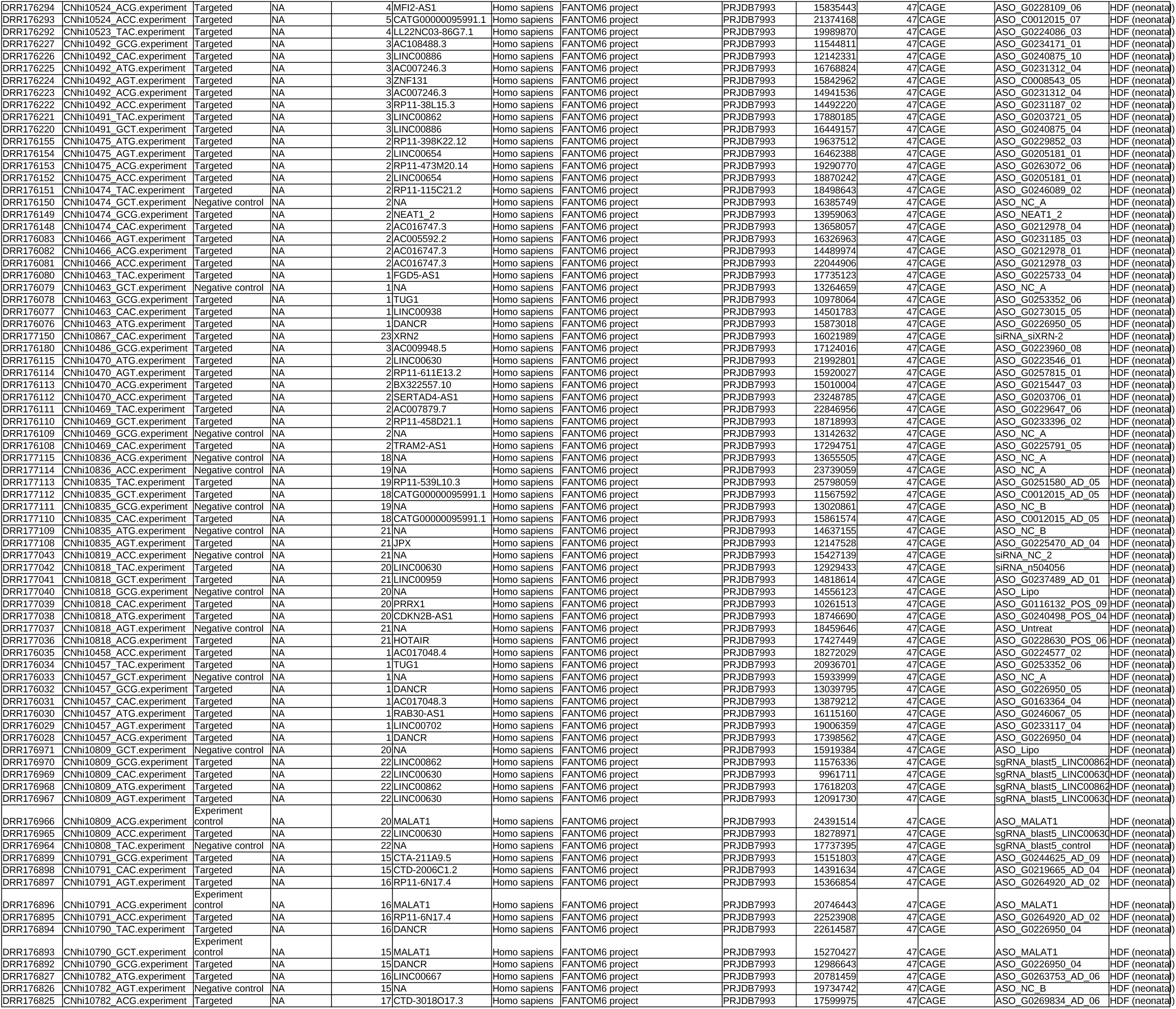
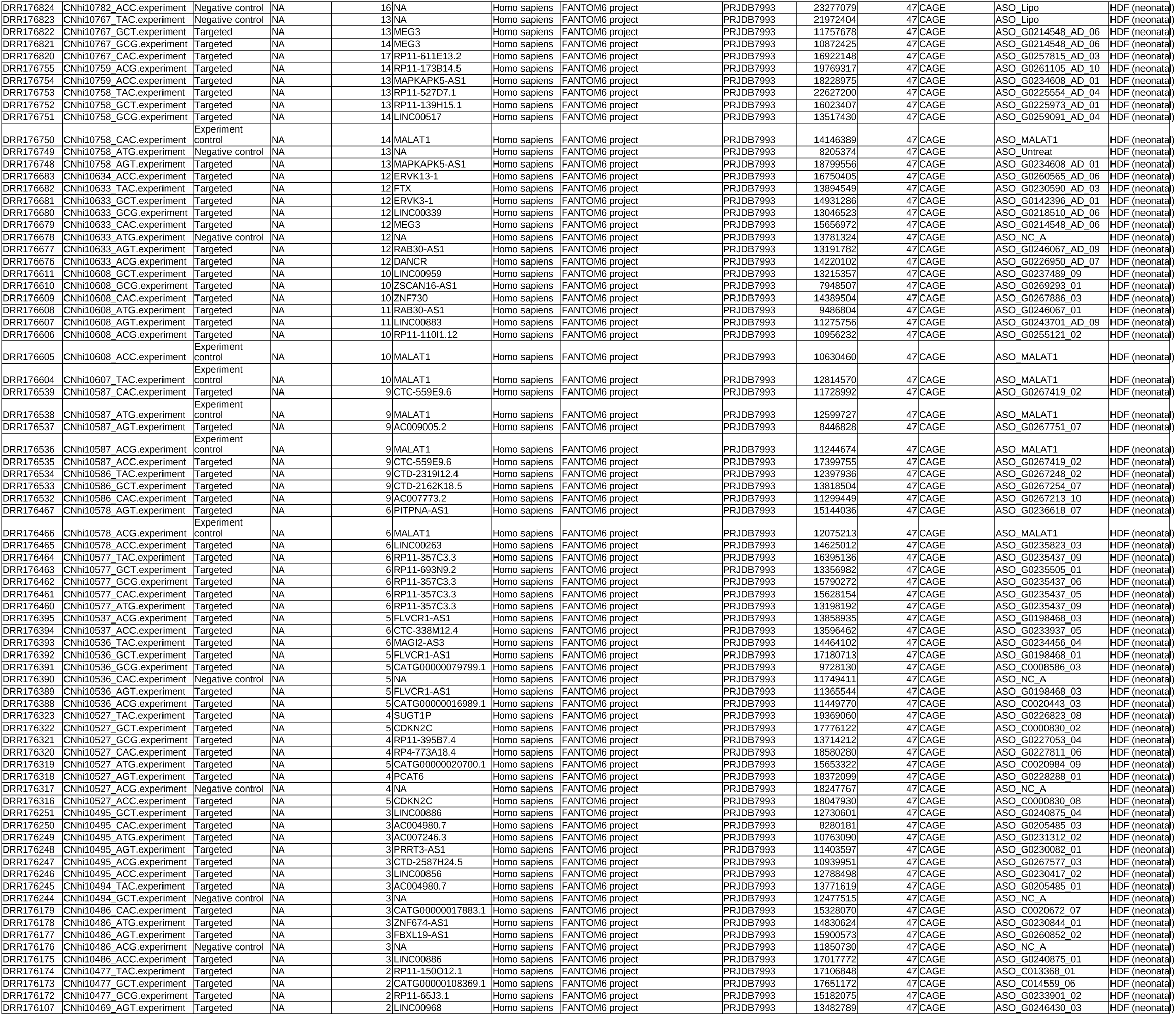
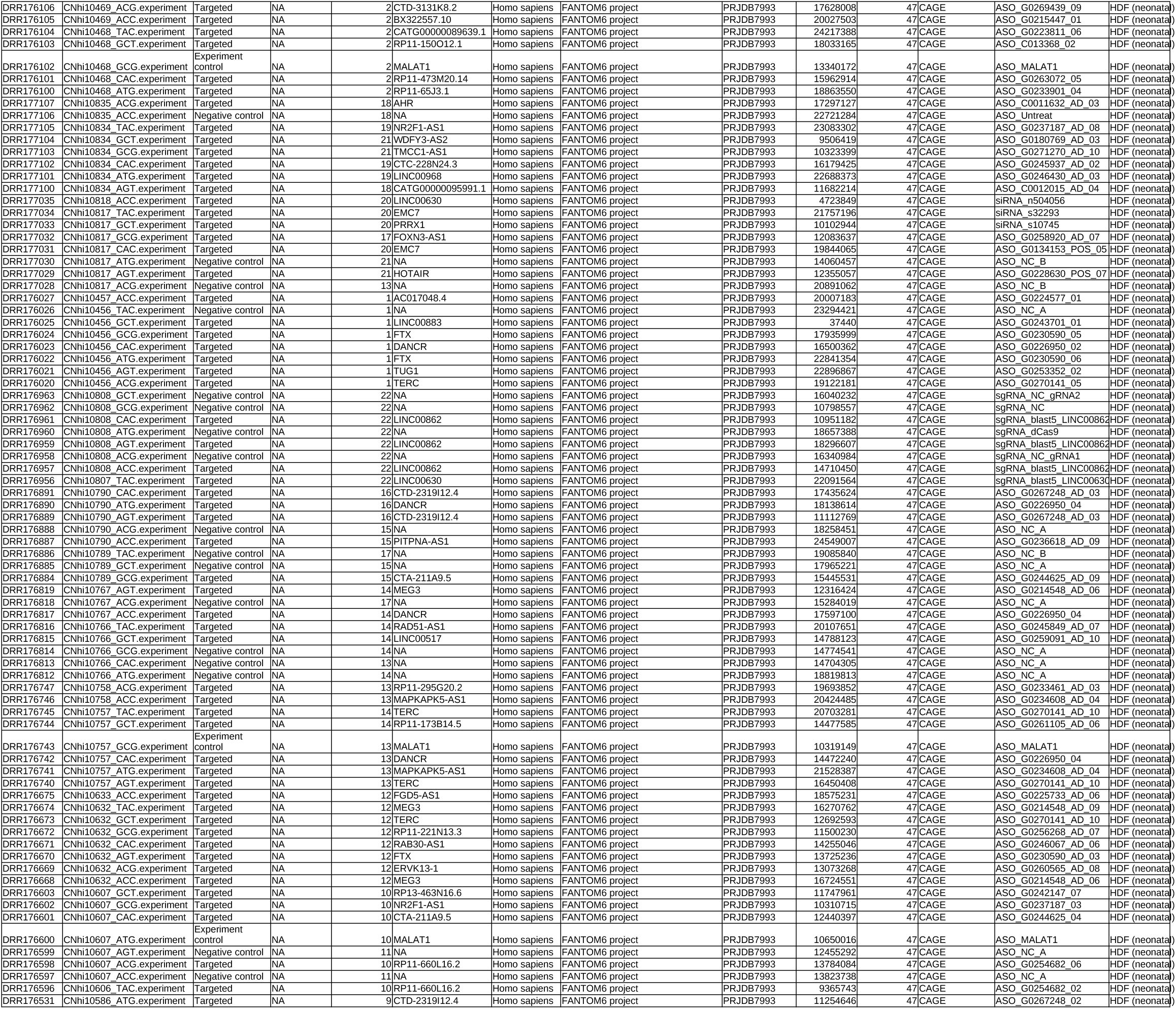
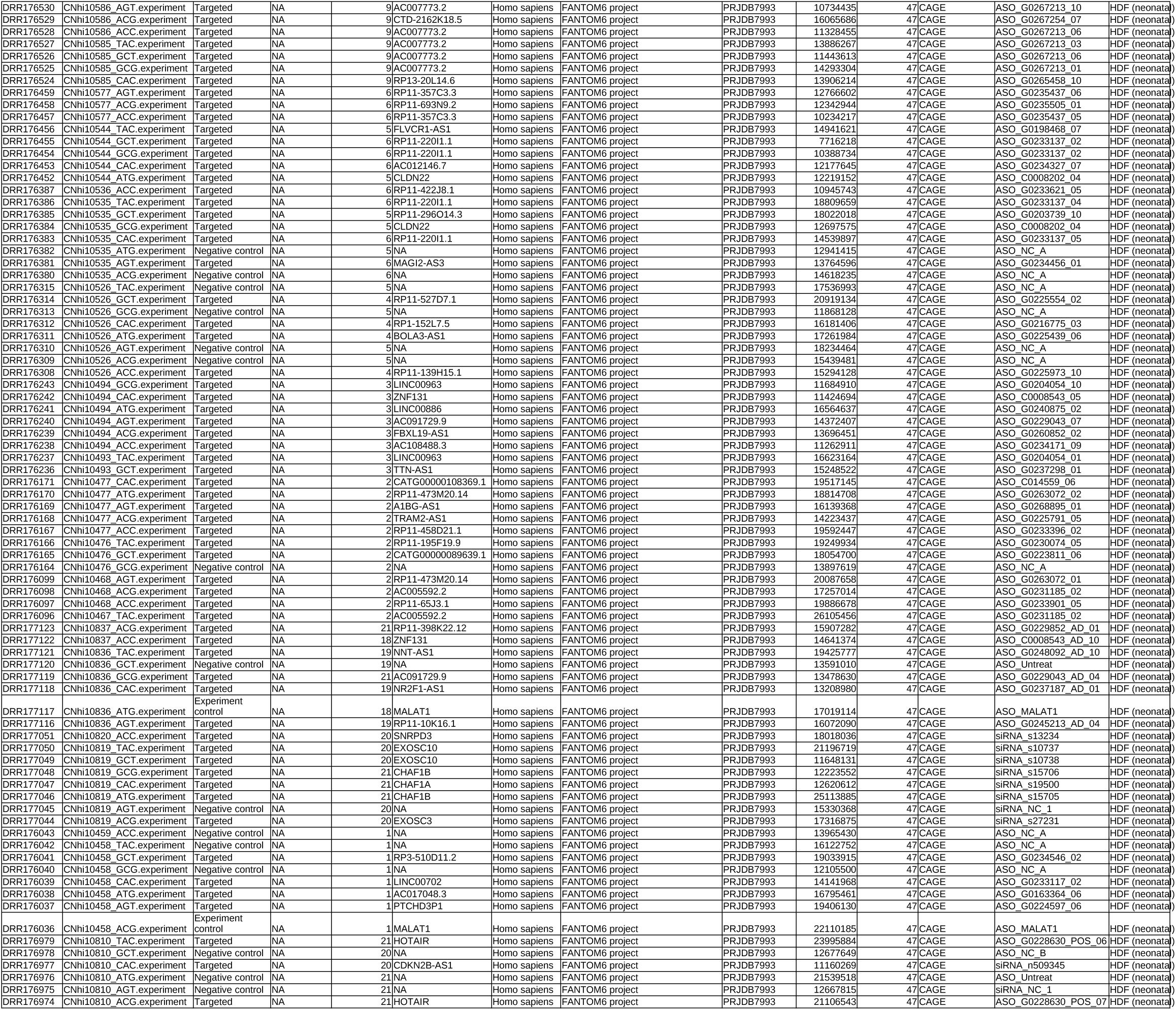
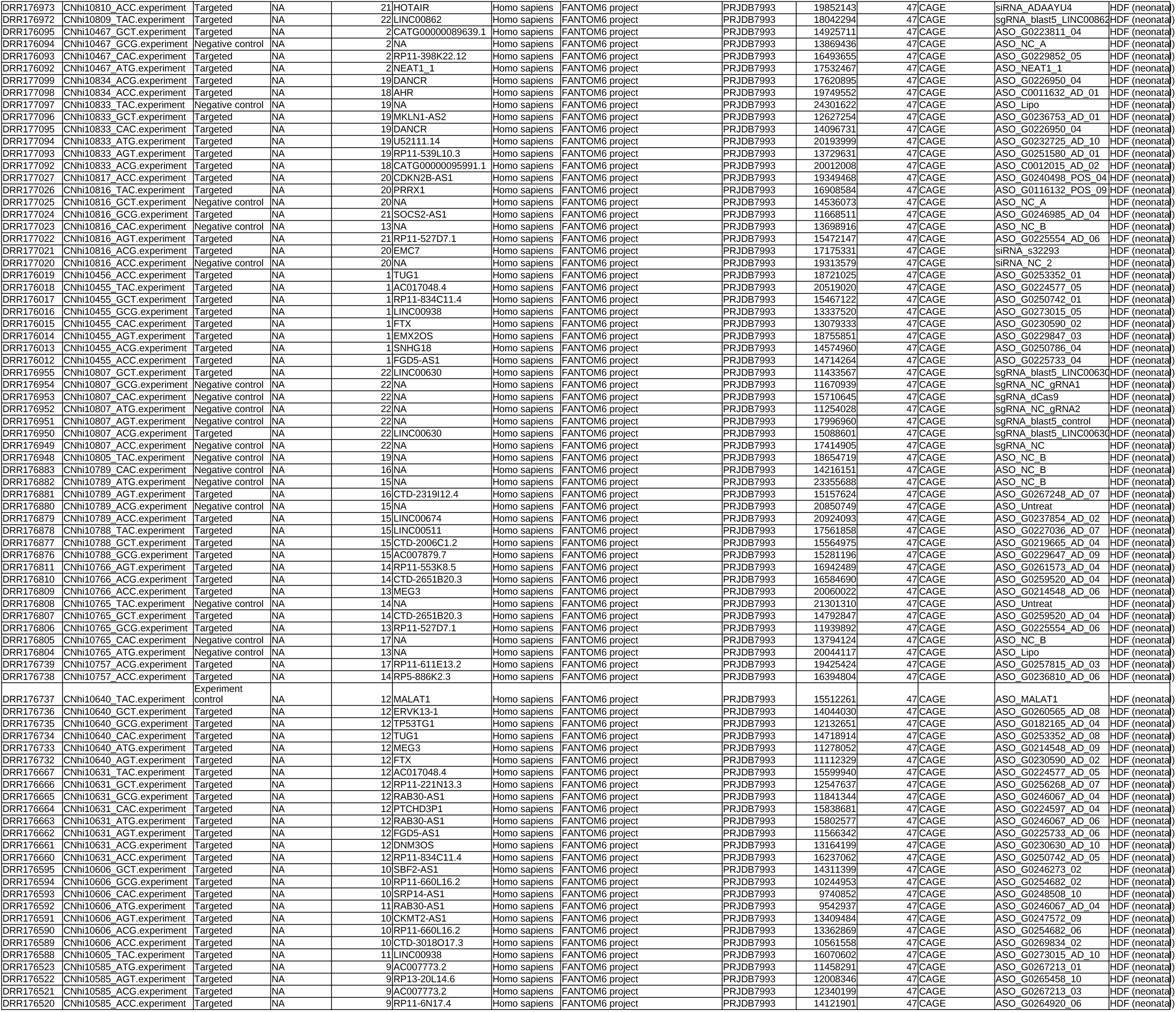
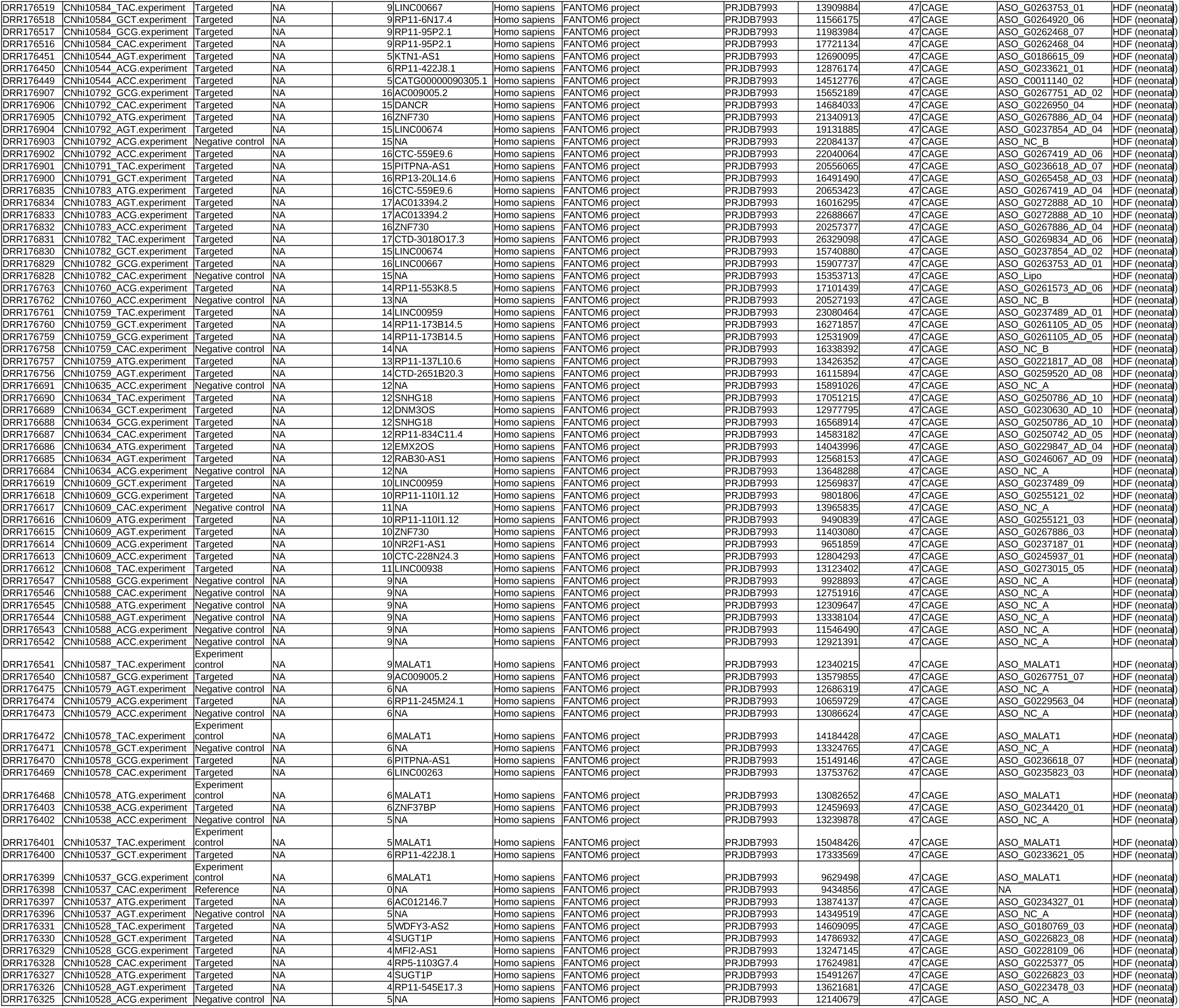
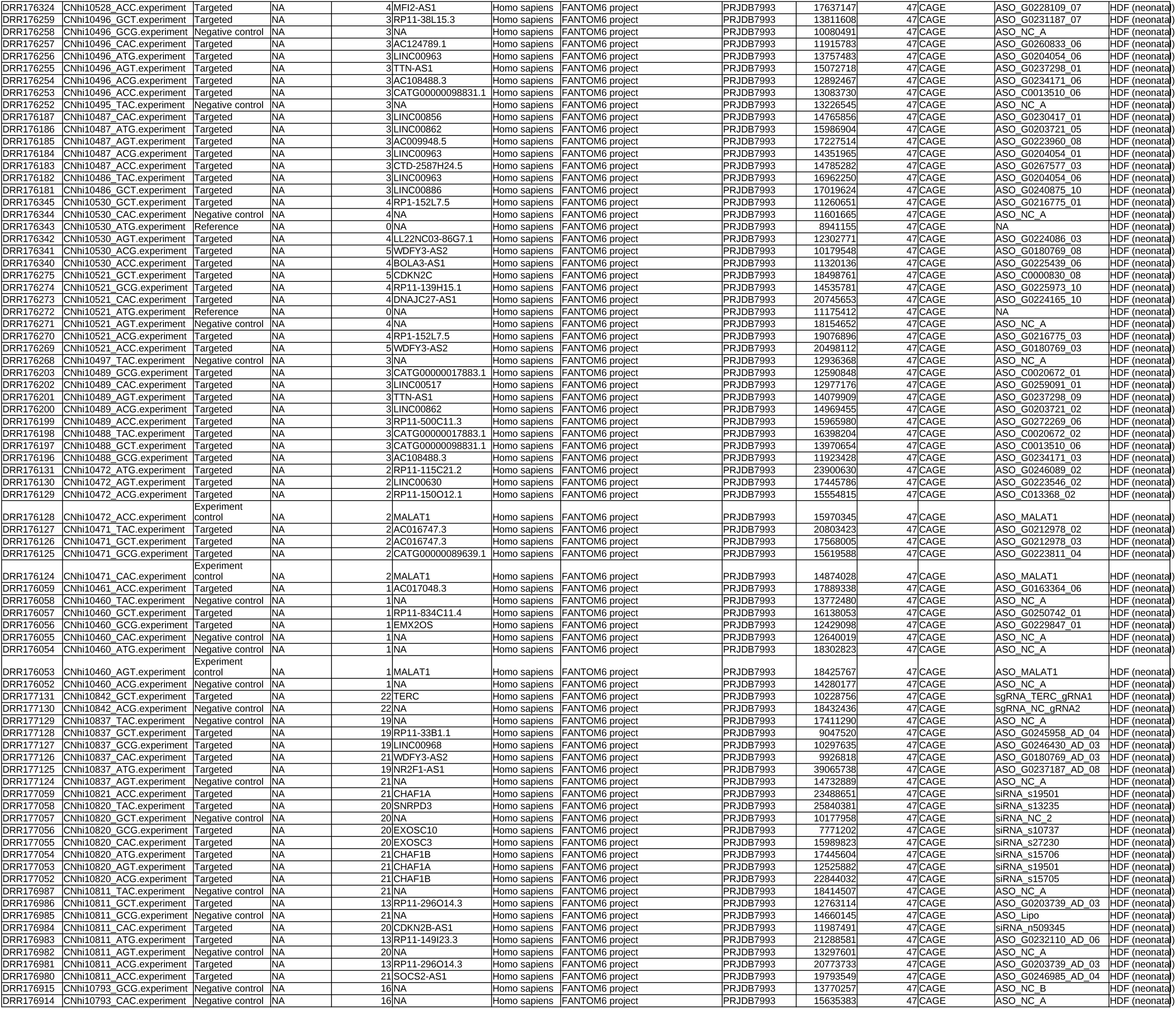
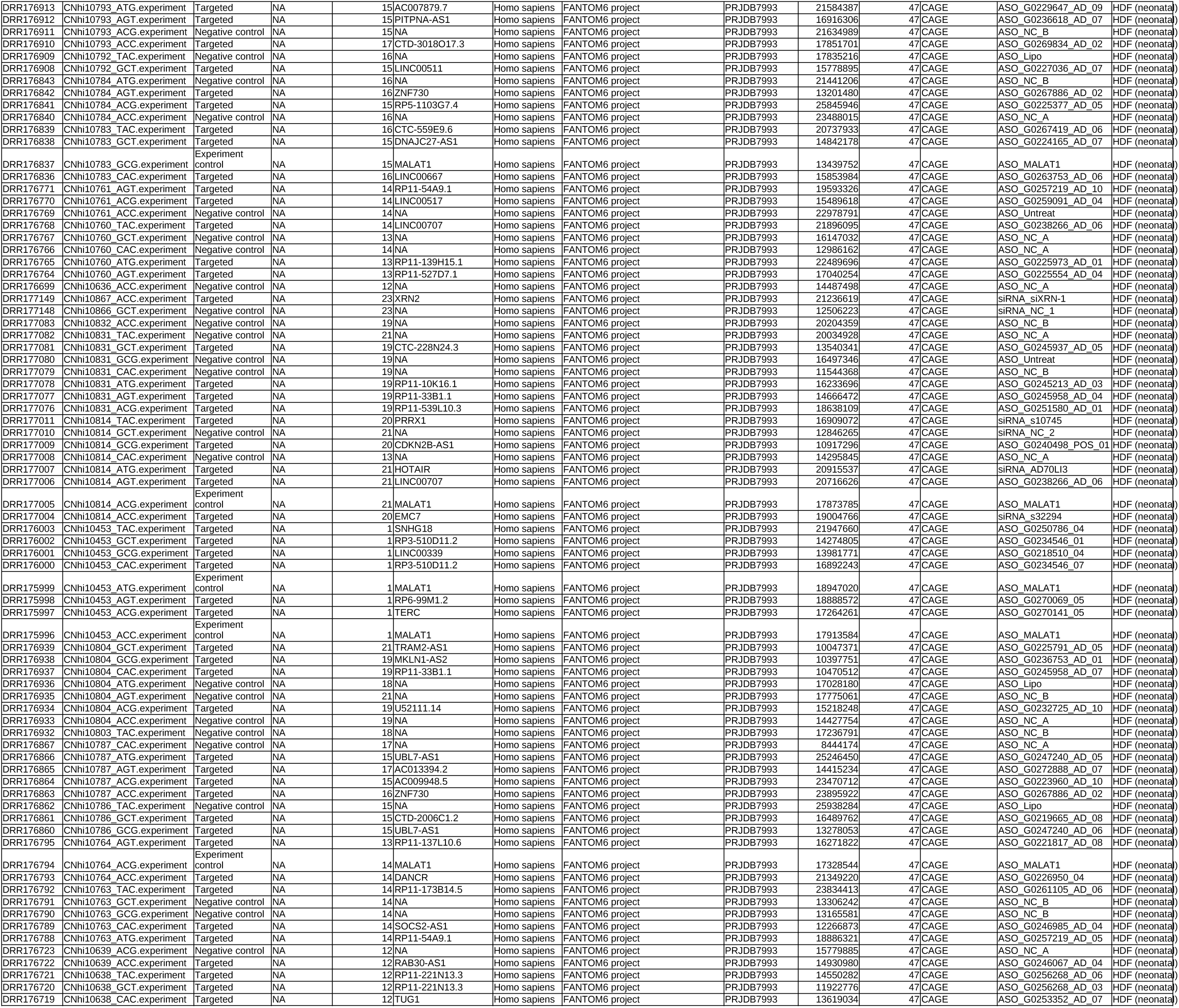
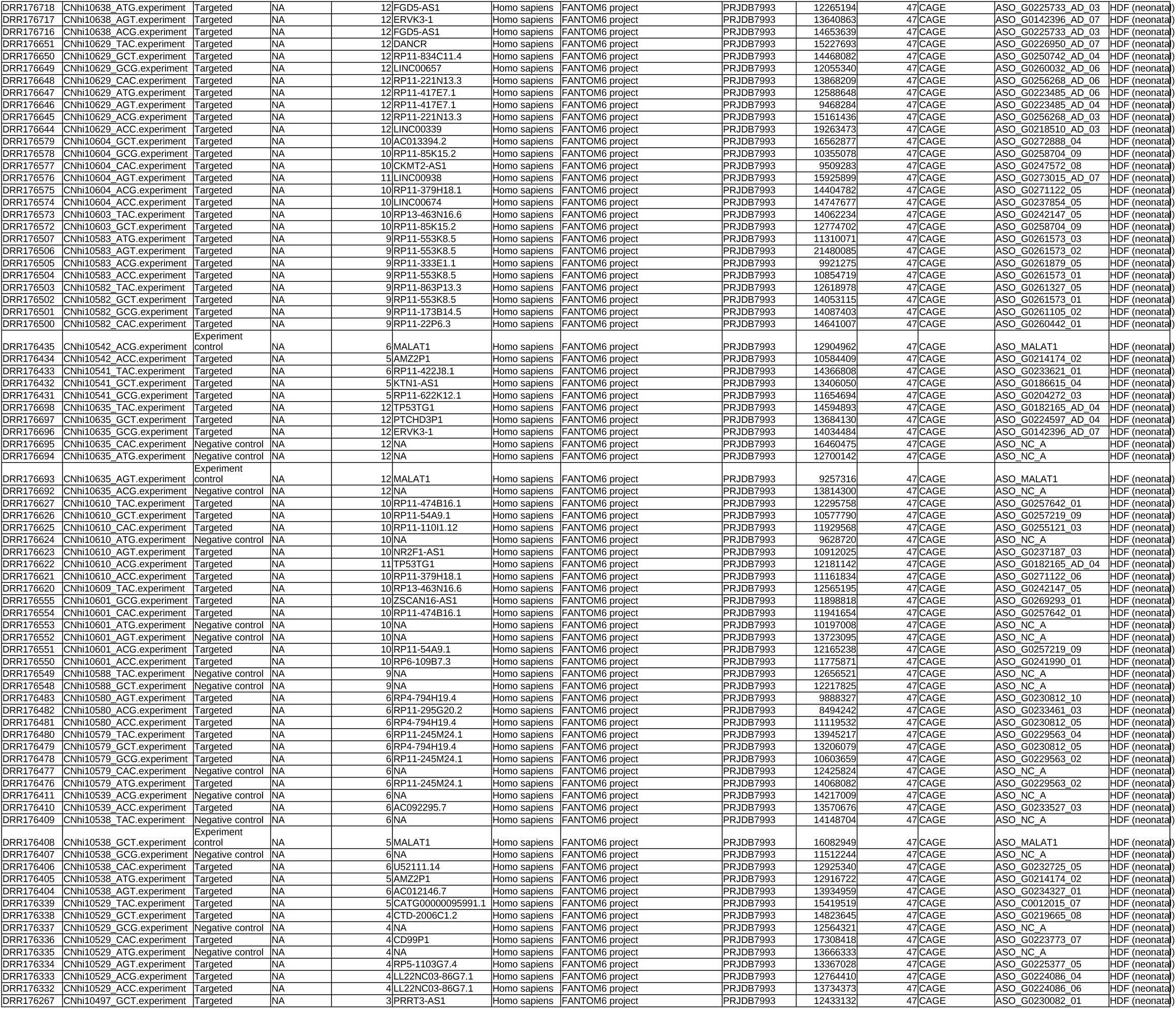
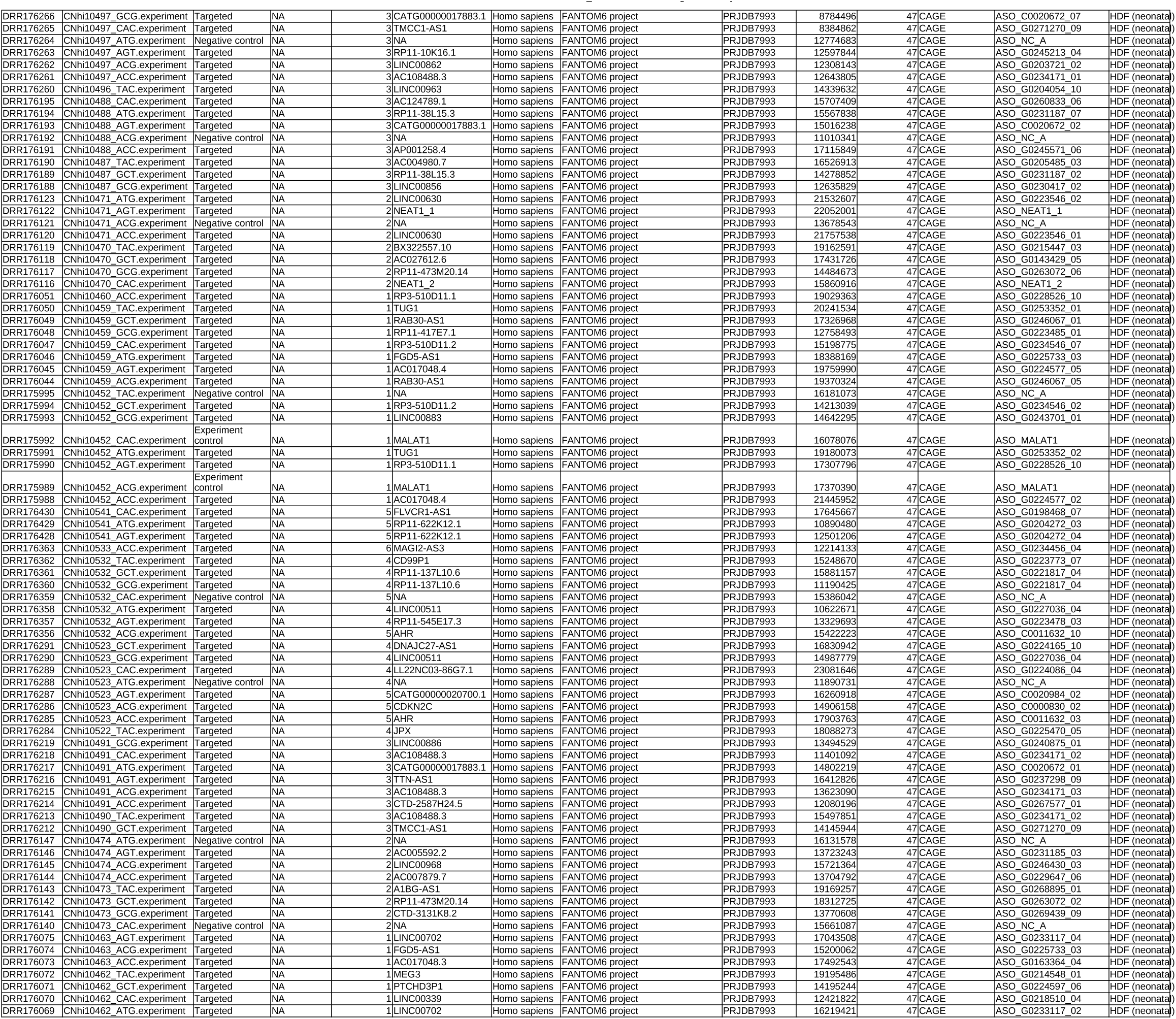
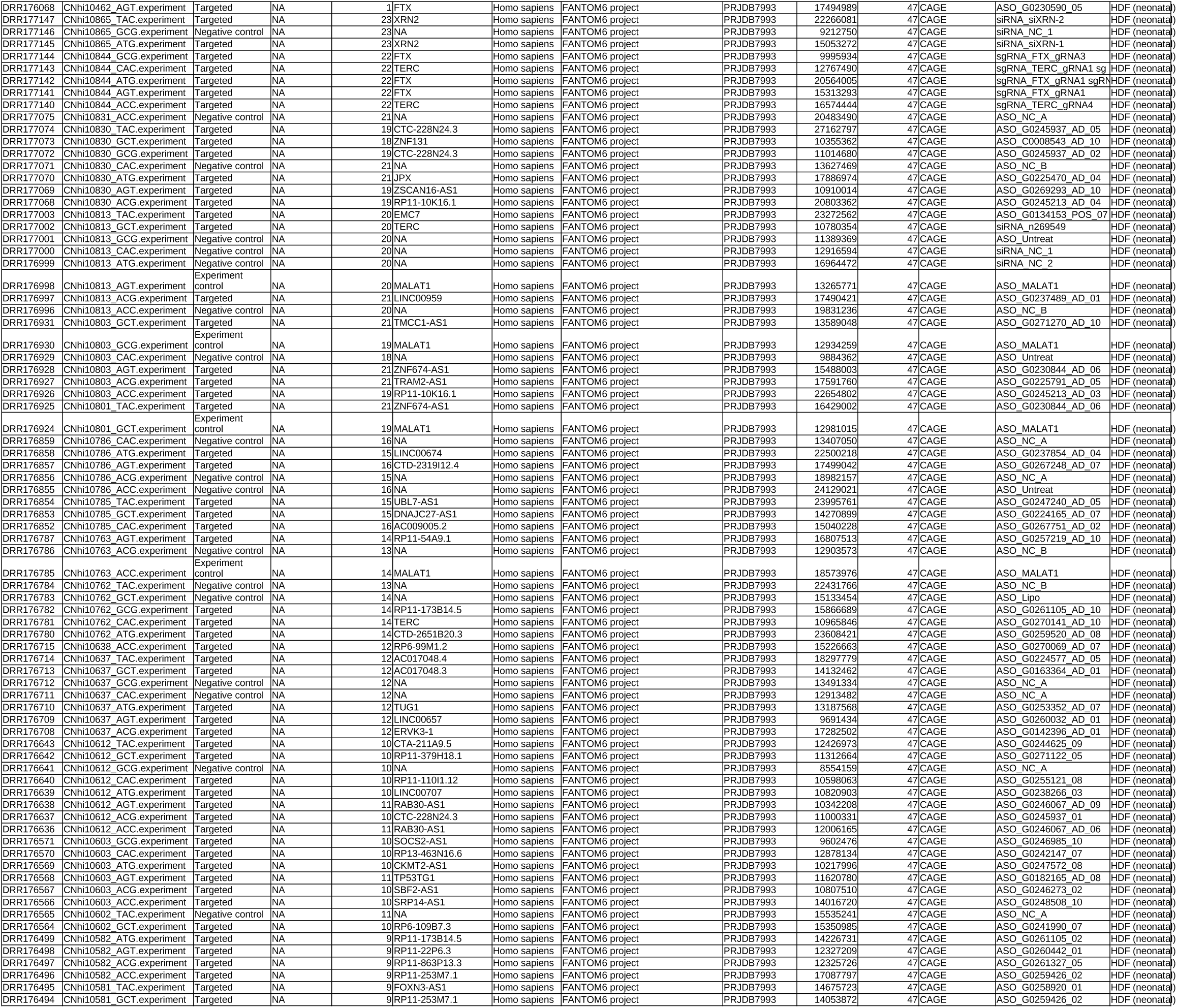
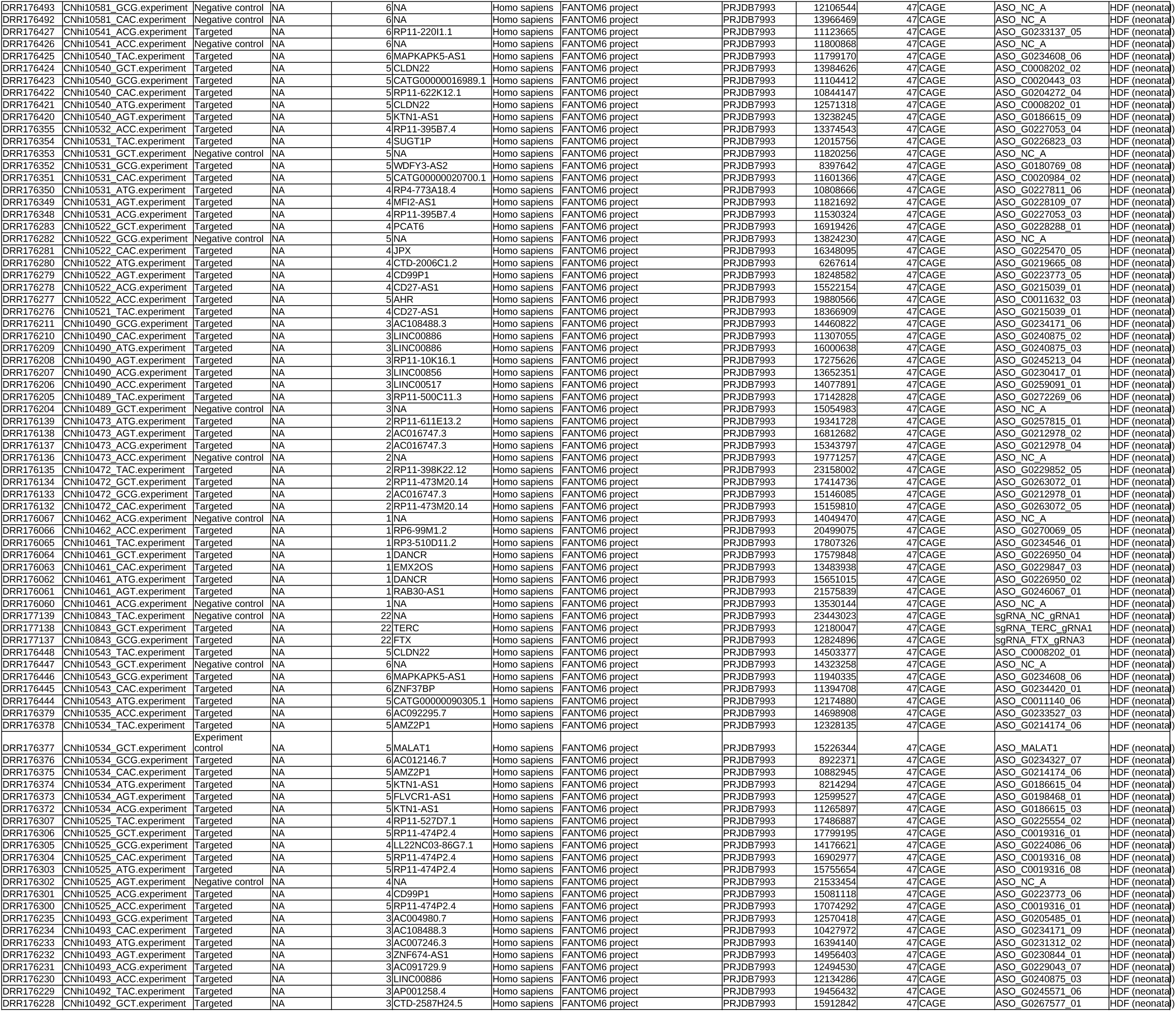
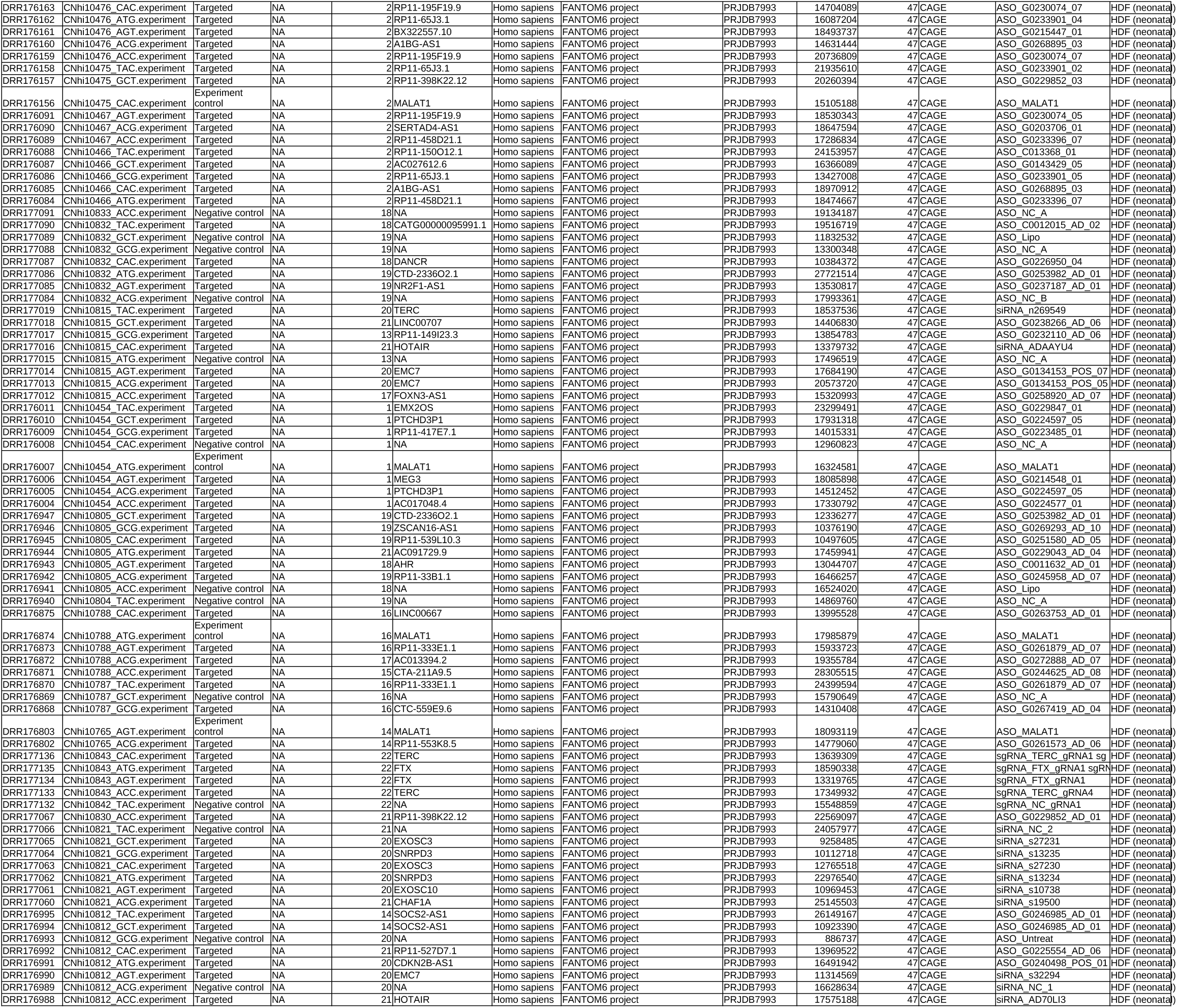
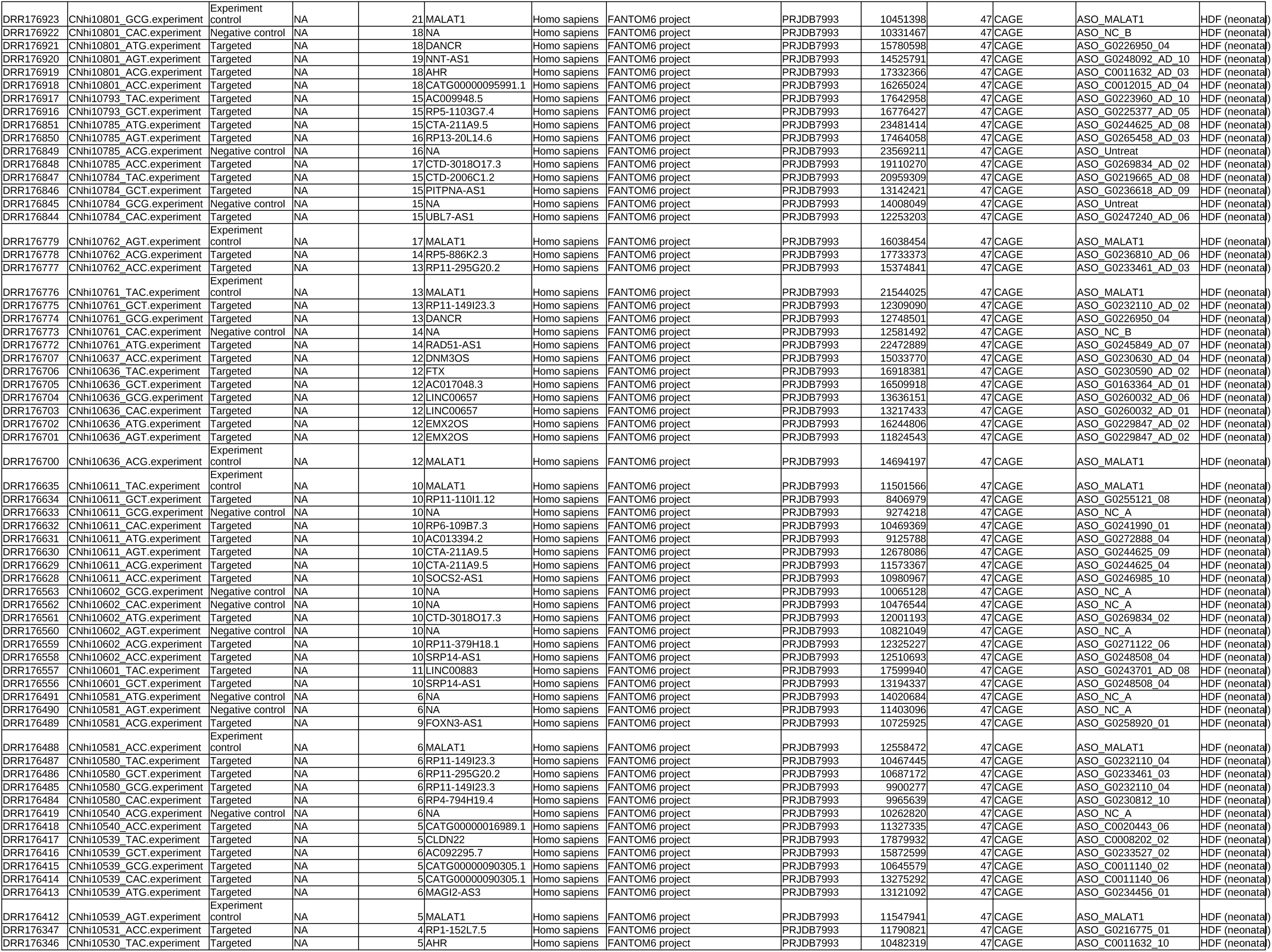
Meatadata for single cell analysis and FANTOM6 kmer research

**Table_S4.**
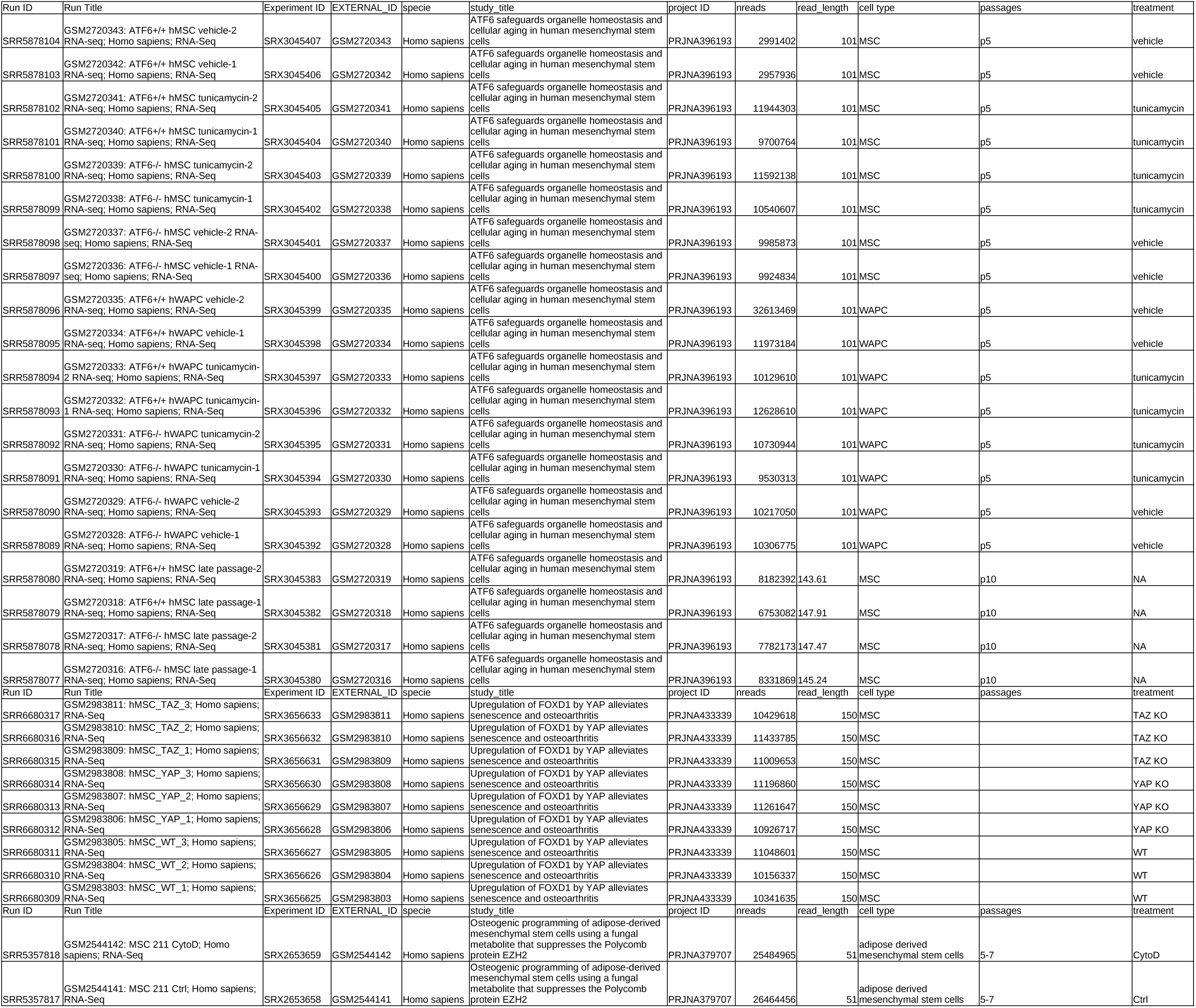
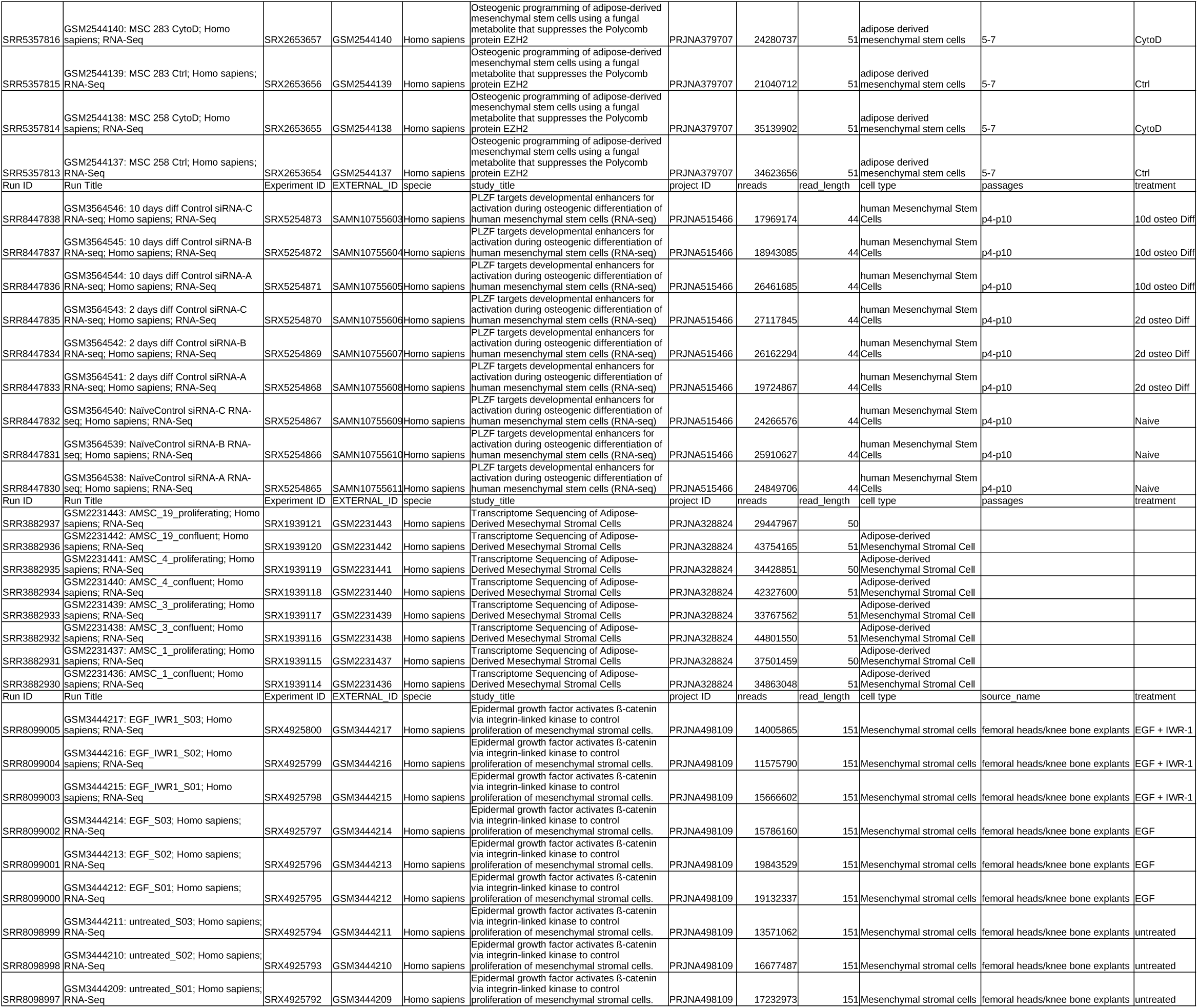
metadata for kmer research in MSC in different conditions

**Table_S5.**
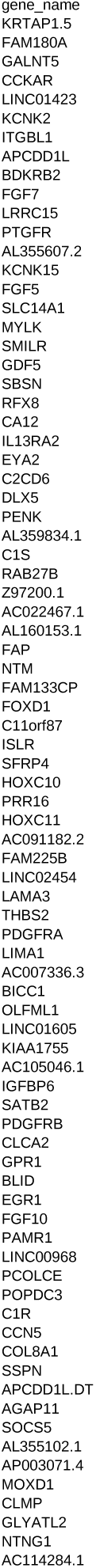
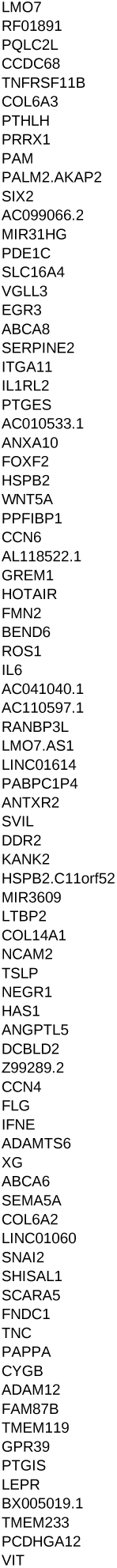
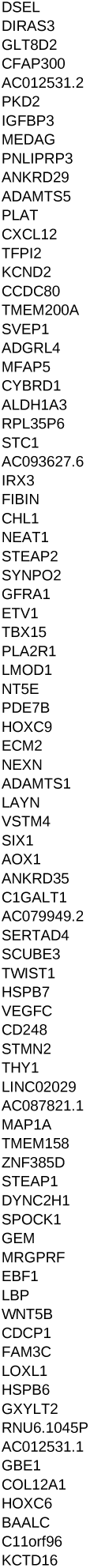
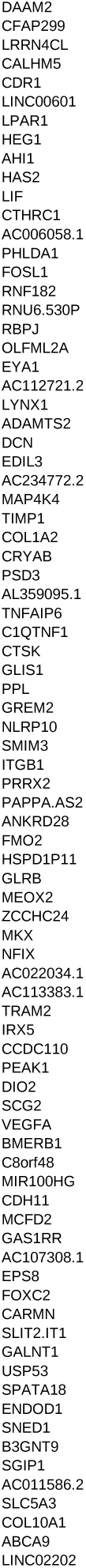
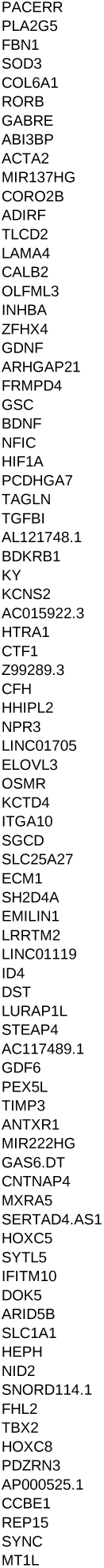
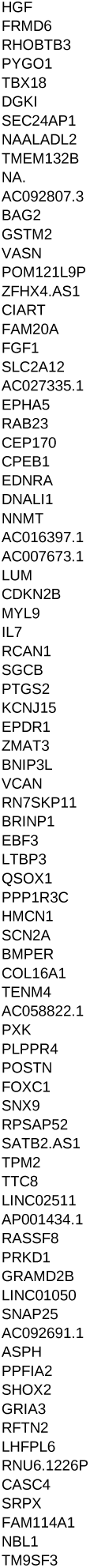
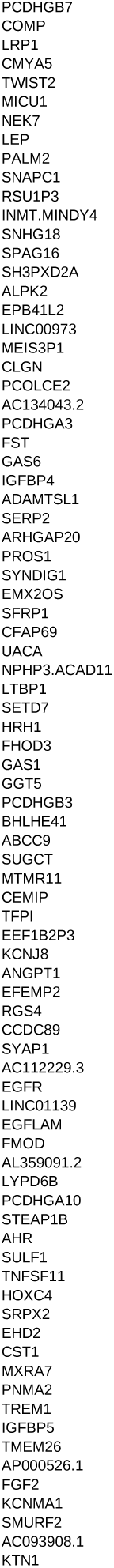
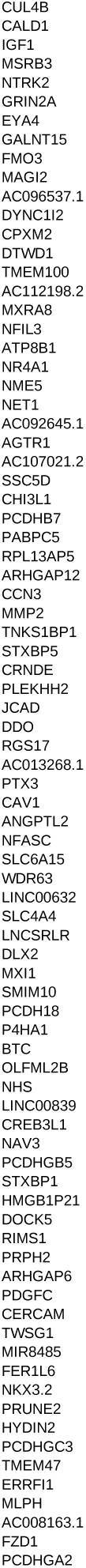
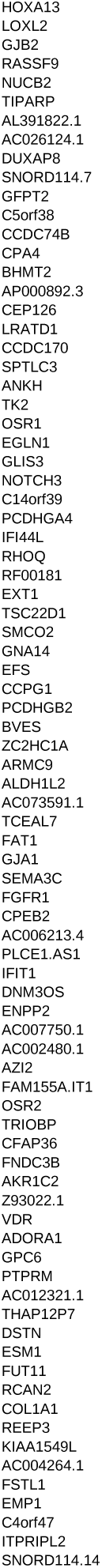
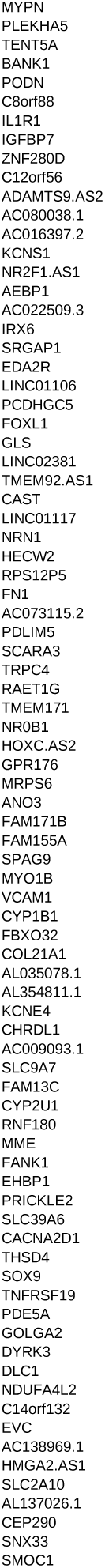
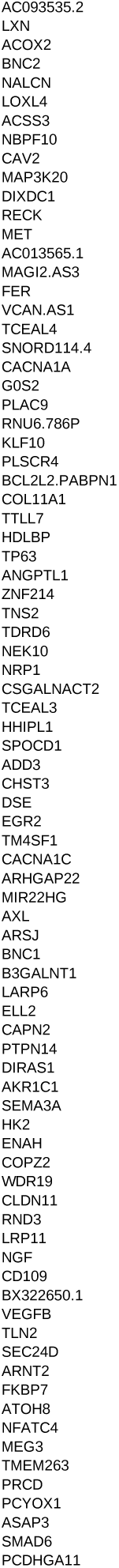
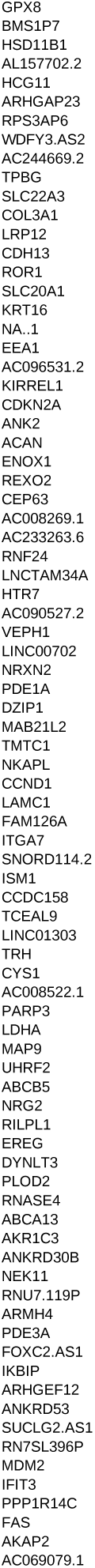
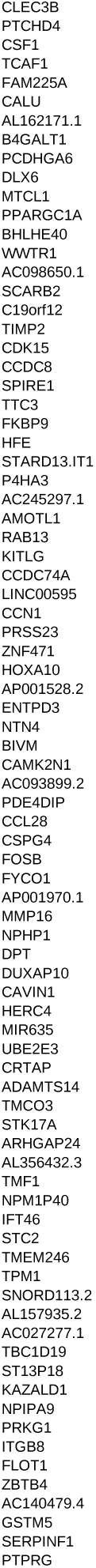
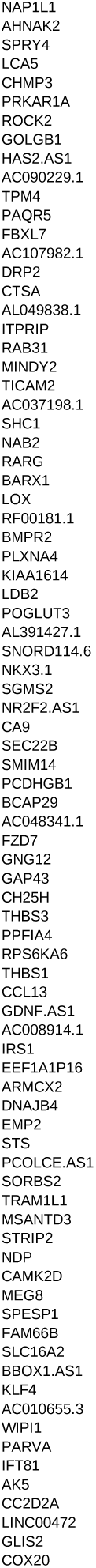
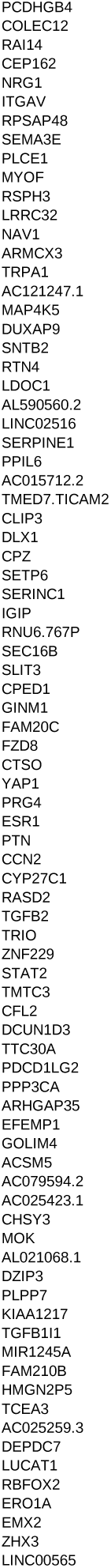
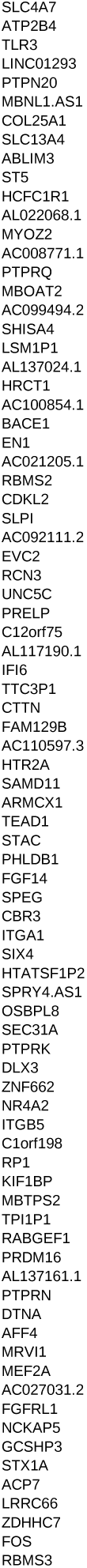
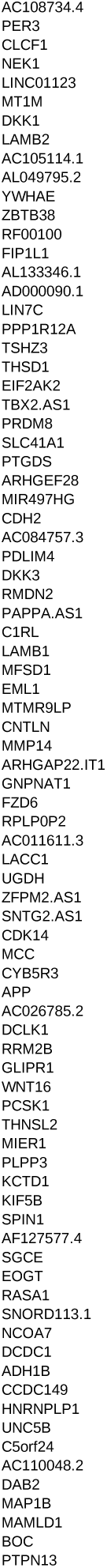
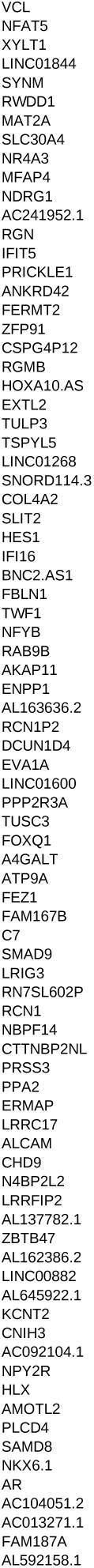
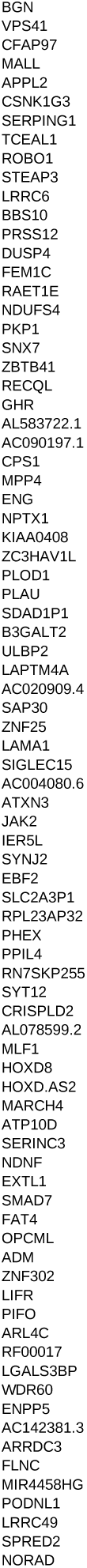
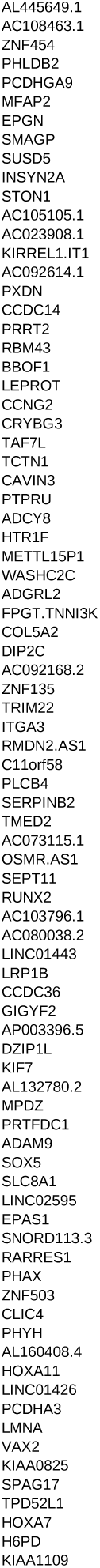
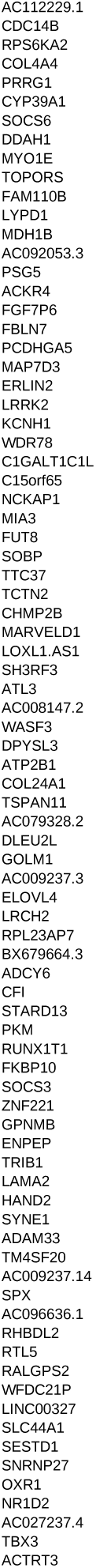
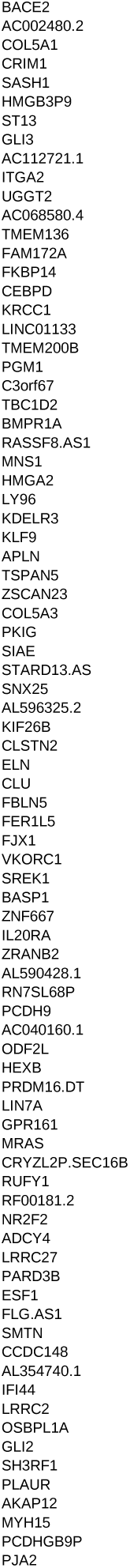
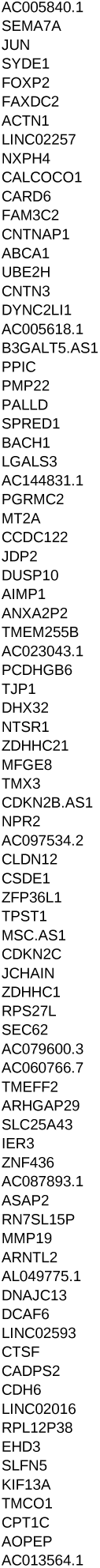
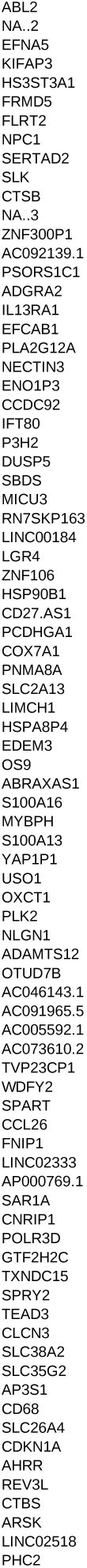
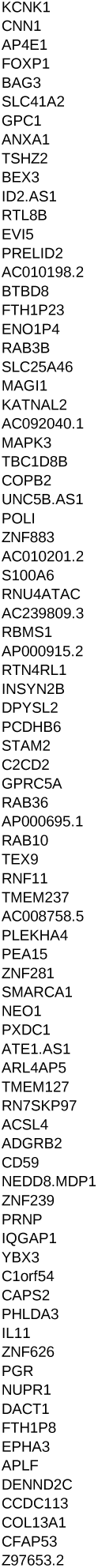
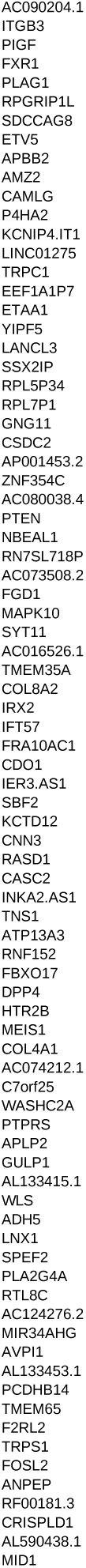
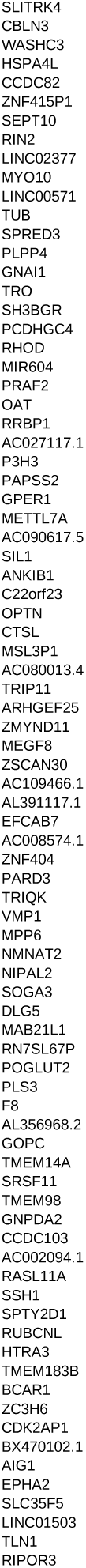
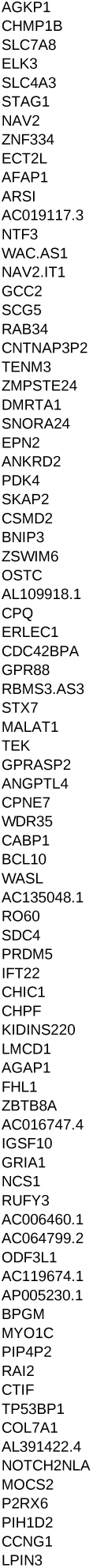
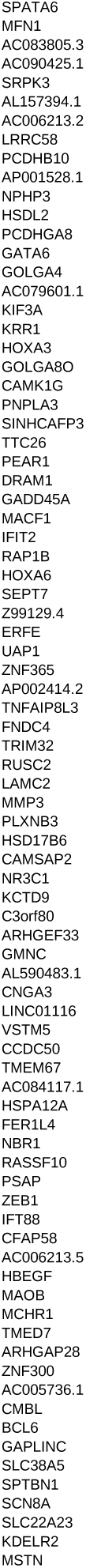
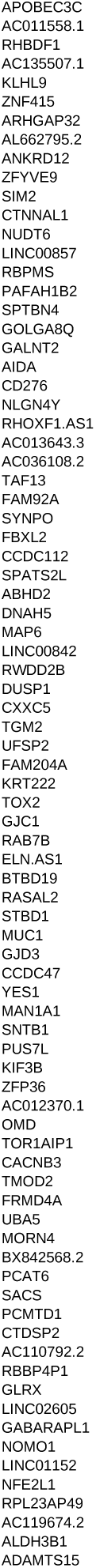
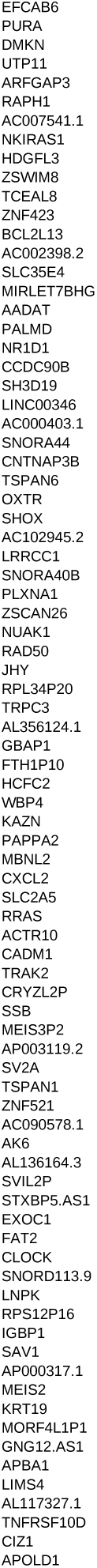
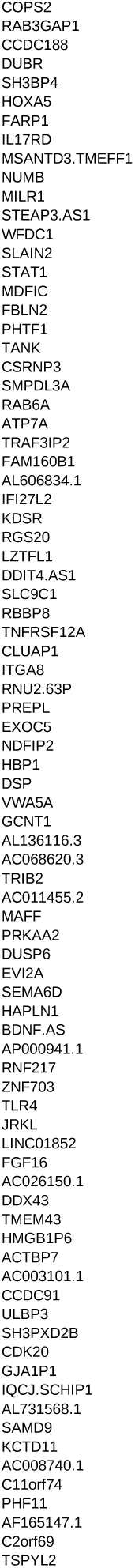
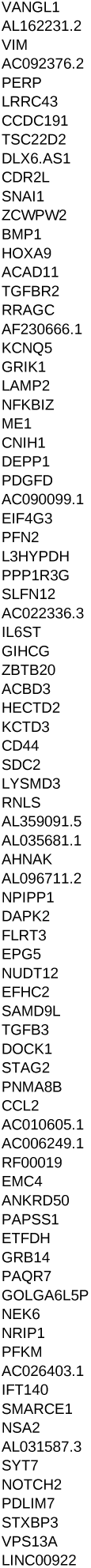
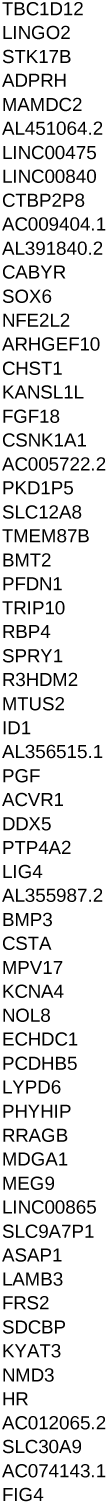
genes names positively enriched in MSC

**Table_S6.**
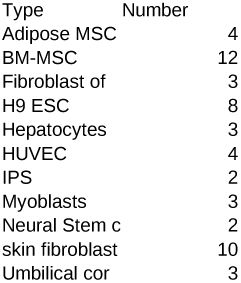
summary of qPCR cohort

**Table_S7.**
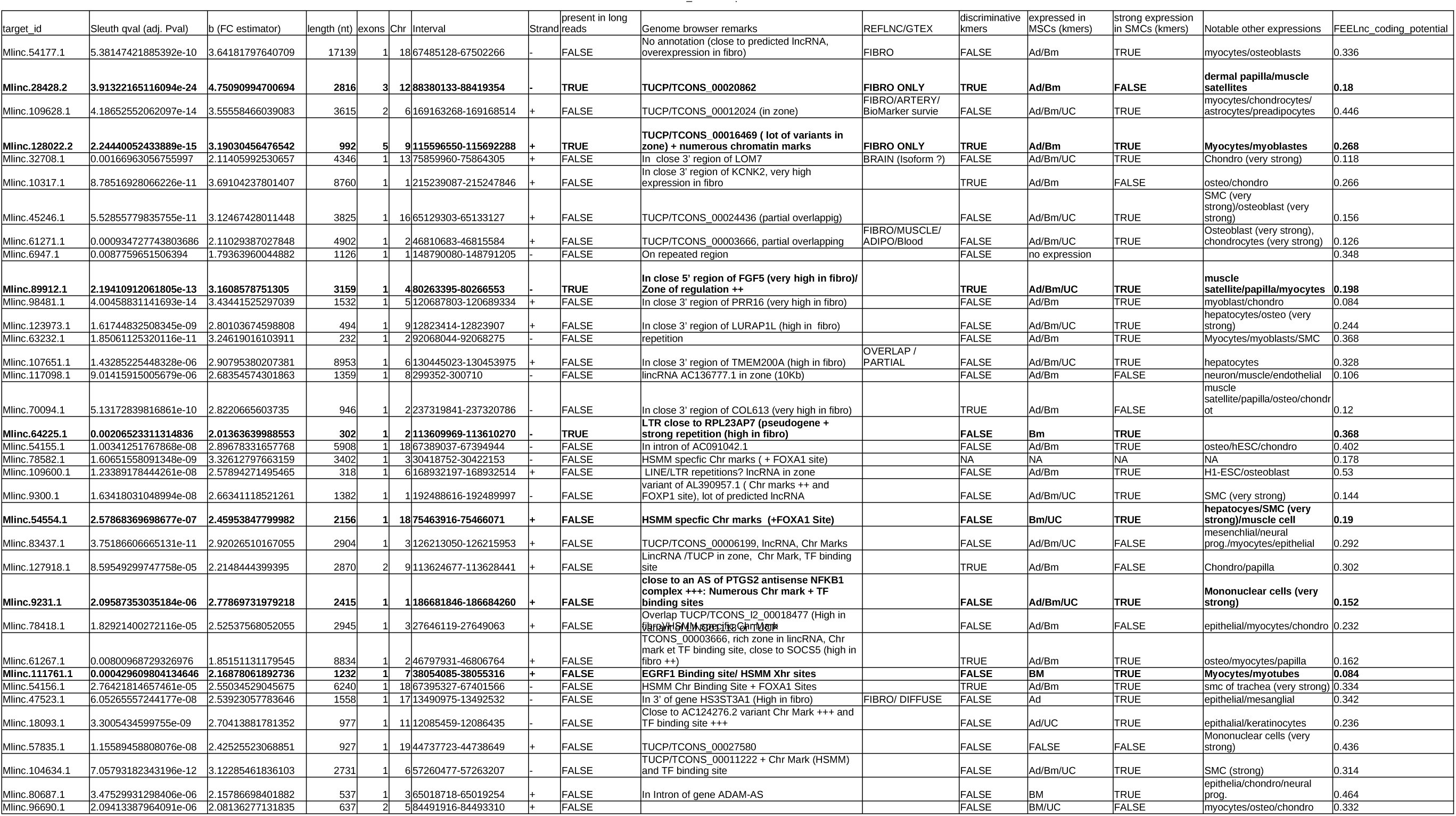
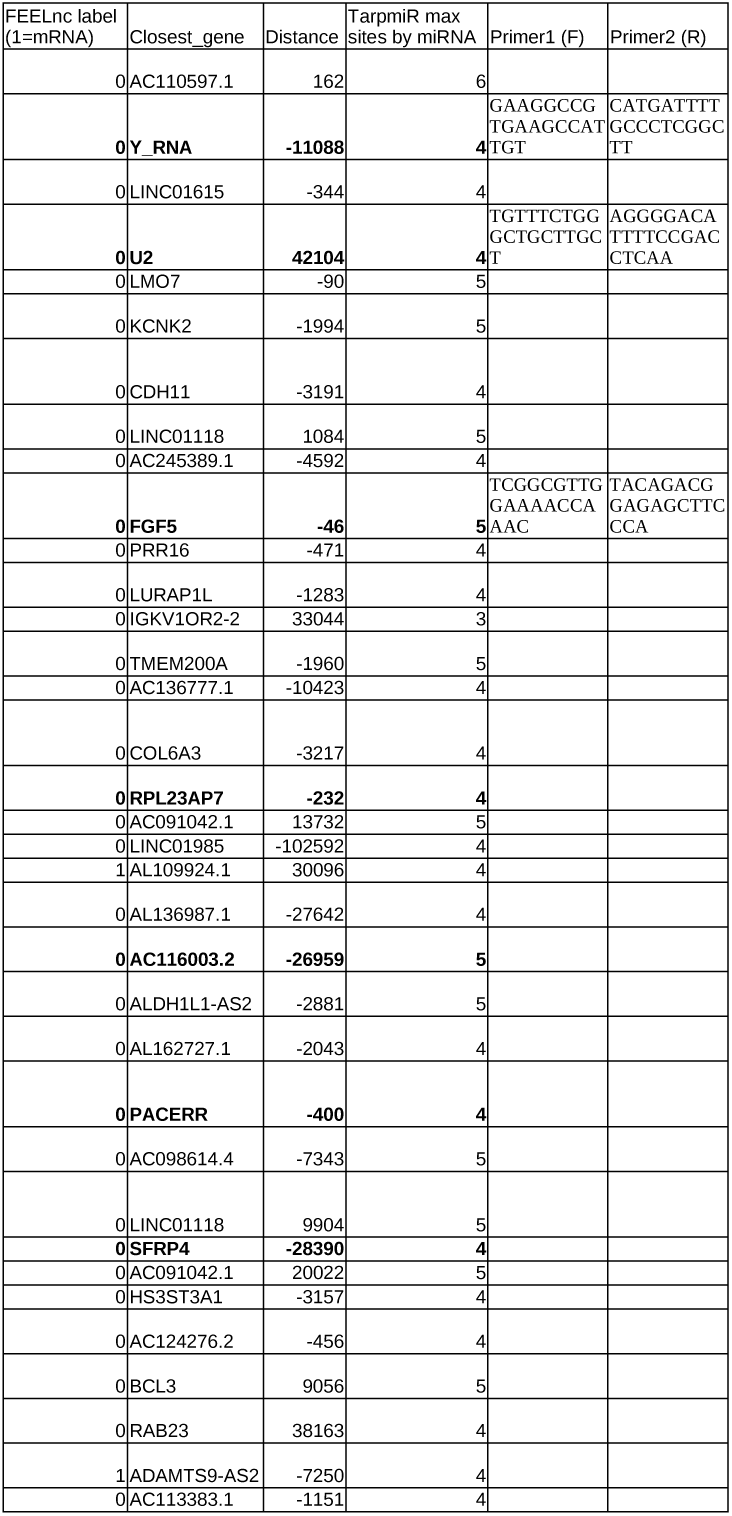
Description of Mlinc Candidates

**Table_S8.**
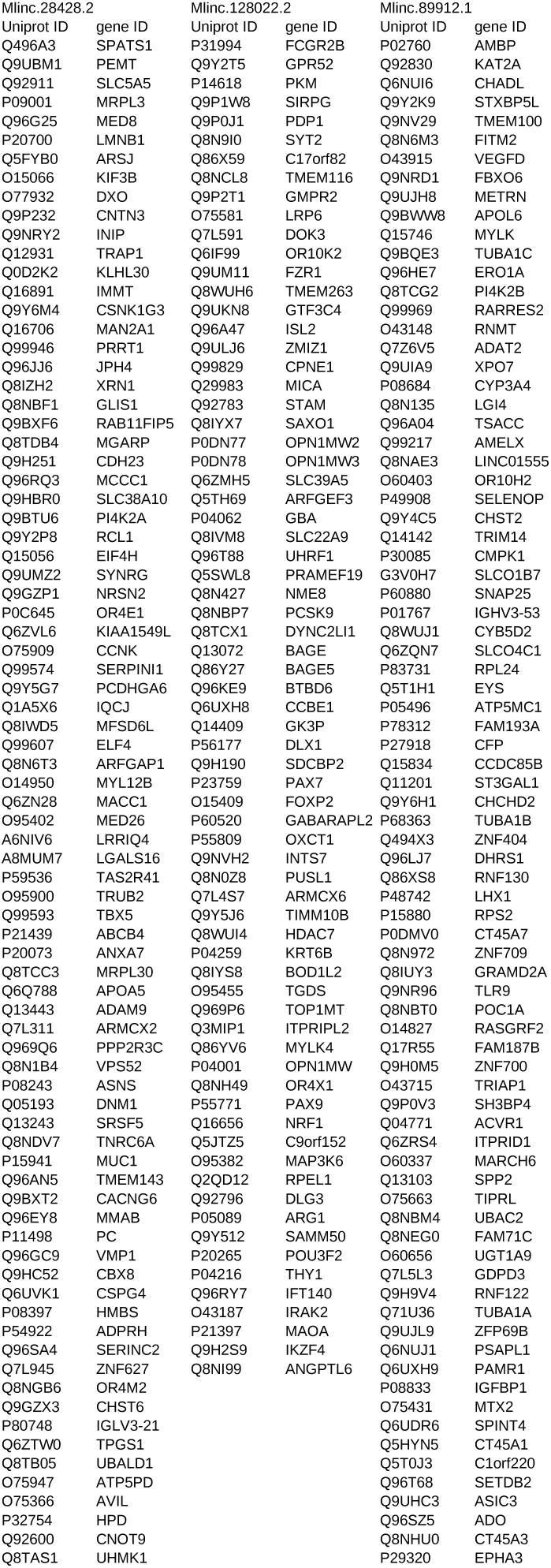
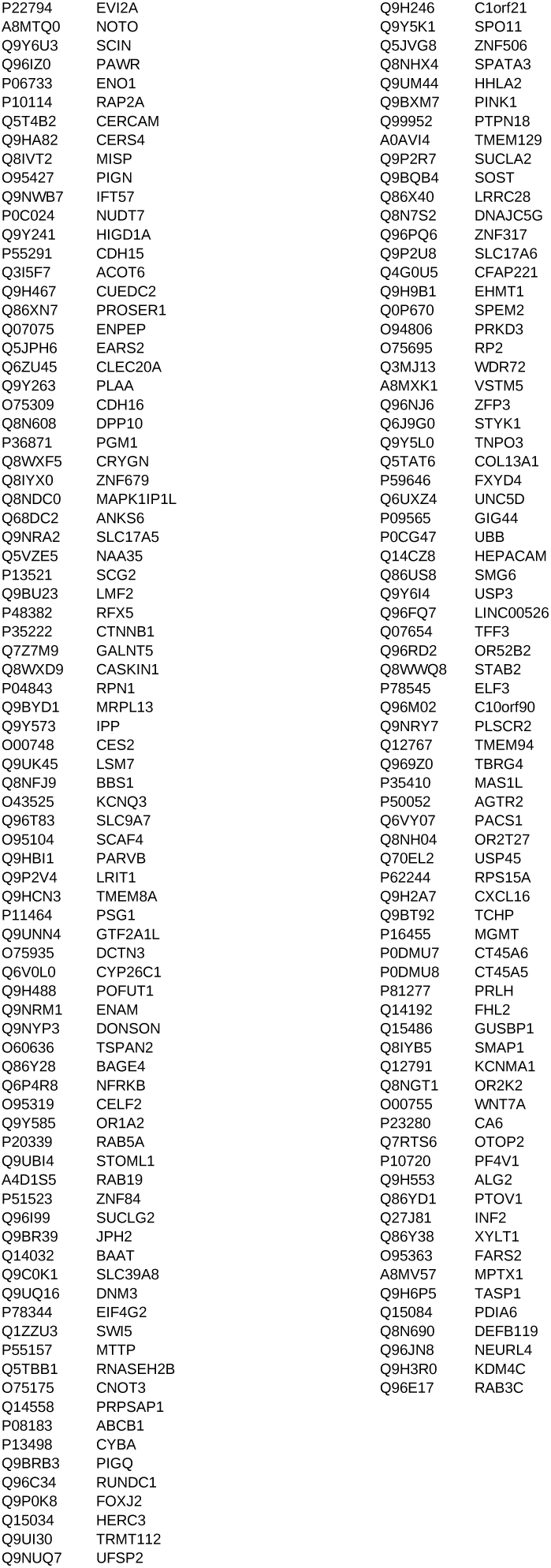
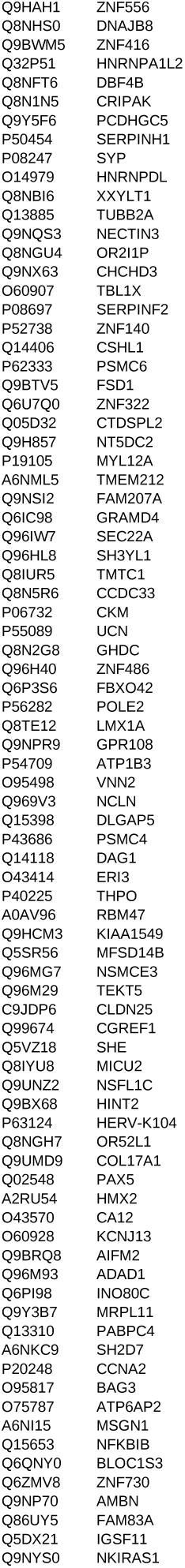
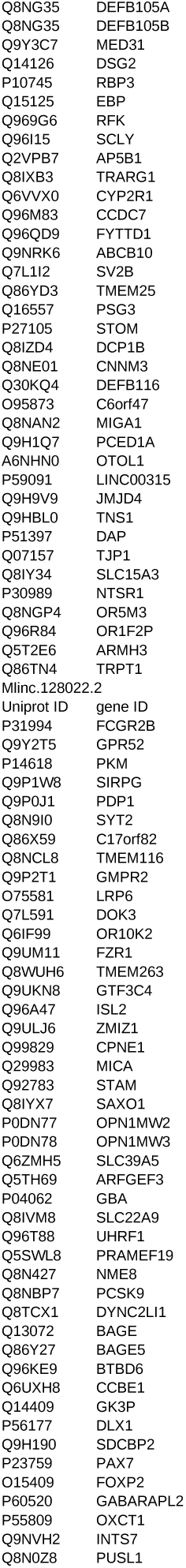
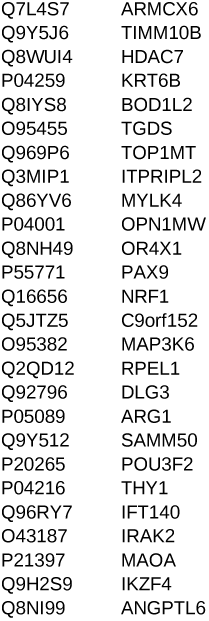
statistically relevant interraction between Mlinc and proteins

## References

1. Gloss, B. S. and Dinger, M. E. (January, 2016) The specificity of long noncoding RNA expression. Biochimica et Biophysica Acta (BBA) - Gene Regulatory Mechanisms, 1859(1), 16–22.

2. Meseure, D., Drak Alsibai, K., Nicolas, A., Bieche, I., and Morillon, A. (2015) Long Noncoding RNAs as New Architects in Cancer Epigenetics, Prognostic Biomarkers, and Potential Therapeutic Targets. BioMed Research International, 2015.

3. Bouckenheimer, J., Assou, S., Riquier, S., Hou, C., Philippe, N., Sansac, C., Lavabre-Bertrand, T., Commes, T., Lematre, J.-M., Boureux, A., and Vos, J. D. (September, 2016) Long non-coding RNAs in human early embryonic development and their potential in ART. Human Reproduction Update,.

4. Li, L. and Chang, H. Y. (October, 2014) Physiological roles of long noncoding RNAs: Insights from knockout mice. Trends in cell biology, 24(10), 594–602.

5. Dhamija, S. and Diederichs, S. (2016) From junk to master regulators of invasion: lncRNA functions in migration, EMT and metastasis. International Journal of Cancer, 139(2), 269– 280.

6. Li, X. and Li, N. (December, 2018) LncRNAs on guard. International Immunopharmacology, 65, 60–63.

7. Morillon, A. and Gautheret, D. (June, 2019) Bridging the gap between reference and real transcriptomes. Genome Biology, 20.

8. Uszczynska-Ratajczak, B., Lagarde, J., Frankish, A., Guig, R., and Johnson, R. (September, 2018) Towards a complete map of the human long non-coding RNA transcriptome. Nature reviews. Genetics, 19(9), 535–548.

9. James, A. R., Schroeder, M. P., Neumann, M., Bastian, L., Eckert, C., Gkbuget, N., Tanchez, J. O., Schlee, C., Isaakidis, K., Schwartz, S., Burmeister, T., von Stackelberg, A., Rieger, M. A., Gllner, S., Horstman, M., Schrappe, M., Kirschner-Schwabe, R., Brggemann, M., Mller-Tidow, C., Serve, H., Akalin, A., and Baldus, C. D. (January, 2019) Long non-coding RNAs defining major subtypes of B cell precursor acute lymphoblastic leukemia. Journal of Hematology & Oncology, 12.

10. Liu, X., Ma, Y., Yin, K., Li, W., Chen, W., Zhang, Y., Zhu, C., Li, T., Han, B., Liu, X., Wang, S., and Zhou, Z. (June, 2019) Long non-coding and coding RNA profiling using strand-specific RNA-seq in human hypertrophic cardiomyopathy. Scientific Data, 6(1), 1–7.

11. Lv, F.-J., Tuan, R. S., Cheung, K. M., and Leung, V. Y. (June, 2014) Concise Review: The Surface Markers and Identity of Human Mesenchymal Stem Cells. STEM CELLS, 32(6), 1408–1419.

12. Soundararajan, M. and Kannan, S. (December, 2018) Fibroblasts and mesenchymal stem cells: Two sides of the same coin?. Journal of Cellular Physiology, 233(12), 9099–9109.

13. Dominici, M., Le Blanc, K., Mueller, I., Slaper-Cortenbach, I., Marini, F., Krause, D., Deans, R., Keating, A., Prockop, D., and Horwitz, E. (2006) Minimal criteria for defining multi-potent mesenchymal stromal cells. The International Society for Cellular Therapy position statement. Cytotherapy, 8(4), 315–317.

14. Fitzsimmons, R. E. B., Mazurek, M. S., Soos, A., and Simmons, C. A. (August, 2018) Mesenchymal Stromal/Stem Cells in Regenerative Medicine and Tissue Engineering. Stem Cells International, 2018.

15. Olsen, T. R., Ng, K. S., Lock, L. T., Ahsan, T., and Rowley, J. A. (June, 2018) Peak MSCAre We There Yet?. Frontiers in Medicine, 5.

16. Tye, C. E., Gordon, J. A. R., Martin-Buley, L. A., Stein, J. L., Lian, J. B., and Stein, G. S. (March, 2015) Could lncRNAs be the missing links in control of mesenchymal stem cell differentiation?. Journal of Cellular Physiology, 230(3), 526–534.

17. Kalwa, M., Hnzelmann, S., Otto, S., Kuo, C.-C., Franzen, J., Joussen, S., Fernandez-Rebollo, E., Rath, B., Koch, C., Hofmann, A., Lee, S.-H., Teschendorff, A. E., Denecke, B., Lin, Q., Widschwendter, M., Weinhold, E., Costa, I. G., and Wagner, W. (December, 2016) The lncRNA HOTAIR impacts on mesenchymal stem cells via triple helix formation. Nucleic Acids Research, 44(22), 10631–10643.

18. Song, W., Gu, W., Qian, Y., Ma, X., Mao, Y., and Liu, W. (2015) Identification of long non-coding RNA involved in osteogenic differentiation from mesenchymal stem cells using RNA-Seq data. Genetics and Molecular Research, 14(4), 18268–18279.

19. Liu, X., Xiang, Q., Xu, F., Huang, J., Yu, N., Zhang, Q., Long, X., and Zhou, Z. (February, 2019) Single-cell RNA-seq of cultured human adipose-derived mesenchymal stem cells. Scientific Data, 6, 190031.

20. Peffers, M. J., Collins, J., Fang, Y., Goljanek-Whysall, K., Rushton, M., Loughlin, J., Proctor, C., and Clegg, P. D. (2016) Age-related changes in mesenchymal stem cells identified using a multi-omics approach. European Cells & Materials, 31, 136–159.

21. Philippe, N., Salson, M., Commes, T., and Rivals, E. (2013) CRAC: an integrated approach to the analysis of RNA-seq reads. Genome Biology, 14, R30.

22. Pertea, M., Pertea, G. M., Antonescu, C. M., Chang, T.-C., Mendell, J. T., and Salzberg, S. L. (March, 2015) StringTie enables improved reconstruction of a transcriptome from RNA-seq reads. Nature Biotechnology, 33(3), 290–295.

23. Quinlan, A. R. and Hall, I. M. (March, 2010) BEDTools: a flexible suite of utilities for comparing genomic features. Bioinformatics, 26(6), 841–842.

24. Ilott, N. E. and Ponting, C. P. (September, 2013) Predicting long non-coding RNAs using RNA sequencing. Methods (San Diego, Calif.), 63(1), 50–59.

25. Li, H. (July, 2016) Minimap and miniasm: fast mapping and de novo assembly for noisy long sequences. Bioinformatics, 32(14), 2103–2110.

26. Bray, N. L., Pimentel, H., Melsted, P., and Pachter, L. (May, 2016) Near-optimal probabilistic RNA-seq quantification. Nature Biotechnology, 34(5), 525–527.

27. Pimentel, H., Bray, N. L., Puente, S., Melsted, P., and Pachter, L. (July, 2017) Differential analysis of RNA-seq incorporating quantification uncertainty. Nature Methods, 14(7), 687– 690.

28. Kursa, M. B., Jankowski, A., and Rudnicki, W. R. (December, 2010) Boruta - A System for Feature Selection. Fundam. Inf., 101(4), 271–285.

29. Agrawal Singh, S., Lerdrup, M., Gomes, A.-L. R., van de Werken, H. J., Vilstrup Johansen, J., Andersson, R., Sandelin, A., Helin, K., and Hansen, K. PLZF targets developmental enhancers for activation during osteogenic differentiation of human mesenchymal stem cells. eLife, 8.

30. Yang, C., Yang, L., Zhou, M., Xie, H., Zhang, C., Wang, M. D., and Zhu, H. (November, 2018) LncADeep: an ab initio lncRNA identification and functional annotation tool based on deep learning. Bioinformatics, 34(22), 3825–3834.

31. Ding, J., Li, X., and Hu, H. (September, 2016) TarPmiR: a new approach for microRNA target site prediction. Bioinformatics, 32(18), 2768–2775.

32. Wucher, V., Legeai, F., Hdan, B., Rizk, G., Lagoutte, L., Leeb, T., Jagannathan, V., Cadieu, E., David, A., Lohi, H., Cirera, S., Fredholm, M., Botherel, N., Leegwater, P. A. J., Le Bguec, C., Fieten, H., Johnson, J., Alfldi, J., Andr, C., Lindblad-Toh, K., Hitte, C., and Derrien, S. (May, 2017) FEELnc: a tool for long non-coding RNA annotation and its application to the dog transcriptome. Nucleic Acids Research, 45(8), e57–e57.

33. Stuart, T., Butler, A., Hoffman, P., Hafemeister, C., Papalexi, E., Mauck, W. M., Hao, Y., Stoeckius, M., Smibert, P., and Satija, R. (June, 2019) Comprehensive Integration of Single-Cell Data. Cell, 177(7), 1888–1902.e21.

34. Butler, A., Hoffman, P., Smibert, P., Papalexi, E., and Satija, R. (May, 2018) Integrating single-cell transcriptomic data across different conditions, technologies, and species. Nature Biotechnology, 36(5), 411–420.

35. Johnson, W. E., Li, C., and Rabinovic, A. (January, 2007) Adjusting batch effects in microarray expression data using empirical Bayes methods. Biostatistics, 8(1), 118–127.

36. Hahne, F. and Ivanek, R. (2016) Visualizing Genomic Data Using Gviz and Bioconductor. In Math, E. and Davis, S., (eds.), Statistical Genomics: Methods and Protocols, Methods in Molecular Biology pp. 335–351 Springer New York New York, NY.

37. Djouad, F., Bony, C., Hupl, T., Uz, G., Lahlou, N., Louis-Plence, P., Apparailly, F., Canovas, F., Rme, T., Sany, J., Jorgensen, C., and Nol, D. (2005) Transcriptional profiles discriminate bone marrow-derived and synovium-derived mesenchymal stem cells. Arthritis Research & Therapy, 7(6), R1304–R1315.

38. Kitzmann, M., Bonnieu, A., Duret, C., Vernus, B., Barro, M., LaoudjChenivesse, D., Verdi, J. M., and Carnac, G. (2006) Inhibition of Notch signaling induces myotube hypertrophy by recruiting a subpopulation of reserve cells. Journal of Cellular Physiology, 208(3), 538–548.

39. Pichard, L., Raulet, E., Fabre, G., Ferrini, J. B., Ourlin, J.-C., and Maurel, P. (2006) Human Hepatocyte Culture. In Phillips, I. R. and Shephard, E. A., (eds.), Cytochrome P450 Protocols, Methods in Molecular Biology pp. 283–293 Humana Press Totowa, NJ.

40. Livak, K. J. and Schmittgen, T. D. (December, 2001) Analysis of Relative Gene Expression Data Using Real-Time Quantitative PCR and the 2CT Method. Methods, 25(4), 402–408.

41. Niazi, F. and Valadkhan, S. (April, 2012) Computational analysis of functional long noncoding RNAs reveals lack of peptide-coding capacity and parallels with 3 UTRs. RNA, 18(4), 825–843.

42. Wang, Y., Xu, T., He, W., Shen, X., Zhao, Q., Bai, J., and You, M. (January, 2018) Genome-wide identification and characterization of putative lncRNAs in the diamondback moth, Plutella xylostella (L.). Genomics, 110(1), 35–42.

43. Cagirici, H. B., Alptekin, B., and Budak, H. (September, 2017) RNA Sequencing and Co-expressed Long Non-coding RNA in Modern and Wild Wheats. Scientific Reports, 7.

44. Salari, R., Aksay, C., Karakoc, E., Unrau, P. J., Hajirasouliha, I., and Sahinalp, S. C. (May, 2009) smyRNA: A Novel Ab Initio ncRNA Gene Finder. PLoS ONE, 4(5).

45. Gu, Q., Tian, H., Zhang, K., Chen, D., Chen, D., Wang, X., and Zhao, J. (2018) Wnt5a/FZD4 Mediates the Mechanical Stretch-Induced Osteogenic Differentiation of Bone Mesenchymal Stem Cells. Cellular Physiology and Biochemistry, 48(1), 215–226.

46. Diederichs, S., Tonnier, V., Mrz, M., Dreher, S. I., Geisbsch, A., and Richter, W. (October, 2019) Regulation of WNT5A and WNT11 during MSC in vitro chondrogenesis: WNT inhibition lowers BMP and hedgehog activity, and reduces hypertrophy. Cellular and molecular life sciences: CMLS, 76(19), 3875–3889.

47. Bermeo, S., Vidal, C., Zhou, H., and Duque, G. (2015) Lamin A/C Acts as an Essential Factor in Mesenchymal Stem Cell Differentiation Through the Regulation of the Dynamics of the Wnt/-Catenin Pathway. Journal of Cellular Biochemistry, 116(10), 2344–2353.

48. Chung, K.-M., Hsu, S.-C., Chu, Y.-R., Lin, M.-Y., Jiaang, W.-T., Chen, R.-H., and Chen, X. (February, 2014) Fibroblast Activation Protein (FAP) Is Essential for the Migration of Bone Marrow Mesenchymal Stem Cells through RhoA Activation. PLoS ONE, 9(2).

49. Ruffl, F., Audoux, J., Boureux, A., Beaumeunier, S., Gaillard, J.-B., Bou Samra, E., Megarbane, A., Cassinat, B., Chomienne, C., Alves, R., Riquier, S., Gilbert, N., Lemaitre, J.-M., Bacq-Daian, D., Boug, A. L., Philippe, N., and Commes, T. (December, 2017) New chimeric RNAs in acute myeloid leukemia. F1000Research, 6, 1302.

50. Krieken, S. E. v. d., Popeijus, H. E., Mensink, R. P., and Plat, J. (2017) Link Between ER-Stress, PPAR-Alpha Activation, and BET Inhibition in Relation to Apolipoprotein A-I Transcription in HepG2 Cells. Journal of Cellular Biochemistry, 118(8), 2161–2167.

51. Delbridge, A. R. D., Kueh, A. J., Ke, F., Zamudio, N. M., El-Saafin, F., Jansz, N., Wang, G.-Y., Iminitoff, M., Beck, T., Haupt, S., Hu, Y., May, R. E., Whitehead, L., Tai, L., Chiang, W., Herold, M. J., Haupt, Y., Smyth, G. K., Thomas, T., Blewitt, M. E., Strasser, A., and Voss, A. K. (April, 2019) Loss of p53 Causes Stochastic Aberrant X-Chromosome Inactivation and Female-Specific Neural Tube Defects. Cell Reports, 27(2), 442–454.e5.

52. Siebringvan Olst, E., Blijlevens, M., de Menezes, R. X., van der MeulenMuileman, I. H., Smit, E. F., and van Beusechem, V. W. (May, 2017) A genomewide siRNA screen for regulators of tumor suppressor p53 activity in human nonsmall cell lung cancer cells identifies components of the RNA splicing machinery as targets for anticancer treatment. Molecular Oncology, 11(5), 534–551.

53. Zhou, Y., Zhong, Y., Wang, Y., Zhang, X., Batista, D. L., Gejman, R., Ansell, P. J., Zhao, J., Weng, C., and Klibanski, A. (August, 2007) Activation of p53 by MEG3 Non-coding RNA. Journal of Biological Chemistry, 282(34), 24731–24742.

54. Uroda, T., Anastasakou, E., Rossi, A., Teulon, J.-M., Pellequer, J.-L., Annibale, P., Pessey, O., Inga, A., Chilln, I., and Marcia, M. (September, 2019) Conserved Pseudoknots in lncRNA MEG3 Are Essential for Stimulation of the p53 Pathway. Molecular Cell, 75(5), 982–995.e9.

55. Haack, T. B., Rolinski, B., Haberberger, B., Zimmermann, F., Schum, J., Strecker, V., Graf, E., Athing, U., Hoppen, T., Wittig, I., Sperl, W., Freisinger, P., Mayr, J. A., Strom, T. M., Meitinger, T., and Prokisch, H. (2013) Homozygous missense mutation in BOLA3 causes multiple mitochondrial dysfunctions syndrome in two siblings. Journal of Inherited Metabolic Disease, 36(1), 55–62.

56. Yu Qiujun, Tai Yi-Yin, Tang Ying, Zhao Jingsi, Negi Vinny, Culley Miranda K., Pilli Jyotsna, Sun Wei, Brugger Karin, Mayr Johannes, Saggar Rajeev, Saggar Rajan, Wallace W. Dean, Ross David J., Waxman Aaron B., Wendell Stacy G., Mullett Steven J., Sembrat John, Rojas Mauricio, Khan Omar F., Dahlman James E., Sugahara Masataka, Kagiyama Nobuyuki, Satoh Taijyu, Zhang Manling, Feng Ning, Gorcsan John, Vargas Sara O., Haley Kathleen J., Kumar Rahul, Graham Brian B., Langer Robert, Anderson Daniel G., Wang Bing, Shiva Sruti, Bertero Thomas, and Chan Stephen Y. (May, 2019) BOLA (BolA Family Member 3) Deficiency Controls Endothelial Metabolism and Glycine Homeostasis in Pulmonary Hypertension. Circulation, 139(19), 2238–2255.

57. Wang, J. and Li, K. (April, 2018) AB042. P013. LncRNAPTCHD3P1 enhances chemosensitivity of gemcitabine in pancreatic cancer and inhibits cancer cell proliferation and metastasis via inhibiting Warburg effect. Annals of Pancreatic Cancer, 1(4).

58. Qin, L., Wang, M., Zuo, J., Feng, X., Liang, X., Wu, Z., and Ye, H. (April, 2015) Cytosolic BolA Plays a Repressive Role in the Tolerance against Excess Iron and MV-Induced Oxidative Stress in Plants. PLoS ONE, 10(4).

59. Kitajima, S., Asahina, H., Chen, T., Guo, S., Quiceno, L. G., Cavanaugh, J. D., Merlino, A. A., Tange, S., Terai, H., Kim, J. W., Wang, X., Zhou, S., Xu, M., Wang, S., Zhu, Z., Thai, T. C., Takahashi, C., Wang, Y., Neve, R., Stinson, S., Tamayo, P., Watanabe, H., Kirschmeier, P. T., Wong, K.-K., and Barbie, D. A. (September, 2018) Overcoming Resistance to Dual Innate Immune and MEK Inhibition Downstream of KRAS. Cancer cell, 34(3), 439–452.e6.

60. Raj, N. and Bam, R. (August, 2019) Reciprocal Crosstalk Between YAP1/Hippo Pathway and the p53 Family Proteins: Mechanisms and Outcomes in Cancer. Frontiers in Cell and Developmental Biology, 7.

61. He, J., Tu, C., and Liu, Y. (2018) Role of lncRNAs in aging and age-related diseases. AGING MEDICINE, 1(2), 158–175.

62. Schuff, M., Rssner, A., Wacker, S. A., Donow, C., Gessert, S., and Knchel, W. (2007) FoxN3 is required for craniofacial and eye development of Xenopus laevis. Developmental Dynamics, 236(1), 226–239.

63. Samaan, G., Yugo, D., Rajagopalan, S., Wall, J., Donnell, R., Goldowitz, D., Gopalakrishnan, R., and Venkatachalam, S. (September, 2010) Foxn3 is essential for craniofacial development in mice and a putative candidate involved in human congenital craniofacial defects. Biochemical and Biophysical Research Communications, 400(1), 60–65.

64. Brum, A. M., van de Peppel, J., van der Leije, C. S., Schreuders-Koedam, M., Eijken, M., van der Eerden, B. C. J., and van Leeuwen, J. P. T. M. (October, 2015) Connectivity Mapbased discovery of parbendazole reveals targetable human osteogenic pathway. Proceedings of the National Academy of Sciences of the United States of America, 112(41), 12711–12716.

65. del Real, A., Prez-Campo, F. M., Fernndez, A. F., Saudo, C., Ibarbia, C. G., Prez-Nez, M. I., Criekinge, W. V., Braspenning, M., Alonso, M. A., Fraga, M. F., and Riancho, J. A. (December, 2016) Differential analysis of genome-wide methylation and gene expression in mesenchymal stem cells of patients with fractures and osteoarthritis. Epigenetics, 12(2), 113–122.

66. Bai, J., Yao, B., Wang, L., Sun, L., Chen, T., Liu, R., Yin, G., Xu, Q., and Yang, W. (2019) lncRNA A1BG-AS1 suppresses proliferation and invasion of hepatocellular carcinoma cells by targeting miR-216a-5p. Journal of Cellular Biochemistry, 120(6), 10310–10322.

67. Li, N., Lee, W. Y.-W., Lin, S.-E., Ni, M., Zhang, T., Huang, X.-R., Lan, H.-Y., and Li, G. (October, 2014) Partial loss of Smad7 function impairs bone remodeling, osteogenesis and enhances osteoclastogenesis in mice. Bone, 67, 46–55.

68. Vishal, M., Vimalraj, S., Ajeetha, R., Gokulnath, M., Keerthana, R., He, Z., Partridge, M. C., and Selvamurugan, N. (2017) MicroRNA-590-5p Stabilizes Runx2 by Targeting Smad7 During Osteoblast Differentiation. Journal of Cellular Physiology, 232(2), 371–380.

69. Nowak, W. N., Taha, H., Kachamakova-Trojanowska, N., Stpniewski, J., Markiewicz, J. A., Kusienicka, A., Szade, K., Szade, A., Bukowska-Strakova, K., Hajduk, K., Klska, D., Kopacz, A., Grochot-Przczek, A., Barthenheier, K., Cauvin, C., Dulak, J., and Jzkowicz, A. (October, 2017) Murine Bone Marrow Mesenchymal Stromal Cells Respond Efficiently to Oxidative Stress Despite the Low Level of Heme Oxygenases 1 and 2. Antioxidants & Redox Signaling, 29(2), 111–127.

70. Balogh, E., Paragh, G., and Jeney, V. (October, 2018) Influence of Iron on Bone Homeostasis. Pharmaceuticals, 11(4).

71. Puri, N., Sodhi, K., Haarstad, M., Kim, D. H., Bohinc, S., Foglio, E., Favero, G., and Abraham, N. G. (June, 2012) Heme Induced Oxidative Stress Attenuates Sirtuin1 and Enhances Adipogenesis in Mesenchymal Stem Cells and Mouse Pre-Adipocytes. Journal of Cellular Biochemistry, 113(6), 1926–1935.

72. Luo, Y., Tao, H., Jin, L., Xiang, W., and Guo, W. (November, 2019) CDKN2B-AS1 Exerts Oncogenic Role in Osteosarcoma by Promoting Cell Proliferation and Epithelial to Mesenchymal Transition. Cancer Biotherapy and Radiopharmaceuticals,.

73. Congrains, A., Kamide, K., Ohishi, M., and Rakugi, H. (January, 2013) ANRIL: Molecular Mechanisms and Implications in Human Health. International Journal of Molecular Sciences, 14(1), 1278–1292.

74. Yin, Z., Ding, H., He, E., Chen, J., and Li, M. (October, 2016) Overexpression of long non-coding RNA MFI2 promotes cell proliferation and suppresses apoptosis in human osteosarcoma. Oncology Reports, 36(4), 2033–2040.

75. Li, C., Tan, F., Pei, Q., Zhou, Z., Zhou, Y., Zhang, L., Wang, D., and Pei, H. (2019) Noncoding RNA MFI2-AS1 promotes colorectal cancer cell proliferation, migration and invasion through miR-574-5p/MYCBP axis. Cell Proliferation, 52(4), e12632.

76. Zhu, C., Huang, L., Xu, F., Li, P., Li, P., and Hu, F. (October, 2019) LncRNA PCAT6 promotes tumor progression in osteosarcoma via activation of TGF-pathway by sponging miR-185-5p. Biochemical and Biophysical Research Communications,.

77. Dong, P., Xiong, Y., Yue, J., Hanley, S. J. B., Kobayashi, N., Todo, Y., and Watari, H. (October, 2018) Long Non-coding RNA NEAT1: A Novel Target for Diagnosis and Therapy in Human Tumors. Frontiers in Genetics, 9.

78. Ahmed, A. S. I., Dong, K., Liu, J., Wen, T., Yu, L., Xu, F., Kang, X., Osman, I., Hu, G., Bunting, K. M., Crethers, D., Gao, H., Zhang, W., Liu, Y., Wen, K., Agarwal, G., Hirose, T., Nakagawa, S., Vazdarjanova, A., and Zhou, J. (September, 2018) Long noncoding RNA NEAT1 (nuclear paraspeckle assembly transcript 1) is critical for phenotypic switching of vascular smooth muscle cells. Proceedings of the National Academy of Sciences, 115(37), E8660–E8667.

79. Taiana, E., Favasuli, V., Ronchetti, D., Todoerti, K., Pelizzoni, F., Manzoni, M., Barbieri, M., Fabris, S., Silvestris, I., Cantafio, M. E. G., Platonova, N., Zuccal, V., Maltese, L., Soncini, D., Ruberti, S., Cea, M., Chiaramonte, R., Amodio, N., Tassone, P., Agnelli, L., and Neri, A. (August, 2019) Long non-coding RNA NEAT1 targeting impairs the DNA repair machinery and triggers anti-tumor activity in multiple myeloma. Leukemia, pp. 1–11.

80. Wan, G., Mathur, R., Hu, X., Liu, Y., Zhang, X., Peng, G., and Lu, X. (May, 2013) Long non-coding RNA ANRIL (CDKN2B-AS) is induced by the ATM-E2F1 signaling pathway. Cellular signalling, 25(5), 1086–1095.

81. Ding, K., Liao, Y., Gong, D., Zhao, X., and Ji, W. (July, 2018) Effect of long non-coding RNA H19 on oxidative stress and chemotherapy resistance of CD133+ cancer stem cells via the MAPK/ERK signaling pathway in hepatocellular carcinoma. Biochemical and Biophysical Research Communications, 502(2), 194–201.

82. Yu, J.-L., Li, C., Che, L.-H., Zhao, Y.-H., and Guo, Y.-B. (2019) Downregulation of long noncoding RNA H19 rescues hippocampal neurons from apoptosis and oxidative stress by inhibiting IGF2 methylation in mice with streptozotocin-induced diabetes mellitus. Journal of Cellular Physiology, 234(7), 10655–10670.

83. Hazell, G. G. J., Peachey, A. M. G., Teasdale, J. E., Sala-Newby, G. B., Angelini, G. D., Newby, A. C., and White, S. J. (December, 2016) PI16 is a shear stress and inflammation-regulated inhibitor of MMP2. Scientific Reports, 6.

84. Puvvula, P. K. (May, 2019) LncRNAs Regulatory Networks in Cellular Senescence. International Journal of Molecular Sciences, 20(11).

85. Spanner, M., Weber, K., Lanske, B., Ihbe, A., Siggelkow, H., Schtze, H., and Atkinson, M. J. (August, 1995) The iron-binding protein ferritin is expressed in cells of the osteoblastic lineage in vitro and in vivo. Bone, 17(2), 161–165.

86. Balogh, E., Tolnai, E., Nagy, B., Nagy, B., Balla, G., Balla, J., and Jeney, V. (September, 2016) Iron overload inhibits osteogenic commitment and differentiation of mesenchymal stem cells via the induction of ferritin. Biochimica et Biophysica Acta (BBA) - Molecular Basis of Disease, 1862(9), 1640–1649.

87. Zarjou, A., Jeney, V., Arosio, P., Poli, M., Antal-Szalms, P., Agarwal, A., Balla, G., and Balla, J. (June, 2009) Ferritin Prevents Calcification and Osteoblastic Differentiation of Vascular Smooth Muscle Cells. Journal of the American Society of Nephrology, 20(6), 1254– 1263.

88. Doi, M., Nagano, A., and Nakamura, Y. (January, 2002) Genome-wide Screening by cDNA Microarray of Genes Associated with Matrix Mineralization by Human Mesenchymal Stem Cells in Vitro. Biochemical and Biophysical Research Communications, 290(1), 381–390.

89. Liu, Z., Zheng, Z., Qi, J., Wang, J., Zhou, Q., Hu, F., Liang, J., Li, C., Zhang, W., and Zhang, X. (December, 2018) CD24 identifies nucleus pulposus progenitors/notochordal cells for disc regeneration. Journal of Biological Engineering, 12(1), 35.

90. Tsai, Y.-H., Lin, K.-L., Huang, Y.-P., Hsu, Y.-C., Chen, C.-H., Chen, Y., Sie, M.-H., Wang, G.-J., and Lee, M.-J. (2015) Suppression of ornithine decarboxylase promotes osteogenic differentiation of human bone marrow-derived mesenchymal stem cells. FEBS Letters, 589(16), 2058–2065.

91. Chang, C.-F., Hsu, K.-H., Shen, C.-N., Li, C.-L., and Lu, J. (2014) Enrichment and Characterization of Two Subgroups of Committed Osteogenic Cells in the Mouse Endosteal Bone Marrow with Expression Levels of CD24. Journal of Bone Research, 2(2), 1–9.

92. Park, G. C., Song, J. S., Park, H.-Y., Shin, S.-C., Jang, J. Y., Lee, J.-C., Wang, S.-G., Lee, B.-J., and Jung, J.-S. (May, 2016) Role of Fibroblast Growth Factor-5 on the Proliferation of Human Tonsil-Derived Mesenchymal Stem Cells. Stem Cells and Development, 25(15), 1149–1160.

93. Kornmann, M., Ishiwata, T., Beger, H. G., and Korc, M. (September, 1997) Fibroblast growth factor-5 stimulates mitogenic signaling and is overexpressed in human pancreatic cancer: evidence for autocrine and paracrine actions. Oncogene, 15(12), 1417–1424.

94. Williamson, E. A., Wray, J. W., Bansal, P., and Hromas, R. (2012) Overview for the Histone Codes for DNA Repair. Progress in molecular biology and translational science, 110, 207– 227.

95. Wang, S., Hu, B., Ding, Z., Dang, Y., Wu, J., Li, D., Liu, X., Xiao, B., Zhang, W., Ren, R., Lei, J., Hu, H., Chen, C., Chan, P., Li, D., Qu, J., Tang, F., and Liu, G.-H. (January, 2018) ATF6 safeguards organelle homeostasis and cellular aging in human mesenchymal stem cells. Cell Discovery, 4.

96. Fu, L., Hu, Y., Song, M., Liu, Z., Zhang, W., Yu, F.-X., Wu, J., Wang, S., Izpisua Belmonte, J. C., Chan, P., Qu, J., Tang, F., and Liu, G.-H. (April, 2019) Up-regulation of FOXD1 by YAP alleviates senescence and osteoarthritis. PLoS Biology, 17(4).

97. Samsonraj, R. M., Dudakovic, A., Manzar, B., Sen, B., Dietz, A. B., Cool, S. M., Rubin, J., and van Wijnen, A. J. (December, 2017) Osteogenic Stimulation of Human AdiposeDerived Mesenchymal Stem Cells Using a Fungal Metabolite That Suppresses the Polycomb Group Protein EZH2. Stem Cells Translational Medicine, 7(2), 197–209.

98. Dudakovic, A., Gluscevic, M., Paradise, C. R., Dudakovic, H., Khani, F., Thaler, R., Ahmed, F. S., Li, X., Dietz, A. B., Stein, G. S., Montecino, M. A., Deyle, D. R., Westendorf, J. J., and van Wijnen, A. J. (April, 2017) Profiling of human epigenetic regulators using a semi-automated real-time qPCR platform validated by next generation sequencing. Gene, 609, 28–37.

99. Camilleri, E. T., Gustafson, M. P., Dudakovic, A., Riester, S. M., Garces, C. G., Paradise, C. R., Takai, H., Karperien, M., Cool, S., Sampen, H.-J. I., Larson, A. N., Qu, W., Smith, J., Dietz, A. B., and van Wijnen, A. J. (August, 2016) Identification and validation of multiple cell surface markers of clinical-grade adipose-derived mesenchymal stromal cells as novel release criteria for good manufacturing practice-compliant production. Stem Cell Research & Therapy, 7.

100. Knight, C., James, S., Kuntin, D., Fox, J., Newling, K., Hollings, S., Pennock, R., and Genever, P. (January, 2019) Epidermal growth factor can signal via -catenin to control proliferation of mesenchymal stem cells independently of canonical Wnt signalling. Cellular Signalling, 53, 256–268.

101. Jiang, S., Cheng, S.-J., Ren, L.-C., Wang, Q., Kang, Y.-J., Ding, Y., Hou, M., Yang, X.-X., Lin, Y., Liang, N., and Gao, G. (September, 2019) An expanded landscape of human long noncoding RNA. Nucleic Acids Research, 47(15), 7842–7856.

102. Chang, T.-H., Huang, H.-D., Ong, W.-K., Fu, Y.-J., Lee, O. K., Chien, S., and Ho, J. H. (April, 2014) The effects of actin cytoskeleton perturbation on keratin intermediate filament formation in mesenchymal stem/stromal cells. Biomaterials, 35(13), 3934–3944.

103. Chang, Y., Li, H., and Guo, Z. (2014) Mesenchymal stem cell-like properties in fibroblasts. Cellular Physiology and Biochemistry: International Journal of Experimental Cellular Physiology, Biochemistry, and Pharmacology, 34(3), 703–714.

104. Denu, R. A., Nemcek, S., Bloom, D. D., Goodrich, A. D., Kim, J., Mosher, D. F., and Hematti, P. (August, 2016) Fibroblasts and Mesenchymal Stromal/Stem Cells Are Phenotypically Indistinguishable. Acta Haematologica, 136(2), 85–97.

105. Ball, S. G., Shuttleworth, A. C., and Kielty, C. M. (April, 2004) Direct cell contact influences bone marrow mesenchymal stem cell fate. The International Journal of Biochemistry & Cell Biology, 36(4), 714–727.

106. Tamama, K., Sen, C. K., and Wells, A. (October, 2008) Differentiation of Bone Marrow Mesenchymal Stem Cells into the Smooth Muscle Lineage by Blocking ERK/MAPK Signaling Pathway. Stem Cells and Development, 17(5), 897–908.

107. Kumar, A., DSouza, S. S., Moskvin, O. V., Toh, H., Wang, B., Zhang, J., Swanson, S., Guo, L.-W., Thomson, J. A., and Slukvin, I. I. (May, 2017) Specification and Diversification of Pericytes and Smooth Muscle Cells from Mesenchymoangioblasts. Cell Reports, 19(9), 1902–1916.

